# Structure-Guided Discovery of High-Affinity Methotrexate Aptamers for Electrochemical Drug Monitoring

**DOI:** 10.64898/2026.10.06.757117

**Authors:** Micaela B. Sequera, Thiago C. Pimenta, Rafael Vieira, Alexandr Beliaev, Kirill Alexandrov, Sergio F. Sousa, Saimon M. Silva, Elena Eremeeva

**Affiliations:** Centre for Agriculture and the Bioeconomy, School of Biology and Environmental Science, Queensland University of Technology, Gardens Point Campus, Brisbane, QLD, 4000, Australia; The Biomedical and Environmental Sensor Technology (BEST) Research Centre, Biosensors Program, La Trobe Institute for Molecular Science (LIMS), La Trobe University, Bundoora, VIC, 3086, Australia; Department of Biochemistry and Chemistry, School of Agriculture, Biomedicine and Environment, La Trobe University, Bundoora, VIC, 3086, Australia; LAQV/REQUIMTE, BioSIM, Department of Biomedicine, Faculty of Medicine, University of Porto, Rua Alameda Prof. Hernâni Monteiro, Porto, 4200-319, Portugal; ARC Centre of Excellence in Synthetic Biology, Canberra, Australian Capital Territory, Australia; Pacific Northwest National Laboratory Richland, WA 99354, United States; CERES, BIA Lab, Department of Chemical Engineering, FCTUC, University of Coimbra, Rua Sílvio Lima, Pólo II – Pinhal de Marrocos, 3030-790 Coimbra, Portugal; ARC Research Hub for Molecular Biosensors at Point-of-Use (MOBIUS), La Trobe University, Melbourne, VIC 3086, Australia; Aptamer Facility, Synthetic Biology Program, La Trobe Institute for Molecular Science (LIMS), La Trobe University, Bundoora, VIC, 3086, Australia

## Abstract

Methotrexate (MTX) is a narrow-therapeutic-index drug widely used in cancer and autoimmune disease treatment, for which delayed clearance can cause severe toxicity and requires close therapeutic drug monitoring. Here, we integrate high-stringency Capture-SELEX, biophysical characterization, molecular modeling, mutational validation, and electrochemical transduction to develop two structurally distinct MTX-binding DNA aptamer families. Full-length 3315 and 930 bound MTX with apparent K_D_ values of 89 and 408 nM, respectively. Systematic truncation generated compact functional variants: 3315-55 retained 63 nM affinity and broad buffer tolerance, whereas 930-65 reached an apparent K_D_ of 44 nM in 10 mM Mg^2+^, revealing markedly different ion dependencies. Biolayer interferometry confirmed rapid and reversible MTX recognition, while 3315 variants retained binding in serum-containing buffer. Computational analysis and site-directed mutagenesis identified distinct recognition architectures: a compact G-rich scaffold coupled to a central binding interface in 3315 and a localized Mg^2+^-stabilized binding pocket in 930. Methylene blue-labeled 3315-55 and 930-65 were integrated into electrochemical aptamer-based sensors, producing concentration-dependent responses to 100-1000 uM MTX in PBS, discriminating MTX from folic acid, and responding to MTX directly in undiluted human blood. Together, these results establish a structure-guided workflow linking aptamer discovery, mechanistic validation, and electrochemical translation, providing new molecular recognition elements and a foundation for future reagent-less MTX therapeutic drug monitoring.

**GRAPHICAL ABSTRACT:** 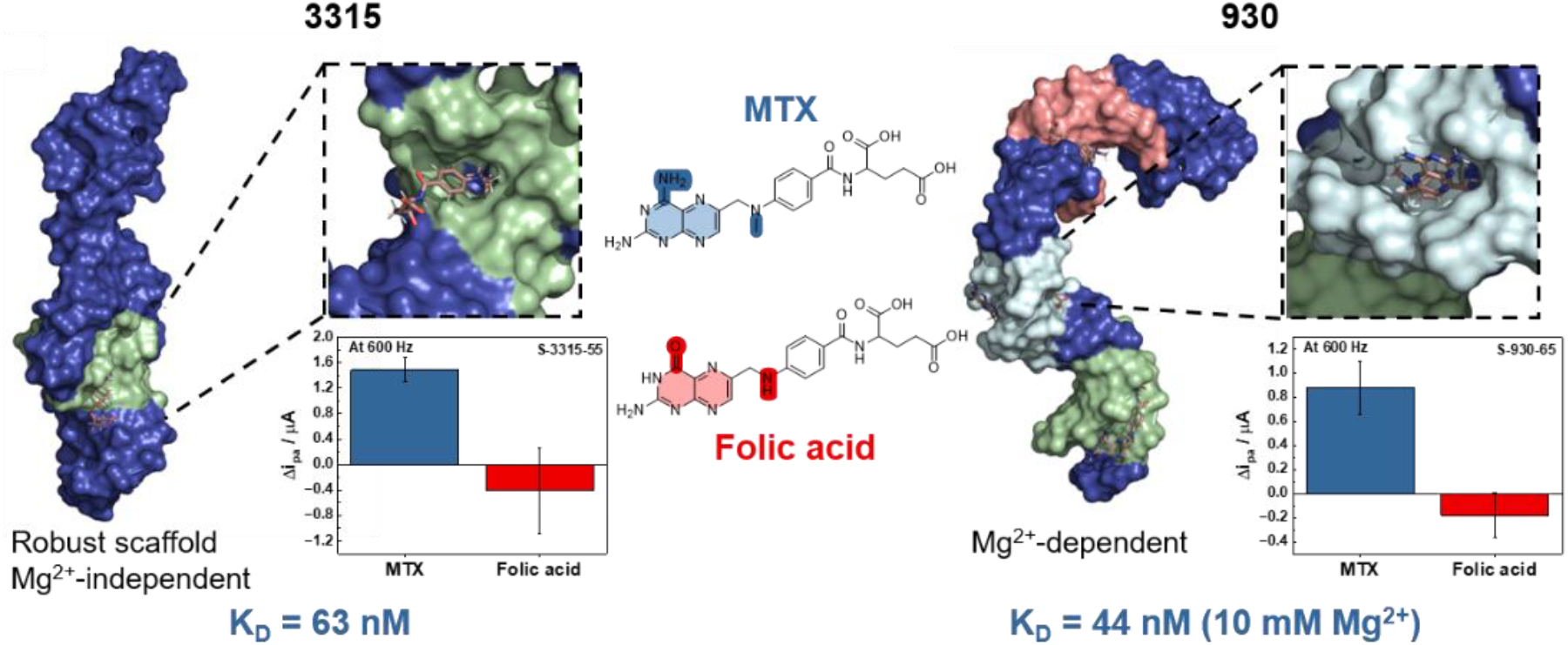

## INTRODUCTION

Methotrexate (MTX) is a widely used antiproliferative agent for treating cancer, autoimmune diseases, and chronic inflammatory disorders.^1–3^ Although clinically effective, MTX has a narrow therapeutic index.^4,5^ Accurate, individualized dosing is therefore essential to minimize adverse effects, including nephrotoxicity, hepatotoxicity, neurotoxicity, and myelosuppression, particularly during high-dose MTX therapy.^3–6^ Conventional therapeutic drug monitoring (TDM) of MTX relies on intermittent blood draws analyzed predominantly by competitive immunoassays,^7,8^ whereas chromatographic methods such as HPLC and LC-MS/MS serve as reference, confirmatory, or high-specificity approaches.^9,10^ Although these methods have clinically validated sensitivity and throughput, they are fundamentally centralized laboratory methods and are not readily adaptable to decentralized or high-frequency MTX measurements at the point of care. This constrains the temporal resolution of MTX monitoring, particularly for patients outside metropolitan areas.

Electrochemical aptamer-based (EAB) sensors have emerged as promising platforms for rapid and continuous monitoring of molecular targets in complex biological fluids.^11,12^ Aptamers are single-stranded nucleic acid ligands that are generated through *in vitro* selection and offer several important advantages over traditional affinity reagents such as antibodies. Unlike proteins, they are chemically stable, can be reproducibly synthesized, can be selected against native small molecules, and can exhibit rapid, reversible target recognition through their dynamic conformational behavior.^13–16^ These properties make aptamers attractive recognition elements for miniaturized electrochemical assays and, in particular, for continuous biochemical sensing platforms.^11,12^ Therapeutic drug monitoring has emerged as an important clinical application of continuous electrochemical aptamer-based sensing.^17–19^ MTX monitoring is particularly well suited to this approach, given its broad clinical use, substantial interpatient pharmacokinetic variability, and the need for careful control of systemic exposure.^3–5^

Several MTX-specific aptamers and aptamer-based sensing formats have been reported previously.^20–24^ However, the molecular basis of MTX recognition by these aptamers remains poorly understood. Here, we report two novel MTX-binding DNA aptamer families, 3315 and 930, developed using high-stringency Capture-SELEX.^25–27^ Both aptamer families demonstrated nanomolar binding affinities toward MTX, with the best-performing variants exhibiting stronger binding than previously reported MTX aptamers. Notably, the two aptamer families displayed markedly different binding behavior: the 3315-family retained strong target recognition across a broad range of magnesium concentrations, whereas the 930-family depended on Mg^2+^ for efficient target binding, suggesting distinct molecular recognition mechanisms. Systematic truncation identified compact functional variants, including 3315-53 and 930-65, that retained high binding performance while providing architectures suitable for sensor integration.

We combined computational modeling with site-directed mutagenesis to identify functional regions within the selected aptamer scaffolds. The optimized aptamers were integrated into EAB sensors that distinguished MTX from folic acid and operated directly in unprocessed human whole blood. Together, these findings establish two new high-affinity MTX aptamer families and demonstrate how integrating high-stringency aptamer selection with computational modeling, experimental validation, and electrochemical characterization provides a rational workflow for engineering aptamers for biosensing applications. This work delivers both new molecular recognition elements for MTX and a foundation for the future development of reagent-free, point-of-care therapeutic drug monitoring platforms.

## EXPERIMENTAL SECTION

Detailed protocols for Capture-SELEX, next-generation sequencing (NGS) library preparation and bioinformatic analysis, secondary-structure prediction, G-quadruplex analysis, and molecular dynamics (MD) simulations are provided in the Supporting Methods. Oligonucleotide sequences, buffer compositions, selection conditions, and computational parameters are summarized in Supporting Tables S1–S7.

### Isothermal Titration Calorimetry (ITC) experiments

Isothermal titration calorimetry (ITC) experiments were performed using a MicroCal PEAQ-ITC instrument (Malvern Panalytical) at 20 °C unless otherwise specified, with selected measurements additionally performed at 30 °C and 37 °C. Aptamers were prepared in the corresponding experimental buffers (Supporting Table S2), heated to 95 °C for 5 min, and gradually cooled to 4 °C over 30 min prior to analysis to ensure proper folding. Typically, 280 µL of 10 µM aptamer solution was loaded into the sample cell, while 40 µL of 100 µM MTX or counter-target solution was loaded into the injection syringe. Each experiment consisted of an initial purge injection of 0.4 µL, followed by 19 sequential injections of 2 µL at 150 s intervals, under continuous stirring at 750 rpm. Raw thermograms were corrected for heats of dilution and analyzed using MicroCal PEAQ-ITC Analysis Software. Binding isotherms were fitted using a single-site binding model to determine dissociation constants (K_D_) and thermodynamic parameters.

### Biolayer Interferometry (BLI) Analysis

Biolayer interferometry (BLI) was used to determine the binding affinity (K_D_) and kinetic parameters (k_on_ and k_off_) of 3315 and 930 aptamer variants toward MTX. All experiments were performed using an Octet RED96 system (ForteBio/Sartorius) equipped with streptavidin-coated SA biosensors (Sartorius) at 25 °C in black 96-well flat-bottom OptiPlate microplates (Revvity; Cat. No. 6008540), using a total volume of 200 µL per well.

Biotinylated aptamers containing BiotinTEG modification at the 5′ or 3′-end were prepared at 250 nM in selection buffer (SB) and renatured prior to analysis by heating to 95 °C followed by gradual cooling to room temperature. SA biosensor tips were hydrated and equilibrated in SB buffer for 10 min prior to measurements. Baseline acquisition was performed in SB buffer for 60 or 100 s, followed by aptamer immobilization onto the biosensor surface for 180 or 300 s. Unbound aptamers were removed by incubation in SB buffer for 60 or 100 s prior to association measurements.

Association kinetics were measured by transferring biosensors into wells containing serial dilutions of MTX or FA (0–30 µM) prepared in SB buffer for 50–100 s. Dissociation was monitored by transferring biosensors into SB buffer for 60 or 300 s. Binding kinetics were analyzed using Data Analysis HT 9.0 software (ForteBio) employing a global 1:1 binding model. Equilibrium K_D_ was additionally determined by plotting MTX concentration versus response (nm) using GraphPad Prism software.

### Strand displacement assay

Strand displacement assays were performed to evaluate target-induced conformational switching of selected MTX aptamers. FAM-labelled aptamers were mixed with complementary Iowa Black quencher-labeled oligonucleotides (Q-Iowa) at a molar ratio of 1:2 in selection buffer (SB). Briefly, 100 µM FAM-labelled aptamer and 100 µM quencher strand stock solutions were combined to obtain final concentrations of 0.1 µM aptamer and 0.2 µM quencher strand in the assay mixture. Aptamer–quencher complexes were renatured prior to use by heating followed by gradual cooling to room temperature. For fluorescence measurements, 50 µL of the aptamer–quencher mixture was combined with MTX solutions prepared in SB to obtain final MTX concentrations of 0, 0.5, 1, and 5 µM in black 96-well flat-bottom OptiPlate microplates (Revvity; Cat. No. 6008540). Samples were incubated for 30 min at room temperature with gentle agitation, protected from light. Fluorescence measurements were performed using a BioTek Synergy Neo2 plate reader at 22 °C with excitation at 487/20 nm and emission at 528/20 nm. MTX-only control solutions at identical concentrations without aptamer or quencher oligonucleotides were used for background correction, and the corresponding fluorescence signals were subtracted from assay measurements.

### Stability of Aptamers in Human Serum

FAM-labelled aptamer variants 3315-88, 3315-63, 3315-55, 3315-53, 930-88, and HMX38 were used to evaluate nuclease stability in human serum. Each aptamer was incubated at a final concentration of 2 µM in SB buffer supplemented with 95% human serum (Merck, H6914) in a total reaction volume of 100 µL at 37 °C. Aliquots of 10 µL were collected at 0, 0.5, 1, 2, 6, 24, and 48 h. Each aliquot was quenched by the addition of 20 µL of 2× gel loading dye and 10 µL of nuclease-free water, denatured at 95 °C for 10 min, and stored at ™20 °C until analysis. Prior to loading, samples were heated again at 95 °C for 10 min, and 10 µL of each sample was analyzed by 15% denaturing urea-PAGE. FAM-labelled DNA bands were visualized using a ChemiDoc MP Imaging System (Bio-Rad) with an Alexa Fluor 488 filter, and band intensities corresponding to intact aptamers were quantified using Image Lab Software (Bio-Rad).

### Electrochemical aptamer (EAB) sensor preparation and characterization

*Electrode functionalization:* Plated gold flexible electrodes (Linxens^®^) were functionalized with methylene blue (MB)-modified DNA aptamers specific for MTX. Two aptamer sequences, designated S-930-65-MB and S-3315-55-MB, were evaluated independently. Prior to immobilization, aptamer stock solutions were reduced using tris(2-carboxyethyl)phosphine (TCEP, Merck -Australia) to cleave disulfide bonds and generate free thiol groups for self-assembly onto the gold surface. The reduced aptamer solutions were prepared at a final concentration of 0.5 µM by mixing the appropriate volume of stock aptamer solution with TCEP prepared in RNase-free phosphate-buffered saline (PBS, Merck – Australia). The reduction mixture was incubated at 4 °C for 1.5 h before use. Following reduction, 30 µL of the aptamer solution was deposited onto the working electrode surface and incubated for 1 h at 4 °C in the dark inside a sealed Petri dish wrapped with aluminum foil to prevent photodegradation of the MB. After incubation, the electrodes were rinsed three times with 100 µL RNase-free PBS to remove unbound aptamer. To passivate the remaining gold surface and minimize nonspecific adsorption, 30 µL of 7.31 mM 6-mercapto-1-hexanol (MCH, Merck – Australia) was added to each electrode and allowed to incubate for 1 h at 4 °C in the dark. The electrodes were subsequently rinsed three times with RNase-free PBS and stored in PBS at 4 °C until electrochemical measurements were performed.

#### Electrochemical measurements and calibration curve

Electrochemical characterization was performed using a MultiEmStat4 LR four-channel potentiostat (Metrohm®) and the square-wave voltammetry (SWV) technique. Calibration curves were constructed by exposing the aptamer-functionalized electrodes to MTX solutions prepared at concentrations of 0 (PBS blank), 0.1, 1, 10, 20, 100, 200, 400, 600, 800, and 1000 µM. In experiments performed in human blood (obtained with ethics approval by the La Trobe University ethics committee under approval number 024-24885-55512-7). For each measurement, 100 µL of the MTX solution was added to the electrode surface, and electrochemical analyses were performed. All measurements were performed in triplicate. All the electrochemical analyses were conducted using square-wave voltammetry at a frequency of 600 Hz to identify

#### Selectivity evaluation

The selectivity of the aptamer-based electrochemical sensors was assessed using folic acid. Functionalized electrodes were incubated with 400 µM folic acid for 10 min at room temperature under the same conditions used for MTX measurements, followed by SWV analysis. All selectivity experiments were performed in triplicate.

#### Relative signal gain calculations

The relative signal change signal gain was calculated using the following equation

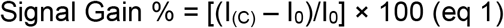

where I_0_ is the baseline current at zero MTX concentration and I_(C)_ is the given MTX concentration in PBS (SWV, 600 Hz).

## RESULTS AND DISCUSSION

### *In vitro* selection and initial characterization of MTX-binding aptamers

A library-immobilized Capture-SELEX strategy was employed to isolate DNA aptamers that recognize native MTX in solution (Figure 1a; Figures S1 and S2).^25–27^ To enrich for target-specific binders, selection stringency was increased over 14 rounds by progressively decreasing MTX concentration, increasing washing stringency, and using structurally similar folic acid (FA) for counterselection (Figure 1b,c; Tables S1–S3). Next-generation sequencing (NGS) analysis of the enriched pools (rounds 9–14) identified ten dominant sequence families (Table S4). Among these 88-nt aptamers, 3315-88 and 930-88 displayed MTX binding when analyzed by isothermal titration calorimetry (ITC; Figure 1d,e; Table S5). Fitting the data led to *K*_D_ values of 89 and 408 nM, respectively. Thermodynamic analysis showed that MTX binding was strongly enthalpy-driven, with favorable Δ*H* values ranging from ™24 to ™29 kcal mol^−1^ and unfavorable ™TΔ*S* values ranging from 14 to 22 kcal mol^−1^. This thermodynamic profile is consistent with the formation of specific intermolecular contacts and ligand-induced ordering of the aptamer structure upon MTX recognition (Table S5).^26,28^ When tested under the same experimental conditions, the previously reported MTX aptamers HMX38 and MTX40 displayed apparent *K*_D_ values of 291 and 1830 nM, respectively. ^20,22^

**Figure 1.**
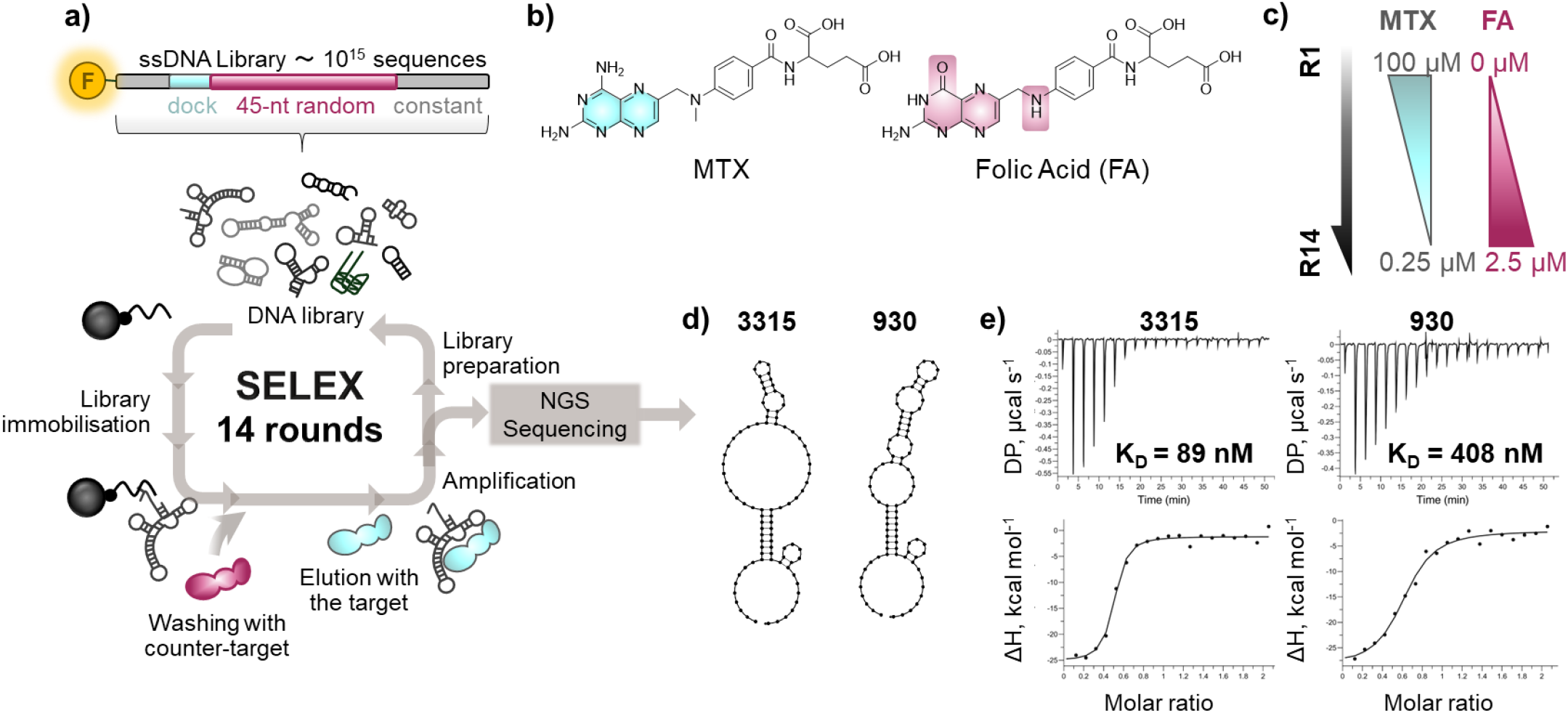
In vitro selection of methotrexate-specific aptamers. a) Design of the randomized DNA library and schematic overview of the Capture-SELEX strategy employing library immobilization and target-induced elution with native MTX. (b) Chemical structures of MTX and the counter-target folic acid (FA). (c) Schematic representation of the progressive selection stringency applied during the MTX selection campaign, including decreasing MTX concentration and increasing FA counter-selection pressure. (d) Predicted secondary structures of isolated full-length aptamers 3315-88 and 930-88 generated using NUPACK under DNA folding conditions (22 °C, 100 mM Na^+^, 2 mM Mg^2+^). (e) Representative ITC thermograms of MTX binding to selected aptamers obtained at 20 °C using 100 µM MTX in the syringe and 10 µM aptamer in the cell. Details are provided in Supporting Tables S1–S5 and Figure S1–S6.

Notably, the 3315 family was not detected among the dominant sequences until later selection rounds, becoming enriched only after increased selection stringency and the introduction of washing buffer containing human serum at round 11 (Tables S3 and S4). This enrichment pattern suggests that prolonged high-stringency selection was important for isolating high-affinity MTX-binding motifs. In addition, the 45 nt randomized region used in this study may have provided greater structural diversity than the 30 nt libraries previously used for MTX aptamer selection.^29^

Sequence and structure comparison identified only limited homology between the newly selected aptamers and previously reported MTX-binding sequences (Figure S3), indicating that MTX can be recognized by structurally diverse DNA scaffolds.^20–22^ Aptamer 3315-88 contained a prominent G-rich region with partial similarity to the G-rich HMX38 aptamer, which has previously been proposed to adopt a G-quadruplex structure. ^20^ However, when 3315-88 was tested in FLAG-ELISA and Direct-ELISA using G-quadruplex-specific nanobody SG4 it produced the same signal as the non-G-quadruplex control mMycG4, suggesting that 3315-88 does not form a canonical G-quadruplex under tested conditions (Figure S4).^30–32^ Biolayer interferometry (BLI) measurements revealed rapid and reversible MTX recognition by both aptamers (Figure S5). In addition, fluorescent strand-displacement assays performed with 3315-88, 930-88 and HMX38 aptamers all revealed MTX-dependent increases in fluorescence.^20^ Interestingly, 930-88 produced the strongest signal change despite its lower affinity (Figure S6). This suggests that 930-88 undergoes a more pronounced target-induced conformational rearrangement, which may be advantageous for structure-switching sensor formats.

### Specificity, temperature, and matrix dependence of aptamers 3315 and 930

For aptamers intended for therapeutic drug monitoring, performance evaluation under physiologically relevant conditions, including elevated temperature, variable ionic composition, and serum-containing environments, is essential. We therefore systematically investigated the binding properties of aptamers 3315-88 and 930-88 by ITC across different temperatures, buffer compositions, matrix conditions, and analytes (Figure 2).

**Figure 2.**
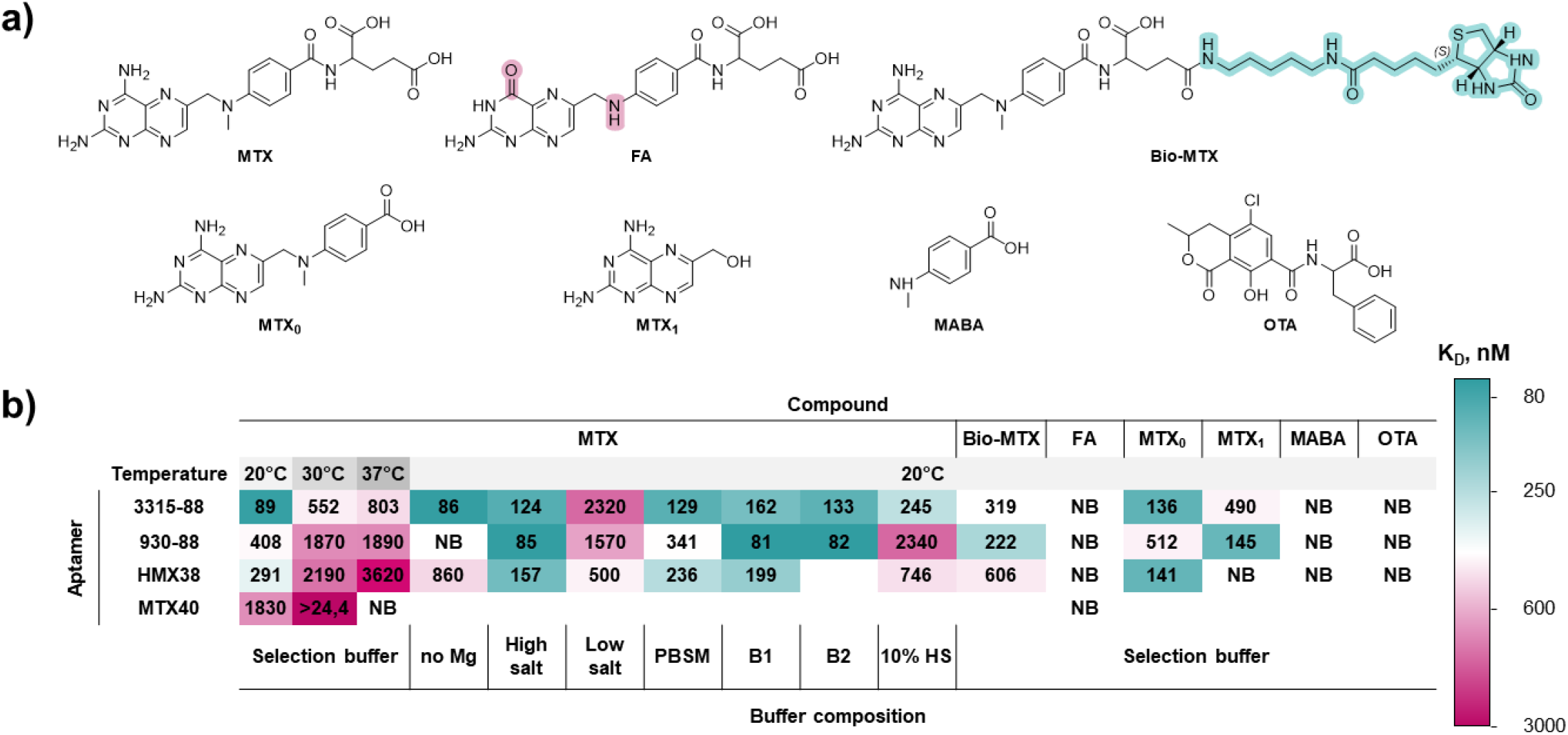
Affinity and specificity analysis of MTX-specific aptamers. (a) Chemical structures of MTX and compounds used for specificity assessment by ITC, including biotinylated-MTX (Bio-MTX), folic acid (FA), 4-deoxy-4-amino-N10-methylpteroic acid (MTX_0_ or DAMPA), 2,4-diamino-6-(hydroxymethyl)pteridine (MTX_1_), 4-methylaminobenzoic acid (MABA), and ochratoxin A (OTA). (b) Heat map of dissociation constants (K_D_, nM) determined by ITC for aptamers 3315-88 and 930-88 together with previously reported MTX aptamers HMX38 and MTX40 under different temperatures, buffer conditions, and analyte compositions. ITC measurements were performed at 20, 30, or 37 °C using 100 µM analyte in the syringe and 10 µM aptamer in the cell. The previously reported aptamer MSmt7 did not demonstrate detectable binding under the experimental conditions used in this study. Detailed buffer compositions and representative ITC isotherms are provided in Figures S7–S10 (Supporting Information).

To evaluate thermal stability, we performed ITC measurements at 20, 30, and 37 °C using 3315-88 and 930-88, with HMX38, MTX40, and MSmt7 as controls.^20–22^ Under the experimental conditions used in this study (100 µM analyte in the syringe and 10 µM aptamer in the cell), MSmt7 did not produce detectable binding isotherms and was therefore excluded from further comparative analysis (Figure S7).^21^

All evaluated aptamers showed reduced affinity at elevated temperatures, consistent with thermal destabilization of aptamer secondary structure and weakened noncovalent target interactions. However, 3315-88 remained stronger MTX binding at physiological temperature than HMX38 and MTX40 tested under the same conditions. The affinity of 3315-88 decreased from 89 nM at 20 °C to 803 nM at 37 °C, whereas the other evaluated aptamers displayed apparent *K*_D_ values above 1 µM at 37 °C. The observed thermal tolerance should enable use of 3315-88 in MTX biosensing applications operating at physiological temperatures.

We next investigated how ionic strength and matrix composition affect aptamer binding (Figure 2 and S8). Aptamer 3315-88 demonstrated broad buffer tolerance, maintaining *K*_D_ values between 86 and 245 nM under most tested conditions, including PBS supplemented with Mg^2+^, albumin-containing buffer B2, and selection buffer containing 10% human serum. Removing Mg^2+^ had minimal effect on 3315-88 binding, suggesting that this aptamer recognizes MTX largely independently of Mg^2+^. In contrast, reduced-ionic-strength conditions containing only 20 mM NaCl substantially weakened binding (*K*_D_ ≈ 2.3 µM), indicating that monovalent cations stabilize the active aptamer conformation. This observation is consistent with the known role of Na^+^ in shielding electrostatic repulsion between phosphate backbones and stabilizing compact nucleic acid folds. ^33^ Overall, 3315-88 is a robust and buffer-tolerant MTX-binding scaffold.

In contrast, aptamer 930-88 exhibited a distinct Mg^2+^-dependent binding profile. Under standard selection-buffer conditions, 930-88 bound MTX with an apparent *K*_D_ of 408 nM; however, increasing the MgCl_2_ concentration to 5–10 mM improved affinity to 81–85 nM. No detectable binding was observed without Mg^2+^, indicating that divalent cations are essential for stabilizing the binding-competent conformation of 930-88. Mg^2+^-dependent folding is commonly observed in structure-switching aptamers and riboswitch-like nucleic acid architectures, where divalent ions facilitate tertiary structure formation and ligand-induced conformational transitions.^34^ The ability to modulate 930-88 activity via Mg^2+^ concentration suggests that this aptamer may function as an ion-dependent conformational switch, a property that could be useful for structure-switching biosensor formats.

Interestingly, the addition of recombinant albumin (100 µg mL^−1^) to buffer B2 did not substantially affect MTX binding by either aptamer, with apparent *K*_D_ values of 133 and 82 nM for 3315-88 and 930-88, respectively. In contrast, 10% human serum had different effects on the aptamers (Figure S8 and S9). While 3315-88 retained strong MTX binding in serum-containing buffer (*K*_D_ = 245 nM), the affinity of 930-88 decreased more than fivefold (*K*_D_ ≈ 2.3 µM). This matrix sensitivity may reflect nonspecific interactions with serum components, altered ionic composition, reduced Mg^2+^ availability, or changes in aptamer folding and target accessibility.

Finally, aptamer specificity was evaluated against MTX and structurally related or unrelated compounds, including folic acid (FA), 4-deoxy-4-amino-N10-methylpteroic acid (MTX_0_/DAMPA), 2,4-diamino-6-(hydroxymethyl)pteridine (MTX_1_), 4-methylaminobenzoic acid (MABA), and ochratoxin A (OTA) (Figure 2a). MTX_0_/DAMPA is a clinically relevant MTX degradation product generated during glucarpidase rescue therapy,^4,35^ making discrimination between MTX and MTX_0_ important in this clinical context. MTX_1_ and MABA were included as structural fragments to probe aptamer recognition features, while OTA served as an unrelated aromatic small-molecule control.

Previously, Nagano et al. reported that MSmt7 showed limited discrimination between MTX and structurally related antifolates, including MTX_0_ and pralatrexate, while displaying selectivity against folic acid, 7-hydroxy-MTX, leucovorin, and pemetrexed.^21^ He et al. showed that HMX38 recognized MTX over several unrelated small molecules, including vancomycin, simvastatin, penicillin, omeprazole, and FA,^20^ while Wu et al. did not report specificity testing of MTX40 against structurally related MTX analogs or FA.^22^

Both 3315-88 and 930-88 showed negligible binding to OTA, MABA, and FA, reflecting selective recognition of the MTX scaffold rather than nonspecific binding to aromatic or folate-like compounds (Figure S10). However, both aptamers also recognized MTX_0_ with affinities comparable to MTX (3315-88: 136 nM vs 89 nM; 930-88: 512 nM vs 408 nM). The previously reported HMX38 aptamer similarly recognized MTX_0_ and, under our experimental conditions, bound MTX_0_ more strongly than MTX itself (141 nM vs 291 nM). Therefore, discrimination between MTX and MTX_0_ remains a limitation of current MTX-binding aptamer scaffolds. This limitation is most relevant in the context of glucarpidase rescue therapy, where DAMPA formation can be substantial, and should be considered in future sensor development. This does not apply to routine high-dose MTX clearance monitoring.^4,35^

### Truncation identifies minimal functional MTX-binding variants

After initial characterization, we systematically truncated the aptamer to define the minimal sequence elements required for MTX binding and to generate compact variants suitable for sensor integration (Figure 3 and Figures S11–S21). For aptamer 3315, sequential truncation preserved and, in some cases, improved MTX binding.

**Figure 3.**
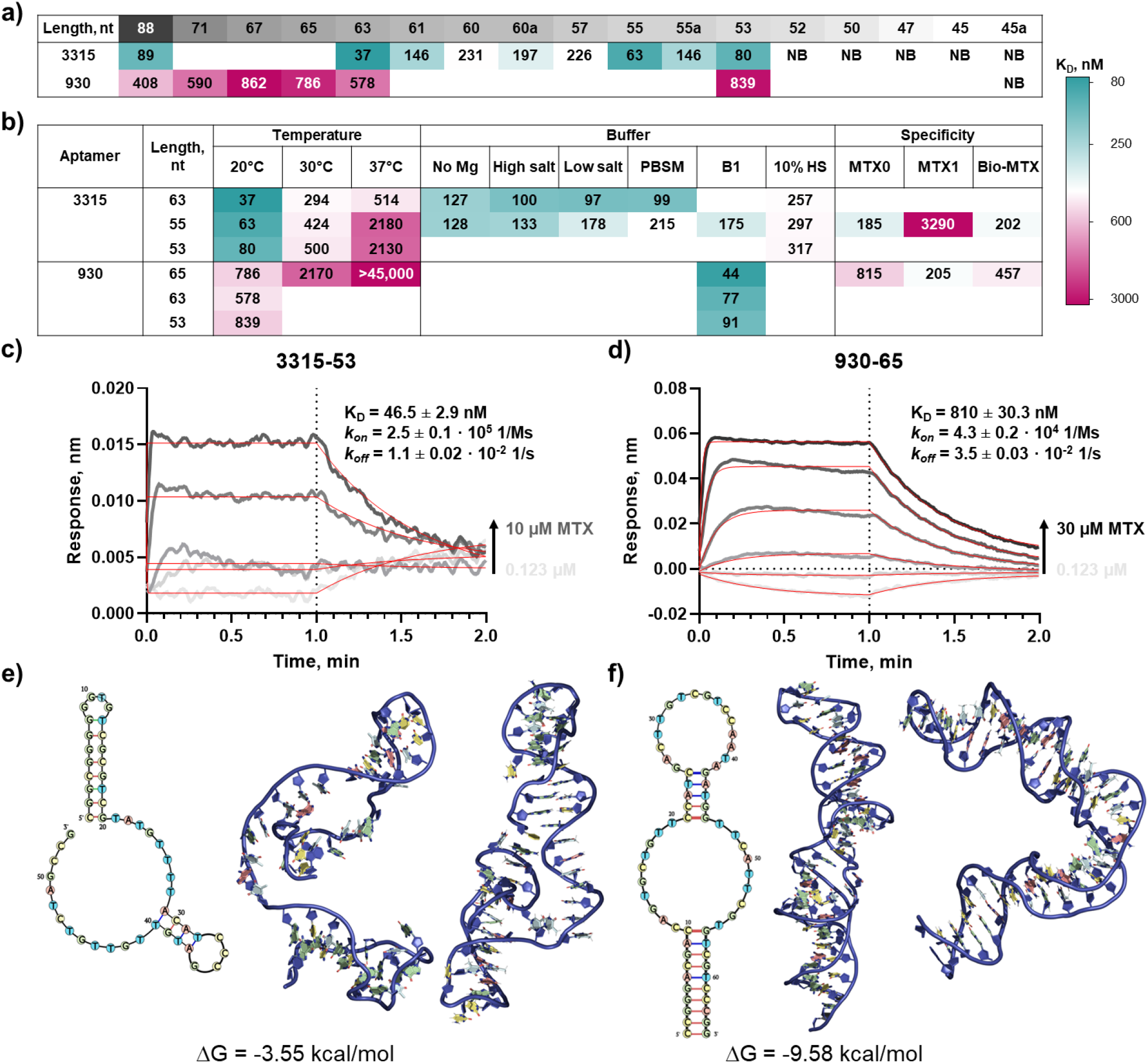
Truncation, binding characterization, and MD-refined structural models of MTX aptamers 3315 and 930. (a)Summary of the sequential truncation of full-length aptamers 3315-88 and 930-88 and MTX-binding analysis by ITC at 20 °C in selection buffer. Values are reported as apparent *K*_D_ in nM; NB indicates no detectable binding under the tested conditions, and blank cells indicate variants that were not tested. (b) ITC-based characterization of selected truncated variants under different temperature, buffer, matrix, and specificity conditions. Buffer conditions included selection buffer without Mg^2+^, high-salt buffer, low-salt buffer, PBS supplemented with Mg^2+^ (PBSM), B1 buffer containing 10 mM Mg^2+^, and buffer containing 10% human serum (10% HS). Specificity measurements were performed using MTX-derived compounds, including MTX_0_, MTX_1_, and biotinylated MTX. Values are shown as apparent *K*_D_ (nM) and color-coded according to binding affinity. (c,d) BLI characterization of MTX binding to 3315-53 (c) and 930-65 (d). (e,f) Predicted secondary structure, initial 3D model, and MD-refined apo conformation of 3315-53 (e) and 930-65 (f). Computational models were generated using secondary-structure-guided 3D modeling followed by molecular dynamics refinement. The Supporting Information provides detailed ITC isotherms, BLI experimental conditions, buffer compositions, structural prediction parameters, and MD simulation details.

Compared with the full-length 3315-88 aptamer (*K*_D_ = 89 nM), the 63-nt, 55-nt, and 53-nt variants displayed stronger affinities of 37, 63, and 80 nM, respectively (Figure 3a and Figures S11 and S12). Other truncations weakened binding, and variants of 52 nt or shorter showed no detectable interaction. These results indicate that the active 3315 scaffold is retained within the 53–55 nt region and that selected terminal residues from the original primer-flanking regions contribute to the binding-competent structure rather than acting as redundant sequence.

Unlike 3315, simple 930 aptamer truncation did not improve MTX binding under standard selection-buffer conditions (Figure 3a and Figures S13 and S14). However, in Mg^2+^-rich B1 buffer, the 930-65, 930-63, and 930-53 variants showed substantially enhanced MTX binding, with apparent *K*_D_ values of 44, 77, and 91 nM, respectively (Figure 3b and Figure S15). This behavior is consistent with the Mg^2+^ dependence observed for full-length 930 and supports a model in which divalent cations stabilize the productive MTX-binding conformation of this aptamer family.

A subset of truncated variants was evaluated under conditions relevant to biosensor development, including elevated temperature, altered buffer composition, 10% human serum, and MTX-derived analytes (Figure 3b and Figures S16–S20). Among the 3315 variants, 3315-63 showed the strongest overall thermal robustness, retaining measurable MTX binding across the 20–37 °C range. The shorter 3315-55 and 3315-53 variants also remained active but showed larger affinity losses at elevated temperatures. Buffer-dependence experiments further confirmed that 3315-63 and 3315-55 retain binding across multiple ionic conditions, including Mg^2+^-free, high-salt, low-salt, and PBSM buffers, supporting the Mg^2+^-independent behavior of the 3315 scaffolds (Figure S18). Importantly, all three tested 3315 truncation variants retained measurable MTX binding in a 10% human serum-containing buffer, with apparent *K*_D_ values of 257–317 nM (Figure S19). Although serum reduced apparent affinity relative to the selection buffer, retention of nanomolar binding in this matrix supports the suitability of the 3315 scaffolds for practical applications.

Specificity analysis showed that truncation preserved recognition of the MTX scaffold while improving analog discrimination for the 3315 family (Figure S20). The 3315-55 variant retained strong MTX binding (*K*_D_ = 63 nM) and showed more than 3-fold weaker binding to MTX_0_/DAMPA and Bio-MTX, with substantially weaker binding to MTX_1_ (*K*_D_ = 3.3 µM). This attests to improved selectivity over the parental 3315-88 aptamer, which binds MTX_0_ with an affinity closer to that of MTX (89 vs 136 nM). In contrast, 930-65 retained measurable binding to MTX-derived compounds, including MTX_0_, MTX_1_, and Bio-MTX, indicating less pronounced discrimination among related MTX fragments. For both truncated aptamer families, we observed no detectable binding to folic acid, MABA, or OTA, supporting selective recognition of MTX-related structural features rather than nonspecific binding to unrelated aromatic or folate-like molecules.

BLI independently confirmed MTX binding by the optimized variants (Figure 3e,f and Figure S21). In selection buffer, 3315-53 and 930-65 showed concentration-dependent binding with apparent kinetic K_D_ values of 46.5 and 810 nM, respectively, consistent with the relative affinities measured by ITC. At 10 mM Mg^2+^, apparent K_D_ values were 76.7 and 593 nM, respectively. Both aptamers exhibited rapid association and dissociation, whereas folic acid produced minimal responses, further supporting selective MTX recognition (Figure S21). CD spectroscopy further indicated MTX-induced conformational changes in both optimized aptamers (Figure S22). Adding 100 μM MTX altered the intensity and shape of the characteristic DNA CD bands, with a more pronounced change for 930-65, consistent with target-induced rearrangement of the aptamer structure.

Because biosensor application in biological samples also requires resistance to nuclease degradation, selected aptamers were evaluated in 95% human serum at 37 °C (Figure S23). Among the truncated variants, 3315-55 showed the best stability, retaining approximately 60% of the intact sequence after 2 h and displaying an estimated serum half-life of approximately 4 h. Other truncated variants degraded more rapidly, indicating that minimization can compromise nuclease resistance even when target binding is retained. Overall, truncation revealed distinct optimization outcomes for the two MTX-binding aptamer families. The 3315 scaffold tolerated minimization and yielded compact variants with strong affinity, broad buffer tolerance, measurable serum-compatible binding, and improved suitability for integration with surface-based sensors. In parallel, 930 retained a stronger dependence on ionic conditions, particularly Mg^2+^, but remained functional after substantial sequence minimization.

### Molecular Dynamics (MD)-guided binding-site prediction and NGS conservation analysis

To link the biochemical binding data to the structural mechanism, we performed MD simulations on 3315-53 and 930-65 aptamers. Following 3D modeling and 500 ns apo-aptamer MD refinement, aptamer 3315-53 adopted a compact conformation in which an exposed central unpaired region formed a well-defined cavity suitable for ligand recognition (Figure 3e,f). In contrast, aptamer 930-65 maintained a more elongated, screw-like architecture with multiple possible binding pockets distributed across paired and unpaired regions. Cluster analysis indicated that the dominant conformations represented approximately 30% and 80% of the apo trajectories for 3315-53 and 930-65,^36^ respectively, suggesting that 930-65 may adopt a more conformationally restricted apo state, whereas 3315-53 samples a broader ensemble before ligand binding.

Pocket prediction using Fpocket^37^ and DoGSite3^38,39^ identified one major MTX-binding candidate site in 3315-53, involving nucleotides within the central unpaired region (Figure 4). For 930-65, three potential binding pockets were identified and evaluated independently. Docking analysis using GOLD supported favorable MTX binding to both aptamers,^40^ with a higher docking score for 3315-53 than for 930-65 (Table S6). Subsequent 300 ns MD simulations of the docked complexes showed further differences in aptamer behavior. MTX remained stably associated with the predicted 3315-53 binding pocket throughout the full simulation, whereas only one of the three 930-65 docking poses remained stable, corresponding to the pocket involving nucleotides 11–15 and 51–57 (pocket 2; Figure 4). The other 930-65 complexes dissociated within the first 100 ns, suggesting that only a subset of the predicted pockets can support stable productive interaction.

**Figure 4.**
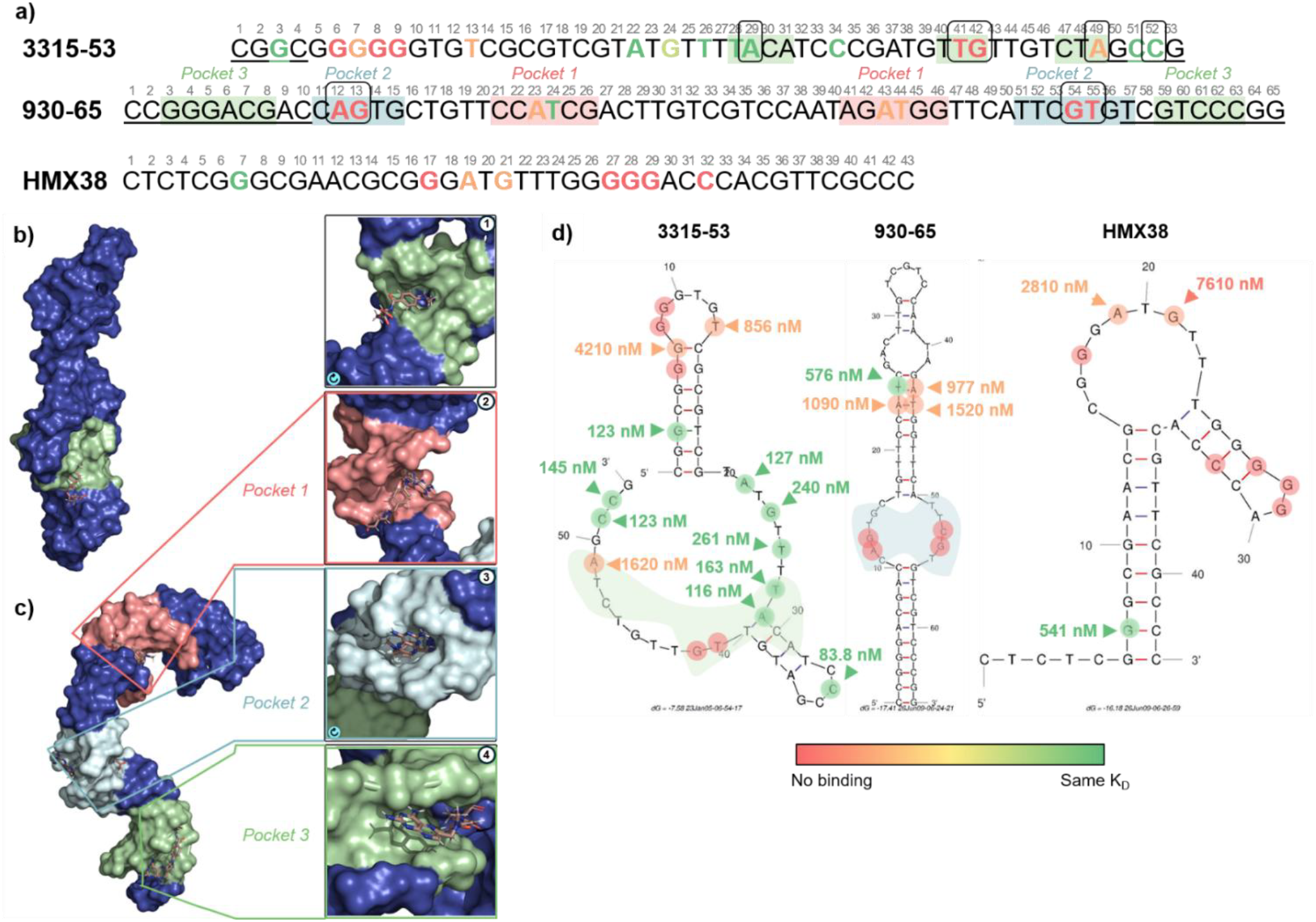
Mutational validation of MD-predicted MTX-binding sites in aptamers 3315-53 and 930-65. (a) Sequence maps of 3315-53, 930-65, and HMX38 showing predicted MTX-binding regions and nucleotides selected for site-directed mutagenesis. Mutated residues are colored according to their effect on apparent MTX-binding affinity as measured by ITC: red indicates no detectable binding, orange indicates substantially weakened binding, and green indicates retained binding with only minor effects on affinity. Nucleotides predicted by MM-GBSA per-nucleotide decomposition to contribute most strongly to MTX recognition are boxed. Primer-derived regions are underlined. (b) MD-refined model of the MTX– 3315-53 complex showing the predicted MTX-binding pocket. (c) MD-refined model of the MTX–930-65 complex showing three candidate binding pockets. (d) Summary of ITC-derived apparent K_D_ values for mutant aptamers mapped onto predicted secondary structures of 3315-53, 930-65, and HMX38. Full ITC thermograms are provided in the Supporting Information.

Molecular Mechanics/Generalized Born Surface Area (MM-GBSA) analysis of the stable complexes further supported stronger MTX binding by 3315-53,^41^ with a calculated binding free energy of ™10.0 ± 4.3 kcal mol^−1^ compared with ™4.4 ± 3.3 kcal mol^−1^ for 930-65 (Table S7). This trend is consistent with the experimentally determined ITC affinities under standard selection buffer conditions, where 3315-53 displayed stronger MTX binding (K_D_ = 80.3 nM, ΔG = ™9.52 kcal mol^−1^; Figure S12) than 930-65 (K_D_ = 786 nM, ΔG = ™8.19 kcal mol^−1^; Figure S14). For both aptamer–MTX complexes, binding was driven primarily by favorable nonpolar contributions, including van der Waals interactions and nonpolar solvation terms, whereas the net polar contribution was unfavorable. Per-nucleotide energy decomposition identified A29, T41, G42, A49, and C52 as the major contributors to MTX binding in 3315-53, while A12, G13, G54, and T55, located within pocket 2, contributed most strongly to MTX binding in 930-65. Notably, the key residues predicted for 3315-53 are retained within the shortest active 3315 construct, whereas further truncation beyond this region removes essential structural elements and abolishes detectable MTX binding. This supports the experimental observation that 3315-53 represents a minimal binding-competent scaffold, while shorter 3315 variants lack critical determinants required for productive target recognition. Overall, the simulations provide a structural rationale for the higher affinity and robustness of the 3315 scaffold and suggest that 930-65 relies on a more conditional binding mode, consistent with its experimentally observed Mg^2+^ dependence and enhanced performance under Mg^2+^-rich buffer conditions.^42–44^

### Mutational validation of predicted MTX-binding sites

To experimentally validate the MD-predicted MTX-binding regions, site-directed mutagenesis was performed on 3315-53 and 930-65 (Figure 4 and Figures S24–S27). For 3315-53, mutations within the predicted binding region had strongly position-dependent effects. Substitution of T41 or G42 abolished detectable MTX binding, whereas mutation of A49 substantially weakened binding. In contrast, A29C and C52A were tolerated. These data indicate that the predicted 3315 binding region contains both essential and permissive positions, with T41, G42, and A49 forming critical determinants of the MTX-binding interface.

Additional mutations were introduced to test whether stabilizing the predicted stem-loop architecture could improve binding to 3315-53. Variants containing stem-stabilizing substitutions, including G3A, T13C, and the combined G3A/T13C/G6C mutant, did not improve affinity and, in some cases, abolished detectable binding (3315-53a,b,c; Figure S24c). This result indicates that MTX recognition by 3315-53 depends not only on global structural stability but also on the precise balance between stem stability, local flexibility, and binding-pocket geometry.

We also investigated the G-rich region of 3315-53 because it partially resembles the G-rich motif in the previously reported HMX38 MTX aptamer (Figure S24e).^20^ Mutations within the G6–G9 cassette of 3315-53 strongly reduced or abolished MTX binding, indicating that this region contributes either directly to ligand recognition or indirectly to the formation of the binding-competent scaffold. Although HMX38 has previously been associated with G-quadruplex-like folding, SG4-based analysis showed that 3315 does not form a stable canonical G-quadruplex under the tested conditions. Mutational analysis of HMX38 nevertheless showed that selected G-rich elements, particularly G17 and the G27–G29 region, are critical for MTX recognition (Figure S25). Together,these results suggest that G-rich elements can support MTX binding through distinct structural arrangements, including canonical G4-like folds in some aptamers and noncanonical G-rich architectures in others.

Unlike 3315-53 and HMX38, aptamer 930-65 is predicted to possess a different binding architecture and lacks the extended G-rich cassette. We computationally identified three candidate binding pockets; however, we did not prioritize pocket 3 for mutational validation because it lies largely within primer-derived terminal residues and is therefore less likely to represent the original SELEX-enriched recognition motif. Mutations in pocket 2, including A12C, G13T, G54T, and T55G, abolished detectable MTX binding (Figure S26). In contrast, mutations in pocket 1 produced only minor to moderate effects on affinity. These data identify pocket 2 as the most likely MTX-binding site in 930-65, consistent with MD simulations showing stable MTX retention only in this pocket.

Importantly, scrambled control variants of 3315-53 and 930-65 showed no detectable MTX binding by ITC, further confirming that target recognition is sequence- and structure-dependent rather than arising from nonspecific interaction between MTX and DNA (Figure S27). Overall, the combined MD, MM-GBSA, and mutagenesis data reveal two distinct MTX-recognition architectures. Aptamer 3315-53 relies on a compact G-rich scaffold coupled to a central binding interface, where both nucleotide identity and local structural flexibility are essential. Aptamer 930-65, in contrast, depends on a more localized pocket 2 architecture, consistent with its stronger Mg^2+^ dependence and conditionally stabilized binding behavior. These mechanistic differences help explain the experimentally observed divergence between the robust, buffer-tolerant 3315 scaffold and the more ion-dependent, structure-switching behavior of 930.

### Electrochemical aptamer-based detection of MTX

Both optimized aptamers were evaluated as recognition elements in electrochemical aptamer-based (EAB) sensors (Figure 5 and Figures S28–S31).^11,24^ 3315-55 and 930-65 were immobilized on gold electrodes through a 5′ C6-thiol linker and modified with a 3′ methylene blue redox reporter (Figure 5a). Target binding alters aptamer conformation and dynamics, thereby modulating reporter–electrode electron transfer, which was monitored by square-wave voltammetry (SWV).

**Figure 5.**
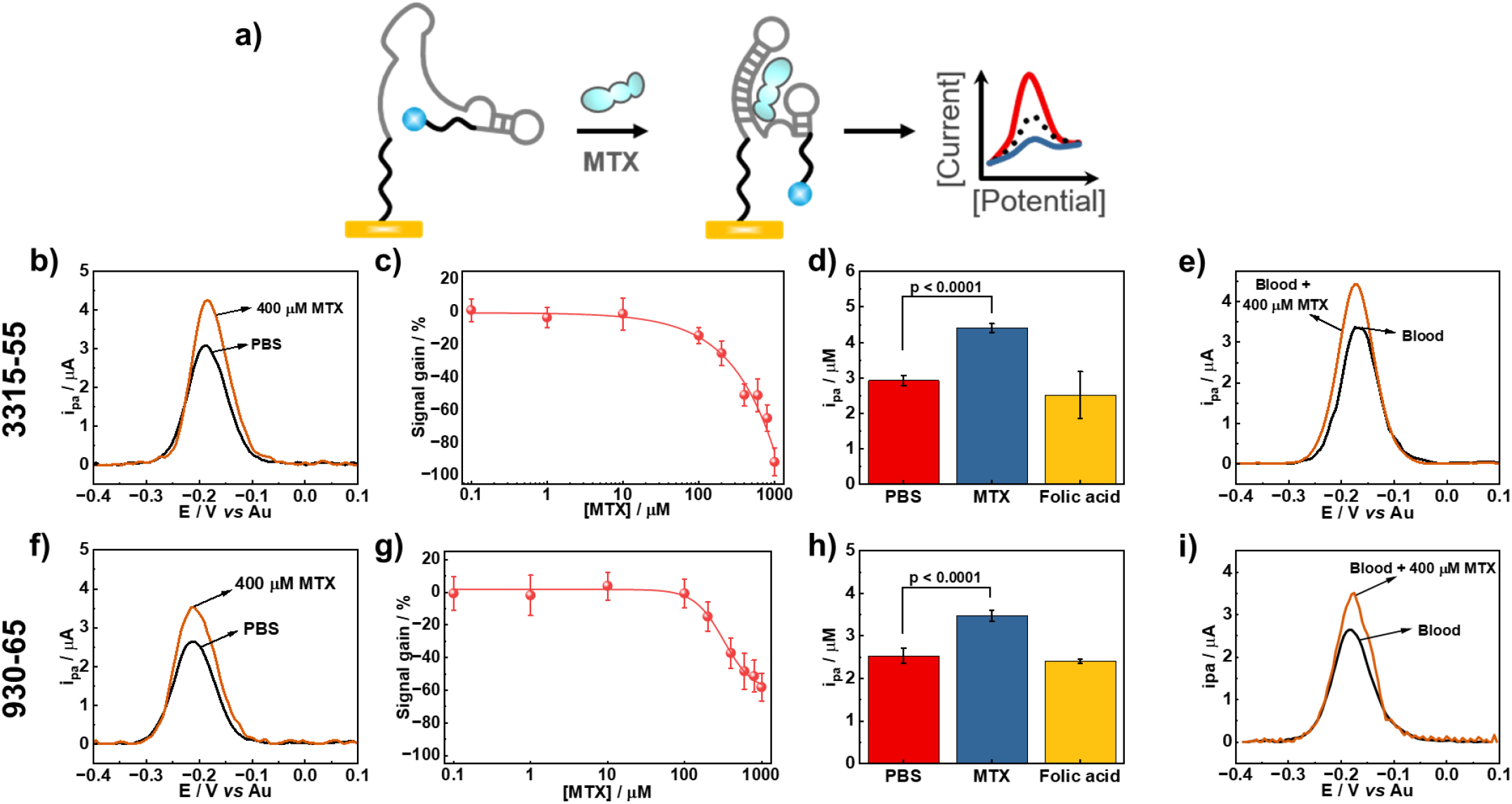
Design and validation of MTX EAB sensors. (a) Schematic of the electrode-bound, methylene-blue-modified aptamer sensor and target-induced modulation of electron transfer. (b–e) Characterization of the 3315-55 EAB sensor: (b)SWV response to 400 µM MTX, (c) concentration-dependent response in PBS (0.1–1000 µM; lowest detectable concentration, 100 µM), (d) selectivity against folic acid (400 µM), and (e) response to MTX in undiluted human blood. (f–i) Corresponding characterization of the 930-65 EAB sensor: (f) response to 400 µM MTX, (g) concentration-dependent response in PBS (0.1–1000 µM; lowest detectable concentration, 100 µM), (h) selectivity against folic acid (400 µM), and (i) response to MTX in undiluted human blood. Data are mean ± SD from n = 3 electrodes; SWV measurements were performed at 600 Hz.

Consistent with the MTX binding observed by ITC and BLI, 3315-55 generated a clear EAB signal change upon addition of 400 µM MTX in PBS (Figure 5b). Concentration-dependent measurements from 0.1 to 1000 µM MTX showed a detectable response from 100 µM under the tested conditions (Figure 5c). The sensor showed minimal response to 400 µM folic acid compared with MTX (Figure 5d), consistent with the molecular selectivity observed during solution-phase aptamer characterization. Importantly, 3315-55 also produced an MTX-dependent signal in undiluted human blood, demonstrating retention of sensor function in a complex biological matrix (Figure 5e).

The independently selected 930-65 aptamer showed comparable behavior in the EAB format. Adding 400 µM MTX produced a clear SWV signal change (Figure 5f), and concentration-dependent measurements again yielded a detectable response at 100 µM MTX (Figure 5g). 930-65 similarly discriminated MTX from folic acid at 400 µM (Figure 5h) and retained MTX-responsive signaling in undiluted human blood (Figure 5i).

Thus, despite differences in their solution-phase binding affinities and structural characteristics, both independently selected aptamers successfully translated into EAB sensors with similar analytical behavior. Their preserved MTX selectivity and function in whole blood further support the compatibility of the selected aptamer scaffolds with electrochemical sensing and their potential for further development toward therapeutic drug monitoring.

## CONCLUSION

In this work, we discovered, optimized, and mechanistically characterized two new DNA aptamer families for methotrexate recognition and demonstrated their translation into electrochemical sensors. High-stringency Capture-SELEX with native MTX elution and folic acid counter-selection yielded two structurally distinct aptamers, 3315 and 930, with nanomolar affinity. Importantly, the two families exhibited fundamentally different recognition behavior: 3315 formed a robust, buffer-tolerant scaffold with limited Mg^2+^ dependence, whereas 930 showed strongly Mg^2+^-dependent binding, revealing two distinct molecular solutions for recognizing the same small-molecule drug.

Systematic truncation generated compact variants suitable for sensor integration. 3315-55 combined high affinity (K_D_ = 63 nM), broad buffer tolerance, and comparatively high serum stability, retaining approximately 60% intact sequence after 2 h in 95% human serum with an estimated half-life of ∼4 h. In contrast, truncation of the 930 family did not substantially enhance binding under standard selection-buffer conditions, but 930-65 reached an apparent K_D_ of 44 nM in 10 mM Mg^2+^, corresponding to an ∼18-fold affinity enhancement relative to standard selection buffer and highlighting its ion-dependent folding behavior.*

Molecular dynamics simulations, docking, MM-GBSA analysis, and site-directed mutagenesis linked these distinct biochemical phenotypes to different MTX-recognition architectures. The 3315 scaffold depends on a compact G-rich architecture coupled to a central binding interface, with T41, G42, and A49 forming critical determinants of productive recognition. In contrast, 930 relies on a localized binding pocket involving A12, G13, G54, and T55. The agreement between computational predictions and mutational validation demonstrates the value of integrating molecular modeling with experimental structure–function analysis for rational aptamer optimization.

Finally, we successfully converted both optimized aptamers into electrochemical aptamer-based sensors. S-3315-55-MB and S-930-65-MB produced concentration-dependent MTX responses, resolved MTX concentrations from 100 µM under the tested conditions, discriminated MTX from folic acid in PBS, and retained MTX-responsive signaling in undiluted human blood. Together, these results establish two new high-affinity MTX recognition scaffolds and, more broadly, demonstrate a structure-guided workflow that connects aptamer selection, molecular mechanism, experimental validation, and electrochemical transduction. This integrated strategy provides a foundation for the rational development of small-molecule aptamers and their future deployment in reagentless therapeutic drug monitoring and point-of-care sensing.

## ACKNOWLEDGMENTS

The authors thank Victoria Coyne for assistance with NGS data collection. We thank Patricia Walden and Maria Micaela Fiorito for their invaluable technical support, laboratory assistance, and management. E.E., T.C.P., and S.M.S. acknowledge support from the Australian Research Council (ARC) Research Hub for Molecular Biosensors at Point-of-Use (MOBIUS; IH240100013). This work was partially funded by the ARC Center of Excellence in Synthetic Biology CE200100029 to K.A. and A.B., the ARC Linkage grant 200200916, and the NHMRC Investigator grant APP 2033951 to K.A. K.A. gratefully acknowledges the financial support of the CSIRO-QUT Synthetic Biology Alliance as well as the support of the CSIRO Synthetic Biology and Advanced Engineering Biology Future Science Platforms. This work was carried out with the support of the CERES Centre and supported by national funds from FCT through the projects https://doi.org/10.54499/UID/00102/2025 and https://doi.org/10.54499/UID/PRR/00102/2025 to S.F.S. S.F.S also received financial support from the PT national funds (FCT/MECI, Fundação para a Ciência e Tecnologia and Ministério da Educação, Ciência e Inovação) through the project UID/50006/2025 DOI 10.54499/UID/50006/2025 -Laboratório Associado para a Química Verde -Tecnologias e Processos Limpos.

