## SUPPORTING INFORMATION for "Structure-Guided Discovery of High-Affinity Methotrexate Aptamers for Electrochemical Drug Monitoring"

[†]Authors contribute equally

[1]Centre for Agriculture and the Bioeconomy, School of Biology and Environmental Science, Queensland University of Technology, Gardens Point Campus, Brisbane, QLD, 4000, Australia

[2]The Biomedical and Environmental Sensor Technology (BEST) Research Centre, Biosensors Program, La Trobe Institute for Molecular Science (LIMS), La Trobe University, Bundoora, VIC, 3086, Australia

[3]Department of Biochemistry and Chemistry, School of Agriculture, Biomedicine and Environment, La Trobe University, Bundoora, VIC, 3086, Australia

[4]LAQV/REQUIMTE, BioSIM, Department of Biomedicine, Faculty of Medicine, University of Porto, Rua Alameda Prof. Hernâni Monteiro, Porto, 4200-319, Portugal

[5]ARC Centre of Excellence in Synthetic Biology, Canberra, Australian Capital Territory, Australia

[6]Pacific Northwest National Laboratory Richland, WA 99354, United States

[7]CERES, BIA Lab, Department of Chemical Engineering, FCTUC, University of Coimbra, Rua Sílvio Lima, Pólo II – Pinhal de Marrocos, 3030-790 Coimbra, Portugal

[8]ARC Research Hub for Molecular Biosensors at Point-of-Use (MOBIUS), La Trobe University, Melbourne, VIC 3086, Australia

[9]Aptamer Facility, Synthetic Biology Program, La Trobe Institute for Molecular Science (LIMS), La Trobe University, Bundoora, VIC, 3086, Australia

***Corresponding author: Elena Eremeeva,** Centre for Agriculture and the Bioeconomy, Queensland University of Technology, Brisbane, Queensland, Australia. School of Biology and Environmental Science, Queensland University of Technology, Brisbane, 4000, Queensland, Australia; Aptamer Facility, Synthetic Biology Program, La Trobe Institute for Molecular Science (LIMS), La Trobe University, Melbourne, Victoria, 3086, Australia.

### Supporting Methods

#### In Vitro Selection of MTX Aptamers by Capture-SELEX

DNA aptamers specific to methotrexate (MTX) were generated using a library-immobilized Capture-SELEX strategy, following previously reported protocols with minor modifications.[1–4] All oligonucleotides used in this study were synthesized by Integrated DNA Technologies (https://sg.idtdna.com/page) with standard desalting purification (Supporting Table S1). Oligonucleotides were dissolved in molecular biology-grade water and quantified using a NanoDrop 2000 spectrophotometer (Thermo Fisher Scientific).

The initial DNA library (L45), containing approximately $10^{15}$ unique sequences, was hybridized to a complementary biotinylated capture oligonucleotide (CO; Supporting Figure S1). For the first two selection rounds, 1000 pmol of DNA library and 2000 pmol of capture oligonucleotide were annealed in 1000 µL of selection buffer (SB; Supporting Table S2) by heating to 95 °C for 5 min, followed by slow cooling to 4 °C and equilibration to room temperature. For subsequent rounds, the selection volume was reduced to 400 µL. The resulting duplex was incubated with pre-washed streptavidin-coated magnetic beads (Dynabeads M270 for rounds 1–2 and MyOne Streptavidin C1 beads for rounds 3–14; Thermo Fisher Scientific) for 1 h at room temperature with shaking at 1000 rpm. Beads were subsequently washed using the corresponding washing conditions described in Supporting Table S3.

Following immobilization, target-induced elution of MTX-binding sequences was performed by incubation with a free MTX solution under progressively increasing selection stringency. In the first round, 100 µM MTX in 1000 µL SB was incubated with the immobilized library for 2.5 h at room temperature with shaking at 1250 rpm. Eluted sequences were collected from the supernatant. Selection stringency progressively increased throughout the 14 selection rounds by reducing MTX concentration, increasing washing duration and temperature, introducing folic acid (FA) counter-selection, and decreasing the library-to-target ratio. Detailed selection parameters, including target and counter-target concentrations, washing conditions, and incubation times, are summarized in Supporting Table S3.

Eluted DNA pools were desalted using Illustra NAP-25 columns (Cytiva) and lyophilized prior to resuspension in Milli-Q water. Selected sequences were amplified by PCR using a FAM-labelled forward primer (FAM-Pr1) and a 5′-phosphorylated reverse primer (P-Pr2) (Supporting Table S1). PCR reactions (50 µL) contained 1× ThermoPol buffer, 1 µM primers, 200 µM dNTPs, and 25 U $mL^{-1}$ Taq DNA polymerase (New England Biolabs). Amplification conditions consisted of initial denaturation at 95 °C for 2 min, followed by 20 cycles of 95 °C for 15 s, 60 °C for 30 s, and 72 °C for 45 s, with final extension at 72 °C for 2 min. Multiple PCR reactions were pooled to obtain sufficient DNA for subsequent rounds.

PCR products were analyzed by 10% denaturing urea-PAGE using FAM-labelled standards and visualized with a ChemiDoc MP Imaging System (Bio-Rad). When sufficient amplification was achieved, dsDNA products were digested using homemade lambda exonuclease ($\lambda_{exo}$) in 1× Lambda Exonuclease buffer at 37 °C for 1 h to regenerate ssDNA libraries. Samples were subsequently purified using the NucleoSpin clean-up kit (Macherey-Nagel) and quantified by denaturing PAGE analysis with standards (a representative gel is shown in Figure S2). Enriched pools from rounds 9–14 were subjected to next-generation sequencing (NGS) analysis.

#### Next-Generation Sequencing and Data Analysis

For NGS analysis, enriched libraries from rounds 9–14 were amplified using NGS-specific primers containing Illumina-compatible overhangs (Supporting Table S1). PCR amplification was performed using Q5 Hot Start High-Fidelity DNA Polymerase (New England Biolabs) in 50 µL reactions containing 50 nM DNA template, 0.5 µM primers, and 200 µM dNTPs in 1× Q5 reaction buffer. Cycling conditions consisted of initial denaturation at 98 °C for 30 s, followed by 5 cycles of 98 °C for 10 s, 68 °C for 10 s, and 72 °C for 10 s, with final extension at 72 °C for 10 min. PCR products were purified by 3% agarose gel electrophoresis followed by extraction using the Wizard SV Gel and PCR Clean-Up System (Promega).

High-throughput sequencing was performed on an Illumina MiSeq platform using MiSeq Reagent Kit v2 (300-cycle format) at the Central Analytical Research Facility (QUT). Sequencing data were analyzed using AptaSUITE software to identify enriched aptamer families based on sequence abundance, conserved motifs, and clustering analysis. Representative sequences from the top ten enriched families were selected for further biochemical and biophysical characterization.

**Prediction of aptamer structure.**

The ten most enriched candidate aptamer sequences identified by NGS analysis (Supporting Table S4) were synthesized as FAM-labelled oligonucleotides and screened for MTX binding using isothermal titration calorimetry (ITC). Among these candidates, aptamers 3315 and 930 demonstrated positive target binding and were selected for further characterization. Secondary structures of aptamers 3315 and 930, together with previously reported MTX aptamers HMX38, MTX40, and MSmt7, as well as their truncated and mutated variants, were predicted using the Mfold and NUPACK online servers under standard DNA folding conditions (100 mM $Na^+$ and 2 mM $Mg^{2+}$ at 22°C).[5,6]

**Identification of G-Quadruplex Formation Probability**

The probability of G-quadruplex (G4) structure formation in aptamers 3315 and 930 was evaluated using the G4-specific nanobody SG4 and previously established Direct- and FLAG-ELISA assays.[7–9] For FLAG-ELISA experiments, Corning 96-well half-area microplates (Cat. No. 3690) were coated with 100 µL of 5 µg $mL^{-1}$ streptavidin in PBS and incubated overnight on ice with gentle rotation. Wells were washed five times with PBST and blocked with PBSTB for 1 h at room temperature under gentle agitation. Following washing, 100 µL of 500 nM renatured biotinylated aptamers (BioTEG-3315 and BioTEG-930) or control oligonucleotides (Bio-MycG4, Bio-mMycG4, and BioTEG-17-43) were added and incubated for 1 h at room temperature. Serial dilutions of FLAG-tagged SG4 nanobody were prepared from a 1000 nM stock solution, with 50 µL added per well followed by incubation for an additional 1 h. After extensive washing, 50 µL of anti-FLAG M2-HRP antibody (1:5000 dilution; Sigma-Aldrich, Cat. No. A8592) was added and incubated for 15 min at room temperature with gentle rotation. Wells were washed extensively before the addition of 50 µL of 1-Step Ultra TMB ELISA substrate (Thermo Scientific Pierce). Color development was allowed to proceed for 15 min in the dark, and the reactions were stopped with 50 µL of 2 M $H_2SO_4$. Absorbance was recorded at 450 nm with background correction at 540 nm using a BioTek Synergy Neo2 plate reader.

For Direct-ELISA experiments, plates were coated overnight on ice with 100 µL of 5 µg $mL^{-1}$ SG4 nanobody in PBS under gentle rotation. Wells were washed five times with PBST and blocked with PBSTB for 1 h at room temperature. Serial dilutions prepared from 3000 nM renatured FAM-labelled aptamers and control oligonucleotides were added (50 µL per well) and incubated for 1 h at room temperature. Following washing, wells were incubated with 50 µL of anti-FITC HRP-conjugated antibody (1:5000 dilution; Invitrogen, Cat. No. PA1-86159) for 1 h at room temperature. After extensive washing, 50 µL of 1-Step Ultra TMB ELISA substrate was added, and the plate was incubated in the dark prior to termination of the reaction with 50 µL of 2 M $H_2SO_4$. Absorbance at 450 nm with background correction at 540 nm was recorded using a BioTek Synergy Neo2 plate reader as described above.

**Molecular dynamics simulations and *in silico* prediction of MTX-binding sites**

*In silico* modeling was performed to obtain mechanistic insight into the structural basis of MTX recognition by aptamers 3315-53 and 930-65.[10–12] Initial secondary structures were predicted using the Mfold web server under DNA folding conditions at 37 °C, with 100 mM $Na^+$ and 1 mM $Mg^{2+}$.[6] Secondary-structure models within 5% of the minimum free energy were used as input for three-dimensional structure generation using the 3dRNA/DNA server.[13] The top-ranked 3D models were selected for molecular dynamics refinement.

Aptamer structures were solvated in an explicit TIP3P water box using the LEaP module of AMBER20,[14,15] with a minimum distance of 12 Å between the solute and the box edge. $Na^+$ and $Cl^-$ ions were added to achieve an ionic strength of 0.15 M using the SPLIT ion-placement approach.[16] DNA systems were described using the parmbsc1 force field.[17] Each system was subjected to sequential energy minimization, including minimization of water molecules, hydrogen atoms, DNA atoms, and finally the complete system. The systems were then heated from 0 to 310.15 K over 50 ps, followed by a 50 ps density equilibration. Production simulations were performed for 500 ns under NPT conditions at 310.15 K and 1 bar using a Langevin thermostat and Berendsen barostat. Hydrogen-containing bonds were constrained using the SHAKE algorithm with a 2 fs time step.[18–20] Trajectories were analyzed using CPPTRAJ,[21] and representative dominant conformations were selected by clustering analysis.

Putative MTX-binding pockets were identified from the refined aptamer conformations using Fpocket[22] and DoGSite3.[23,24] MTX docking was performed using GOLD software.[25] For each predicted binding pocket, the docking search space was defined using the selected active nucleotides with a 30 Å radius, and 500 genetic algorithm runs were performed using the GOLD score fitness function. The most favorable docking poses were used as starting structures for 300 ns MD simulations of the MTX–aptamer complexes (Supporting Table S6). Complexes in which MTX remained stably associated with the predicted binding pocket throughout the simulation were further analyzed using MM-GBSA.[26,27] Binding free energies were calculated using MM-PBSA.py in AMBER20 from 200 frames extracted from the final 100 ns of each trajectory.[28] Per-nucleotide energy decomposition was performed to identify residues contributing most strongly to MTX binding.

The binding free energy (eq 1) is calculated by the difference in variation of total free energy between the bound and unbound states:

$$\Delta G_{Bind} = \Delta G_{MTX:Aptamer} - (\Delta G_{MTX} + \Delta G_{Aptamer}) \quad (1)$$

The complex is considered thermodynamically stable when $\Delta G_{Bind}$ is negative, which means that the sum of $\Delta G$ of the aptamer and MTX is higher than the $\Delta G$ of the complex. All data are summarized in Table S7.

### Supporting Tables

#### Table S1. Oligonucleotide sequences used in this study.

| Name | Sequence, 5´ → 3´ |
|---|---|
| **1. Library and primers used for Capture-SELEX:** | |
| **L45** | /56-FAM/GGAGGCTCTCGGGACGACNNNNNNNNNNNNNNNNNNNNNNNNNNNNNNNNNNNNNNNNNNNNNGTCGTCCCGCCTTTAGGATTTACAG |
| **F-Pr1** | /56-FAM/GGAGGCTCTCGGGACGAC |
| **P-Pr2** | **/5Phos/**CTGTAAATCCTAAAGGCGGGACGAC |
| **CO** | GTCGTCCCGAGAGCCATA**/3BioTEG/** |
| **NGS-Pr1** | TCGTCGGCAGCGTCAGATGTGTATAAGAGACAGGGAGGCTCTCGGGACGAC |
| **NGS-Pr2** | GTCTCGTGGGCTCGGAGATGTGTATAAGAGACAGCTGTAAATCCTAAAGGCGGGACGAC |
| **2. Selected aptamers:** | |
| **121** | GGAGGCTCTCGGGACGACCACGGAGTGGGCGGGTAGTTCTGTCTTCCCCCTCCAACCCGGCCAGTCGTCCCGCCTTTAGGATTTACAG |
| **339** | GGAGGCTCTCGGGACGACGCGAGCGTTGCGAGCGAGTGCGAAGCTCCATGCCTTCGTGTTGCGGTCGTCCCGCCTTTAGGATTTACAG |
| **318** | GGAGGCTCTCGGGACGACCACGCAGTGCCCATAATGCTGCCCAGTTCAGCGCTTTTGCGGTCCGTCGTCCCGCCTTTAGGATTTACAG |
| **524** | GGAGGCTCTCGGGACGACCACGCGTAAGCGTCAAGTGTCGCACCCACTTCTCTTACCGGGTGCGTCGTCCCGCCTTTAGGATTTACAG |
| **597** | GGAGGCTCTCGGGACGACCACAGCGCACTTCAGCGATGTGGTTGATCTCGTTGTCAACGTTGCGTCGTCCCGCCTTTAGGATTTACAG |
| **3315** | GGAGGCTCTCGGGACGACGGGGGGTGTCGCGTCGTATGTTTTACATCCCGATGTTGTTGTCTAGTCGTCCCGCCTTTAGGATTTACAG |
| **1312** | GGAGGCTCTCGGGACGACCAGGAGCGAGAGCCACGCGATCGTTAACAGTTGATCTGCTTCGCAGTCGTCCCGCCTTTAGGATTTACAG |
| **930** | GGAGGCTCTCGGGACGACCAGTGCTGTTCCATCGACTTGTCGTCCAATAGATGGTTCATTCGTGTCGTCCCGCCTTTAGGATTTACAG |
| **1896** | GGAGGCTCTCGGGACGACCAGGACCGAGGCCGGATGTGTGTGTCACACAAGCGGTCCTCCCGTGTCGTCCCGCCTTTAGGATTTACAG |
| **2542** | GGAGGCTCTCGGGACGACGACACCACATGCACATTGTGGGGTCGCCCTGCTTCACGTCTGCTTGTCGTCCCGCCTTTAGGATTTACAG |
| **BioTEG-3315-88** | **/5BiotinTEG/**GGAGGCTCTCGGGACGACGGGGGGTGTCGCGTCGTATGTTTTACATCCCGATGTTGTTGTCTAGTCGTCCCGCCTTTAGGATTTACAG |
| **BioTEG-930-88** | **/5BiotinTEG/**GGAGGCTCTCGGGACGACCAGTGCTGTTCCATCGACTTGTCGTCCAATAGATGGTTCATTCGTGTCGTCCCGCCTTTAGGATTTACAG |
| **3. Truncated variants** | |
| **930-45** | CAGTGCTGTTCCATCGACTTGTCGTCCAATAGATGGTTCATTCGT |
| **930-53** | CG**G**CCAGTGCTGTTCCATCGACTTGTCGTCCAATAGATGGTTCATTCGTG**C**CG |
| **930-63** | CGGGACGACCAGTGCTGTTCCATCGACTTGTCGTCCAATAGATGGTTCATTCGTGTCGTCCCG |
| **930-65** | **C**CGGGACGACCAGTGCTGTTCCATCGACTTGTCGTCCAATAGATGGTTCATTCGTGTCGTCCCG**G** |
| **930-67** | **GG**CGGGACGACCAGTGCTGTTCCATCGACTTGTCGTCCAATAGATGGTTCATTCGTGTCGTCCCGCC |
| **930-71** | CGGGACGACCAGTGCTGTTCCATCGACTTGTCGTCCAATAGATGGTTCATTCGTGTCGTCCCGCCTTTAGG |
| **3315-38** | CG**G**CGGGGGGTGTCGCGTCGTATGTTTTACATCCCGAT |
| **3315-39** | **C**CG**G**CGGGGGGTGTCGCGTCGTATGTTTTACATCCCGAT |
| **3315-39a** | ACGACGGGGGGTGTCGCGTCGTATGTTTTACATCCCGAT |
| **3315-45a** | **C**CG**G**CGGGGGGTGTCGCGTCGTATGTTTTACATCCCGATAGCCGG |
| **3315-45** | GGGGGGTGTCGCGTCGTATGTTTTACATCCCGATGTTGTTGTCTA |
| **3315-47** | ACGACGGGGGGTGTCGCGTCGTATGTTTTACATCCCGATGTTGTCGT |

**3315-50** AATACGACGGGGGGTGTCGCGTCGTATGTTTTACATCCCGATGTTGTTGT

**3315-52** GGAGGCTCTCGGGACGACGGGGGGTGTCGCGTCGTATGTTTTACATCCCGAT

**3315-53** CGGCGGGGGGTGTCGCGTCGTATGTTTTACATCCCGATGTTGTTGTCTAGCCG

**3315-55** CCGGCGGGGGGTGTCGCGTCGTATGTTTTACATCCCGATGTTGTTGTCTAGCCGG

**3315-55a** ACGACGGGGGGTGTCGCGTCGTATGTTTTACATCCCGATGTTGTTGTCTAGTCGT

**3315-57** GCCGGCGGGGGGTGTCGCGTCGTATGTTTTACATCCCGATGTTGTTGTCTAGCCGGC

**3315-60** ATACGACGGGGGGTGTCGCGTCGTATGTTTTACATCCCGATGTTGTTGTCTAGTCGTCCC

**3315-60a** AGGACGACGGGGGGTGTCGCGTCGTATGTTTTACATCCCGATGTTGTTGTCTAGTCGTCC

**3315-61** GGGACGACGGGGGGTGTCGCGTCGTATGTTTTACATCCCGATGTTGTTGTCTAGTCGTCCC

**3315-63** CGGGACGACGGGGGGTGTCGCGTCGTATGTTTTACATCCCGATGTTGTTGTCTAGTCGTCCCG

**930-65-BioTEG** CCGGGACGACCAGTGCTGTTCCATCGACTTGTCGTCCAATAGATGGTTCATTCGTGTCGTCCCGG/3BioTEG/

**3315-53-BioTEG** CGGCGGGGGGTGTCGCGTCGTATGTTTTACATCCCGATGTTGTTGTCTAGCCG/3BioTEG/

### 4. Aptamer mutants:

**3315-88-G7C** GGAGGCTCTCGGGACGACGGCGGGTGTCGCGTCGTATGTTTTACATCCCGATGTTGTTGTCTAGTCGTCCCGCCTTTAGGATTTACAG

**3315-88-G8C** GGAGGCTCTCGGGACGACGGGCGGTGTCGCGTCGTATGTTTTACATCCCGATGTTGTTGTCTAGTCGTCCCGCCTTTAGGATTTACAG

**3315-88-G9C** GGAGGCTCTCGGGACGACGGGGCGTGTCGCGTCGTATGTTTTACATCCCGATGTTGTTGTCTAGTCGTCCCGCCTTTAGGATTTACAG

**3315-88-A22T** GGAGGCTCTCGGGACGACGGGGGGTGTCGCGTCGTTTGTTTTACATCCCGATGTTGTTGTCTAGTCGTCCCGCCTTTAGGATTTACAG

**3315-88-G24C** GGAGGCTCTCGGGACGACGGGGGGTGTCGCGTCGTATCTTTTACATCCCGATGTTGTTGTCTAGTCGTCCCGCCTTTAGGATTTACAG

**3315-88-T26A** GGAGGCTCTCGGGACGACGGGGGGTGTCGCGTCGTATGTATTACATCCCGATGTTGTTGTCTAGTCGTCCCGCCTTTAGGATTTACAG

**3315-88-C34A** GGAGGCTCTCGGGACGACGGGGGGTGTCGCGTCGTATGTTTTACATCACGATGTTGTTGTCTAGTCGTCCCGCCTTTAGGATTTACAG

**3315-53a** CGACGGGGGGTGTCGCGTCGTATGTTTTACATCCCGATGTTGTTGTCTAGTCG

**3315-53b** CGACGGGGGGTGCCGCGTCGTATGTTTTACATCCCGATGTTGTTGTCTAGTCG

**3315-53c** CGACGCGGGGTGCCGCGTCGTATGTTTTACATCCCGATGTTGTTGTCTAGTCG

**3315-54** ACGACGCGGGGTGCCGCGTCGTATGTTTTACATCCCGATGTTGTTGTCTAGTCG

**3315-54a** ACGGCGGGGGGTGTCGCGTCGTATGTTTTACATCCCGATGTTGTTGTCTAGCCG

**3315-53-T28G** CGGCGGGGGGTGTCGCGTCGTATGTTTGACATCCCGATGTTGTTGTCTAGCCG

**3315-53-A29C** CGGCGGGGGGTGTCGCGTCGTATGTTTTCCATCCCGATGTTGTTGTCTAGCCG

**3315-53-T41G** CGGCGGGGGGTGTCGCGTCGTATGTTTTACATCCCGATGTGGTTGTCTAGCCG

**3315-53-G42T** CGGCGGGGGGTGTCGCGTCGTATGTTTTACATCCCGATGTTTTTGTCTAGCCG

**3315-53-A49C** CGGCGGGGGGTGTCGCGTCGTATGTTTTACATCCCGATGTTGTTGTCTCGCCG

**3315-53-C52A** CGGCGGGGGGTGTCGCGTCGTATGTTTTACATCCCGATGTTGTTGTCTAGCAG

**930-65-A12C** CCGGGACGACCCGTGCTGTTCCATCGACTTGTCGTCCAATAGATGGTTCATTCGTGTCGTCCCGG

**930-65-G13T** CCGGGACGACCATTGCTGTTCCATCGACTTGTCGTCCAATAGATGGTTCATTCGTGTCGTCCCGG

**930-65-A23C** CCGGGACGACCAGTGCTGTTCCCTCGACTTGTCGTCCAATAGATGGTTCATTCGTGTCGTCCCGG

**930-65-T24G** CCGGGACGACCAGTGCTGTTCCAGCGACTTGTCGTCCAATAGATGGTTCATTCGTGTCGTCCCGG

**930-65-G42T** CCGGGACGACCAGTGCTGTTCCATCGACTTGTCGTCCAATATATGGTTCATTCGTGTCGTCCCGG

**930-65-T44G** CCGGGACGACCAGTGCTGTTCCATCGACTTGTCGTCCAATAGAGGGTTCATTCGTGTCGTCCCGG

**930-65-G54T** CCGGGACGACCAGTGCTGTTCCATCGACTTGTCGTCCAATAGATGGTTCATTCTTGTCGTCCCGG

**930-65-T55G** CCGGGACGACCAGTGCTGTTCCATCGACTTGTCGTCCAATAGATGGTTCATTCGGGTCGTCCCGG

**HMX38-M1** CTCTCGCGCGAACGCGGGATGTTTGGGGGACCCACGTTCGCCC

**HMX38-M2** CTCTCGGGCGAACGCGCGATGTTTGGGGGACCCACGTTCGCCC

| | |
|---|---|
| **HMX38-M3** | CTCTCGGGCGAACGCGGGATGTTTGG**C**GGACCCACGTTCGCCC |
| **HMX38-M4** | CTCTCGGGCGAACGCGGGATGTTTGGG**C**GACCCACGTTCGCCC |
| **HMX38-M5** | CTCTCGGGCGAACGCGGG**T**TGTTTGGGGGACCCACGTTCGCCC |
| **HMX38-M6** | CTCTCGGGCGAACGCGGGAT**C**TTTGGGGGACCCACGTTCGCCC |
| **HMX38-M7** | CTCTCGGGCGAACGCGGGATGTTTGGGG**C**ACCCACGTTCGCCC |
| **HMX38-M8** | CTCTCGGGCGAACGCGGGATGTTTGGGGGAC**A**CACGTTCGCCC |
| **3315-53-SCR** | TACATAAGCCGGGTGAGGCTCCTTCGTTTTCGCTTCCGCGGTTGTGGGGTGTG |
| **930-65-SCR** | ACTCGCCTGTGTACACGCCGCGGTGTTATCTGGTCTTATGACGTTACGTGCTCCGACTACGAG |

### 5. Fluorescent sensors

| | |
|---|---|
| **FAM-3315-88** | **/56-FAM/** GGAGGCTCTCGGGACGACGGGGGGTGTCGCGTCGTATGTTTTACATCCCGATGTTGTTGTCTAGTCGTCCCGCCTTTAGGATTTACAG |
| **FAM-930-88** | **/56-FAM/** GGAGGCTCTCGGGACGACCAGTGCTGTTCCATCGACTTGTCGTCCAATAGATGGTTCATTCGTGTCGTCCCGCCTTTAGGATTTACAG |
| **Q-Iowa** | GGTCGTCCCGAGAG**/3IABkFQ/** |
| **FAM-HMX38** | **/56-FAM/**CTCTCGGGCGAACGCGGGATGTTTGGGGGACCCACGTTCGCCC |
| **HMX38-Q** | CGTTCGCCCGAGAG**/3IABkFQ/** |

### 6. Aptamers for electrochemical sensors

| | |
|---|---|
| **S-3315-55-MB** | **/5ThioMC6-D/C**CCG**G**CGGGGGGTGTCGCGTCGTATGTTTTACATCCCGATGTTGTTGTCTAG**C**CG**G/3MeBlN/** |
| **S-930-65-MB** | **/5ThioMC6-D/** **C**CGGGACGACCAGTGCTGTTCCATCGACTTGTCGTCCAATAGATGGTTCATTCGTGTCGTCCCG**G/3MeBlN/** |

### 7. Published aptamer and sensors:

| | |
|---|---|
| **HMX38** | CTCTCGGGCGAACGCGGGATGTTTGGGGGACCCACGTTCGCCC |
| **MTX40** | ACCATTCTGCAACTTTGTGCTAGTCTTGGCCTTTGGTGGT |
| **MSmt7** | CGAGCATAGGCAGAACTTACCCGGTGCTATATCTCGGTGTATTTCGAAGCCCTGTCGTAAGAGCGAGTCATTC |

### 8. Control sequences:

| | |
|---|---|
| **MycG4** | TGAGGGTGGGTAGGGTGGGTAA |
| **mMycG4** | TGAGTGTGCGTAGTGTGTGTAA |
| **17-43** | ACTCGTCTCTTGGGGTTTGGCGTCGGGCCTGGCGTGGACGAGT |

DNA sequences are written using Integrated DNA Technologies (IDT) modification codes: /56-FAM/ and /5Phos/ represent 6-carboxyfluorescein (FAM) fluorescent dye and 5'-phosphorylation, respectively; /5Biosg/ indicates a biotin tag with a short linker; /5BiotinTEG/ and /3BiotinTEG/ indicate biotin tags attached via a TEG linker at the 5' and 3' ends, respectively. Forward primer sequences are highlighted in green, reverse primers in blue, and mutations are indicated in bold red. Poly-T and poly-A linkers used in dimeric constructs are shown in italicized pink.

### Table S2. Buffer and solutions used in this study

| Buffer | Composition |
|---|---|
| **SB** | 40 mM Tris-HCl, 100 mM NaCl, 2 mM $MgCl_2$, 1 mM $CaCl_2$, pH 7.4 |
| **SB-HS (high-salt buffer)** | 20 mM Tris-HCl, 250 mM NaCl, 5 mM $MgCl_2$, 1 mM $CaCl_2$, pH 7.4 |
| **WB1** | SB with 0.05% Tween-20 and 0.1% BSA |
| **WB2** | WB1 with 0.05% Milk |
| **WB3** | SB-HS with 0.05% Tween-20, 0.1% BSA, 0.1% Milk, and 20 mM Glucose |
| **WB4** | WB3 with 10% heat-inactivated human serum |
| **EB** | 50 mM EDTA in 95% formamide |
| **SB-Mg (SB without $Mg^{2+}$)** | 40 mM Tris-HCl, 100 mM NaCl, 1 mM $CaCl_2$, pH 7.4 |
| **SB-LS (low salt buffer)** | 20 mM Tris-HCl, 20 mM NaCl, 0.5 mM $MgCl_2$, 1 mM $CaCl_2$, pH 7.4 |
| **PBS** | 10 mM phosphate buffer, 2.7 mM KCl, 137 mM NaCl, pH 7.4 |
| **PBSM** | PBS with 5 mM $MgCl_2$ |
| **PBST** | PBS with 0.05% Tween-20 and 0.1% BSA |
| **PBSTB** | PBST with 5% BSA |
| **B1** | 50 mM Tris (pH 7.9), 100 mM NaCl, 10 mM $MgCl_2$ |
| **B2** | 50 mM Tris (pH 7.9), 100 mM NaCl, 10 mM $MgCl_2$, 100 μg $mL^{-1}$ recombinant albumin |
| **2× gel loading buffer** | 95% formamide, 50 mM EDTA, 0.05% bromophenol blue |
| **1× ThermoPol Buffer purchased from NEB** | 20 mM Tris-HCl, 10 mM $(NH_4)_2SO_4$, 10 mM KCl, 2 mM $MgSO_4$, 0.1% TritonX-100, pH 8.8 @ 25 °C |
| **1× Lambda Exonuclease Buffer** | 67 mM Glycine-KOH (pH 9.4), 2.5 mM $MgCl_2$, 0.01% Triton X-100 |
| **10% Urea-PAGE** | 10% Acrylamide: bisacrylamide (19:1), 7 M Urea in 1× TBE (89 mM Tris-borate, 89 mM boric acid, 2 mM EDTA, and pH 8.3) |

### Table S3. Capture-SELEX strategy against MTX[a]

| SELEX Round | Ratio Library:MTX:FA | Library | | Counter target (FA) | | | | Target (MTX) | | | Washing | | | |
|---|---|---|---|---|---|---|---|---|---|---|---|---|---|---|
| | | V, µL | pmol | pmol | C, µM | Times | Time, min | pmol | C, µM | Time, min | Buffer[b] | Times | Time, min | T,°C |
| R1 | 1:100:0 | 1000 | 1000 | | | | | 100,000 | 100 | 150 | SB | 3 | 1 | RT |
| R2 | 1:142:0 | 1000 | 705 | | | | | 100,000 | 100 | 60 | WB1 | 2 | 10 | 30 |
| R3 | 1:50:0 | 400 | 1000 | | | | | 50,000 | 125 | 30+5 | WB1 | 2 | 15 | 30 |
| R4 | 1:20:0 | 400 | 1000 | | | | | 20,000 | 50 | 30+5 | WB2 | 3 | 15 | 30 |
| R5 | 1:10:0 | 400 | 1000 | | | | | 10,000 | 25 | 30+5 | WB2 | 3 | 20 | 30 |
| R6 | 1:5:0 | 400 | 1000 | | | | | 5,000 | 12.5 | 30+5 | WB3 | 3 | 30 | 30 |
| R7 | 1:3:0 | 400 | 700 | | | | | 2,000 | 5 | 30+5 | WB3 | 3 | 30 | 30 |
| R8 | 1:1.4:0.1 | 400 | 700 | 100 | 0.25 | 1 | 10 | 1,000 | 2.5 | 30+5 | WB3 | 3 | 30 | 30 |
| R9 | 1:0.5:0.5 | 400 | 1000 | 500 | 1.25 | 1 | 10 | 500 | 1.25 | 15+5 | WB3 | 3 | 30 | 30 |
| R10 | 1:0.2:1 | 400 | 1000 | 1000 | 2.5 | 2 | 15 | 200 | 0.5 | 15+5 | WB3 | 3 | 40 | 30 |
| R11 | 1:0.2:1 | 400 | 1000 | 1000 | 2.5 | 2 | 15 | 200 | 0.5 | 15+5 | WB4+WB3 | 1+2 | 10+40 | 30 |
| R12 | 1:0.2:1 | 400 | 1000 | 1000 | 2.5 | 2 | 15 | 200 | 0.5 | 15+5 | WB4+WB3 | 1+2 | 40+40 | 30 |
| R13 | 1:0.1:1 | 400 | 1000 | 1000 | 2.5 | 4 | 15 | 100 | 0.25 | 15+5 | WB4+WB3 | 1+5 | 30+60/30/5 | 20/30 |
| R14 | 1:0.2:1.6 | 400 | 600 | 1000 | 2.5 | 5 | 15 | 100 | 0.25 | 15+5 | WB4+WB3 | 1+5 | 30+60/30/5 | 20/30 |

[a]For the first two rounds of selection, Streptavidin Dynabeads M270 were used, and Streptavidin MyOne C1 beads were used for the following rounds, both at a concentration of 10 µg mL$^{-1}$

[b]Refer to Table S2 for the description of buffer composition.

**Table S4. NGS sequencing results**

| | ID | Sequence of random region, 5´→3´ | Count the number in each round | | | | | | | Enrichment |
|---|---|---|---|---|---|---|---|---|---|---|
| | | | R9 | R10 | R11 | R12 | R13 | R14 | Total | R14/R9 |
| 1 | 121 | CACGGAGTGGGCGGGTAGTTCTGTCTTCCCCCTCCAACCCGGCCA | 10 | 49 | 5 | 562 | 2245 | 3721 | 6592 | 372 |
| 2 | 339 | GCGAGCGTTGCGAGCGAGTGCGAAGCTCCATGCCTTCGTGTTGCG | 13 | 35 | 1 | 454 | 860 | 1284 | 2647 | 99 |
| 3 | 318 | CACGCAGTGCCCATAATGCTGCCCAGTTCAGCGCTTTTGCGGTCC | 33 | 195 | 2 | 722 | 1188 | 1061 | 3201 | 32 |
| 4 | 524 | CACGCGTAAGCGTCAAGTGTCGCACCCACTTCTCTTACCGGGTGC | 1 | 5 | 0 | 96 | 115 | 499 | 716 | 499 |
| 5 | 597 | CACAGCGCACTTCAGCGATGTGGTTGATCTCGTTGTCAACGTTGC | 14 | 45 | 0 | 222 | 317 | 473 | 1071 | 34 |
| 6 | 3315 | GGGGGGTGTCGCGTCGTATGTTTTACATCCCGATGTTGTTGTCTA | 0 | 3 | 0 | 50 | 300 | 439 | 792 | >439 |
| 7 | 1312 | CAGGAGCGAGAGCCACGCGATCGTTAACAGTTGATCTGCTTCGCA | 15 | 39 | 1 | 269 | 495 | 356 | 1175 | 24 |
| 8 | 930 | CAGTGCTGTTCCATCGACTTGTCGTCCAATAGATGGTTCATTCGT | 34 | 145 | 0 | 195 | 425 | 356 | 1155 | 10 |
| 9 | 1896 | CAGGACCGAGGCCGGATGTGTGTGTCACACAAGCGGTCCTCCCGT | 1 | 8 | 0 | 50 | 130 | 355 | 544 | 355 |
| 10 | 2542 | GACACCACATGCACATTGTGGGGTCGCCCTGCTTCACGTCTGCTT | 4 | 23 | 0 | 88 | 154 | 244 | 513 | 61 |

**Table S5. ITC-derived thermodynamic parameters for MTX binding to full-length aptamers**

| Aptamer | $K_D$, nM | $\Delta H$, kcal mol$^{-1}$ | $-T\Delta S$, kcal mol$^{-1}$ | $\Delta G$, kcal mol$^{-1}$ | $N$, sites |
|---|---|---|---|---|---|
| **3315-88** | 88.7 ± 26.6 | -24.0 ± 0.821 | 14.5 | -9.47 | 0.466 ± 0.0095 |
| **930-88** | 408 ± 115 | -26.7 ± 1.49 | 18.2 | -8.57 | 0.607 ± 0.019 |
| **HMX38** | 291 ± 58.4 | -23.6 ± 0.809 | 14.8 | -8.77 | 0.625 ± 0.012 |
| **MTX40** | 1830 ± 419 | -29.3 ± 2.63 | 21.5 | -7.70 | 0.636 ± 0.024 |

ITC measurements were performed at 20 °C in selection buffer (SB: 40 mM Tris-HCl, 100 mM NaCl, 2 mM $MgCl_2$, 1 mM $CaCl_2$, pH 7.4), using 10 µM aptamer in the cell and 100 µM MTX in the syringe. Values were obtained by fitting the integrated binding isotherms to a single-site binding model after correction for heats of dilution. $K_D$, ΔH, and N are reported as fitted values ± fitting error. ΔG was calculated from $\Delta G = RT \ln K_D$, and −TΔS was calculated from ΔG − ΔH. Stoichiometry values below 1 likely reflect the fraction of binding-competent folded aptamer under the assay conditions and/or uncertainty associated with fitting low-c-value ITC experiments for high-affinity nucleic acid–small molecule interactions. Full isotherms and corresponding fits are provided in Figure S7.

**Table S6. GOLD Score obtained by docking MTX to 3315-53 and 930-65 aptamers.**

| Aptamer | Docking GOLD Score obtained with MTX | | |
|---|---|---|---|
| **3315-53** | 82.02 | | |
| **930-65** | **Pocket 1** | **Pocket 2** | **Pocket 3** |
| | 62.78 | 68.70 | 72.79 |

**Table S7. Results obtained from MM-GBSA calculations of 3315-53 and 930-65 aptamers with MTX**

| Aptamer ID | Van der Waals energy (kcal mol$^{-1}$) | Electrostatic energy (kcal mol$^{-1}$) | EGB (kcal mol$^{-1}$) | ESURF (kcal mol$^{-1}$) | Nonpolar contribution (kcal mol$^{-1}$) | Polar contribution (kcal mol$^{-1}$) | ΔG Binding Free Energy (kcal mol$^{-1}$) |
|---|---|---|---|---|---|---|---|
| **3315-53** | -61.41 | 1914.59 | -1857.51 | -5.74 | -67.1 | 57.1 | -10.0±4.3 |
| **930-65** | -54.52 | 2182.17 | -2127.17 | -4.79 | -59.3 | 54.9 | -4.4±3.3 |

#### Table S8. Currently available MTX-specific aptamer-based biosensors

| Aptamer | Optimized aptamer sequence, 5'→3' | Size, nt | Selection method | $K_D$ | Binding measurement | Buffer conditions | Sensor format | LOD | Linear range | Specificity | Clinical human samples | *In vivo* monitoring | Ref. |
|---|---|---|---|---|---|---|---|---|---|---|---|---|---|
| **HMX38** | GGGCGAACGCGGGATGTTTGGGGGACCCACGTTCGCCC | 38 | Capture-SELEX (streptavidin column), 18 rounds, N30 | 165 ± 82.5 nM in PBS with 10 mM $MgCl_2$ | ITC | PBS | Fluorescent structure-switching aptasensor; Direct; endpoint/offsite | 30 nM in buffer; 180 nM in 50% fetal bovine serum | 0.1–2 µM (buffer); 0.5–10 µM (50% serum) | | No | No | [29] |
| **MSmt7** | CGAGCATAGGCAGAACTTACGATAGCTAGGGTGTGTTCTTGTGTGTTGTGGATGTCGTAAGAGCGAGTCATTC | 73 | †MACE-SELEX, MTX-immobilised, 3 rounds N33 | 0.57 µM in buffer 50 mM HEPES [pH 7.4], 100 mM NaCl, and 5 mM MgCl2 | ITC | 20 mM Tris-HCl pH 7.4, 10 mM NaCl, 1 mM MgCl2, and 0.01% Tween 20 | ‡Competitive ELISA-like assay; Indirect/competitive; endpoint/offsite | 118 nM in 50% human serum; 0.029 nM after RCA | Unspecified | Binds to DAMPA (MTX 0, 0.53 µM), no FA | No | No | [30] |
| **MTX40 stem-4** | ACCATTCTGCAACTTTGTGCTAGTCTTGGCCTTTGGTGGT | 40 | Capture-SELEX (streptavidin column) N36 | 1 µM in buffer, 27 µM in buffer, 1.05 ± 0.09 mM in whole blood at 37°C | MTX fluorescence quenching, ThT, EAB | PBS, whole blood, live rat plasma | EAB, implantable; Direct; Real-time; on-site | Unspecified | Unspecified | Unspecified | No | Yes (rats) | [31,32] |

†*Microbead-assisted capillary electrophoresis (MACE)-SELEX*
‡*Hemin-DNAzyme colorimetric sensor; RCA-amplified assay*

### Table S9. Comparison of available methods for MTX detection

| Sensor type | Equipment | Detection method | Linear range in matrix | LOD in matrix | Time/labor | Cost | Ref |
|---|---|---|---|---|---|---|---|
| Competitive ELISA | ELISA reader | Antibody-based colorimetric immunoassay | 0.0003–2.2 μM | 0.19 nM | •• | ■■ | Abcam ELISA kit |
| Homogeneous immunoassay | Roche Cobas C systems | Competitive homogeneous enzyme immunoassay | 0.04–1.2 μM | 35 nM | •• | ■■ | Roche ONLINE TDM MTX |
| Homogeneous immunoassay | Automated clinical chemistry analyzer | ARK Methotrexate II; homogeneous enzyme immunoassay | 0.03–1.3 μM | 30 nM | •• | ■■ | ARK Methotrexate II |
| FPIA | Abbott TDx/FLx | Fluorescence polarization immunoassay | 0.03–1 μM | 30 nM | •• | ■■ | [33] |
| FPIA | Abbott TDx/FLx | Fluorescence polarization immunoassay | 0.05–810 μM | NR | •• | ■■ | [34] |
| EMIT | Siemens Dimension / clinical chemistry analyzer | Syva EMIT Methotrexate | 0.3–2 μM | 300 nM | •• | ■■ | commercial/clinical method [35] |
| EMIT | Siemens VIVA-E / clinical chemistry analyzer | Modified enzyme-multiplied immunoassay technique | 0.05 μM–1 μM | 50 nM | •• | ■■ | [36] |
| CMIA | Abbott Architect i1000 | Chemiluminescent microparticle immunoassay | 0.04–1.5 μM | 40 nM | •• | ■■ | [33] |
| HPLC | HPLC-UV | Chromatographic separation with UV detection | 0.05–5 μM | 3 nM | ••• | ■■■ | [37] |
| HPLC | HPLC-UV | Chromatographic separation with UV detection | 0.1–5 μM | 10 nM | ••/••• | ■■■ | [38] |
| LC/MS-MS | LC/MS-MS | Tandem mass spectrometry | 0.01–1 μM | 3 nM | ••• | ■■■ | [39] |
| LC/MS-MS | LC/MS-MS | Tandem mass spectrometry | 0.01–25 μM | 10 nM | ••• | ■■■ | [40] |
| UHPLC–MS/MS | UHPLC–MS/MS | Tandem mass spectrometry | 0.002–2 μM | 2 nM | ••• | ■■■ | [41] |
| SPR | SPR | Surface plasmon resonance immunosensor | 0.028–0.5 μM | 28 nM | • | ■■ | [42] |
| PEC | Implantable PEC device | Photoelectrochemical detection using $Fe_1$–$Zn_1$–$TiO_2$ photoelectrode | 1–100 μM in vivo, mice | 684 nM | ○ | ■ | 43 |
| Aptasensor | Plate reader | Aptamer-based, Fluorescence | 0.5–10 μM in 50% serum | 180 nM | •• | ■■ | [29] |
| EAB | *In vivo*, an electrochemical reader | Aptamer-based, Electrochemical | 10– 900 μM | 10,000 nM | ○ | ■ | [44] |
| EAB/aptasensor | Potentiostat/gold electrodes | Aptamer-based, electrochemical | 100–1000 μM | 100 μM | • | ■ | Current assay |

FPIA, fluorescence polarization immunoassay; EMIT, enzyme-multiplied immunoassay technique; PEC, photoelectrochemical; EAB, electrochemical aptamer-based sensor; SPR, surface plasmon resonance; NR, not reported.

**Time/labor:**

••• - centralized, labor-intensive workflow requiring sample preparation, chromatographic separation, trained personnel, and/or batching; analytical runtime may be minutes, but total turnaround can be hours to >1 day;

•• - automated or plate-based laboratory assay; moderate hands-on time, usually same-day after sample arrival;

• - rapid measurement, typically ≤10–30 min after setup, minimal washing/separation;

○ – real-time, in vivo, implantable, or continuous monitoring format;

**Cost:**

■■■ – high capital and operational cost, specialized instrumentation and trained staff;

■■ – moderate cost, established laboratory platform or commercial kit;

■ – low per-measurement consumable cost, although specialized sensor setup may be required.;

### Supporting Figures

#### Figure S1. Capture-SELEX workflow for selecting MTX-binding DNA aptamers.

a)

| | Dock / Competitive stem region |
|---|---|
| L45 | 5′-GGA<u>GGCTCTCGGGACGAC</u>-$N_{45}$-GTCGTCCCGCCTTTAGGATTTACAG |
| CO | 3′-Bio-ATA<u>CCGAGAGCCCTGCTG</u> |

b)

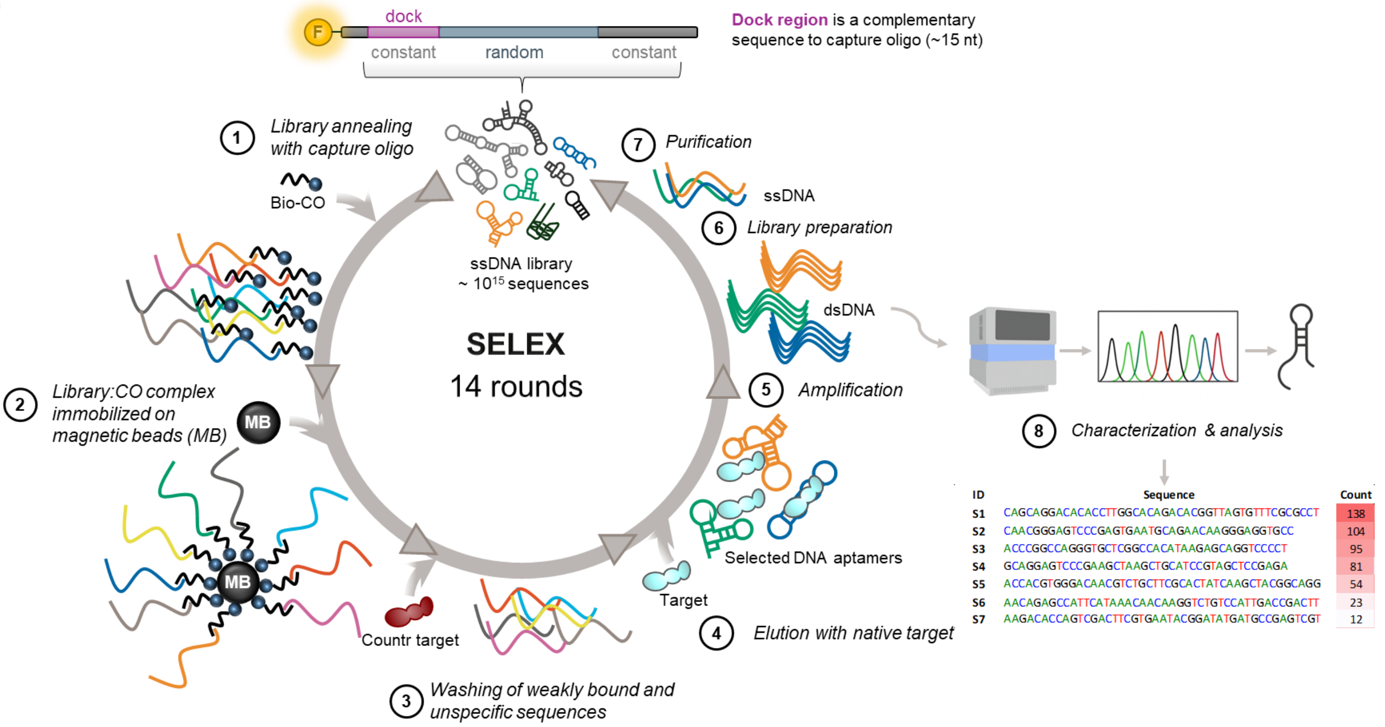

**Figure S1. Capture-SELEX workflow for selecting MTX-binding DNA aptamers**. (a) Design of the ssDNA library (L45) used for Capture-SELEX, including 5′ and 3′ primer-binding regions, a central 45 nt random region ($N_{45}$), and a 15 nt docking sequence.[1–4] The ssDNA library was immobilized on streptavidin-coated magnetic beads (MB) by hybridizing the docking sequence to a complementary biotinylated capture oligonucleotide (CO). The library and capture oligonucleotide were mixed at a 1:2 molar ratio (L45:CO). (b) Schematic representation of the Capture-SELEX strategy employed for enrichment of methotrexate (MTX)-binding aptamers over 14 selection rounds: (1) annealing of library to capture oligonucleotide; (2) immobilization of the L45:CO complex on magnetic beads; (3) removal of nonhybridized sequences and, in later rounds, incubation with the counter-target folic acid (FA); (4) incubation with MTX, resulting in target-induced release of MTX-binding sequences; (5) recovery of eluted sequences followed by PCR amplification; (6) strand separation and (7) regeneration of the ssDNA library for the next selection round; (8) bioinformatic analysis of enriched aptamer candidates, followed by validation and characterization of the top candidates.

**Figure S2. Representative 10% urea-PAGE analysis of the Capture-SELEX workflow.**

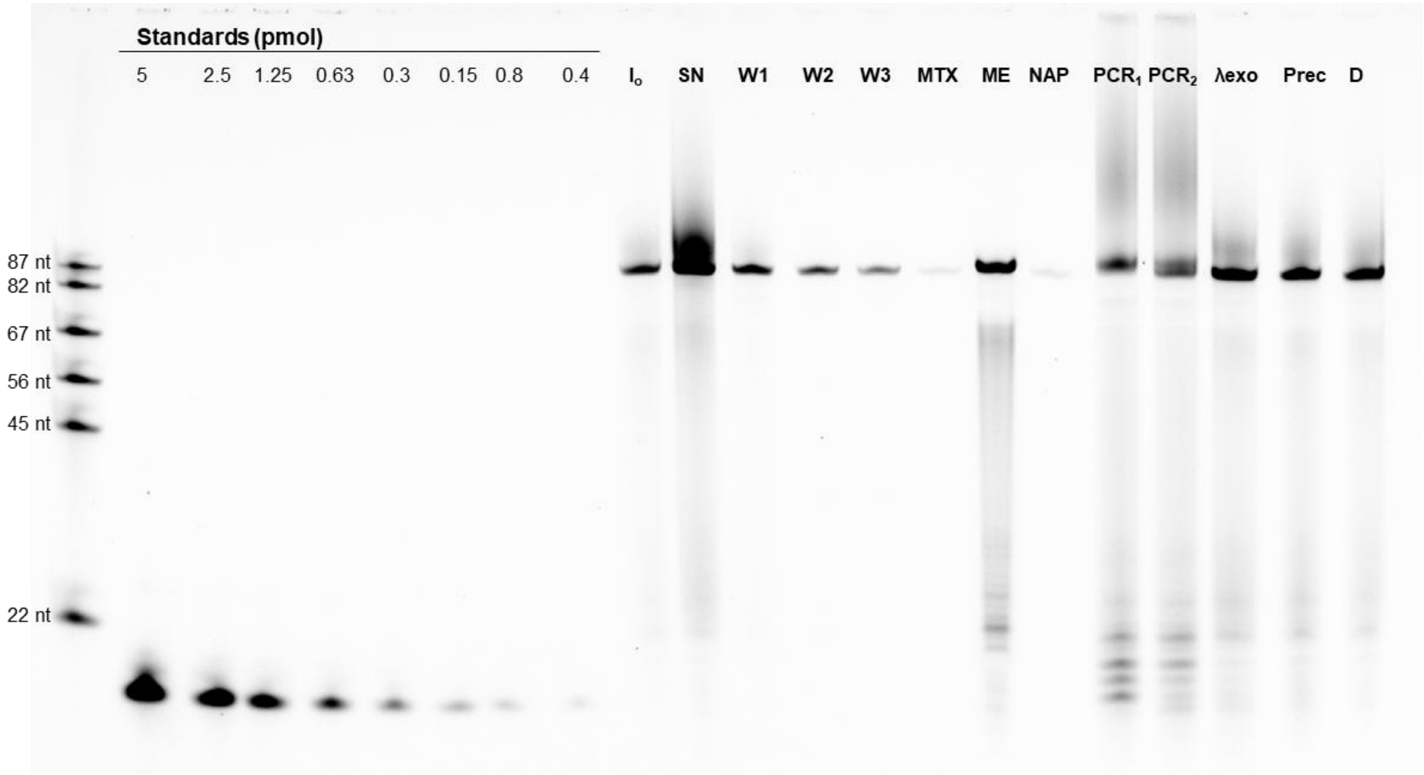

**Figure S2. Representative 10% urea-PAGE analysis of the Capture-SELEX workflow following selection round 4.** Fluorescently labeled DNA samples collected during the selection and recovery workflow were analyzed by 10% denaturing urea-PAGE. $I_0$, initial library:capture oligonucleotide complex before selection round (1:800 dilution); SN, supernatant collected after incubation of the library:capture oligonucleotide complex with magnetic beads (1:80 dilution); W1–W3, wash fractions collected after washing of magnetic beads (1:80 dilution); MTX, library fraction specifically eluted with methotrexate target (1:200 dilution); ME, nonspecific library fraction eluted from magnetic beads using formamide-containing elution buffer (1:800 dilution); NAP, sample after desalting using an illustra NAP-25 column (1:200 dilution); $PCR_1$ and $PCR_2$, products after consecutive PCR amplifications (1:100 and 1:500 dilutions, respectively); λexo, sample after lambda exonuclease digestion of dsDNA (1:575 dilution); Prec, sample after ethanol precipitation (1:800 dilution); D, final desalted and concentrated sample (1:800 dilution). Sample dilutions are indicated to enable comparison between fractions. Fluorescence standards correspond to known amounts (0.04–5 pmol) of FAM-labeled forward primer (F-Pr1) used for approximate quantification of recovered DNA.

**Figure S3. Comparison of sequences and predicted secondary structures of selected and previously reported MTX aptamers.**

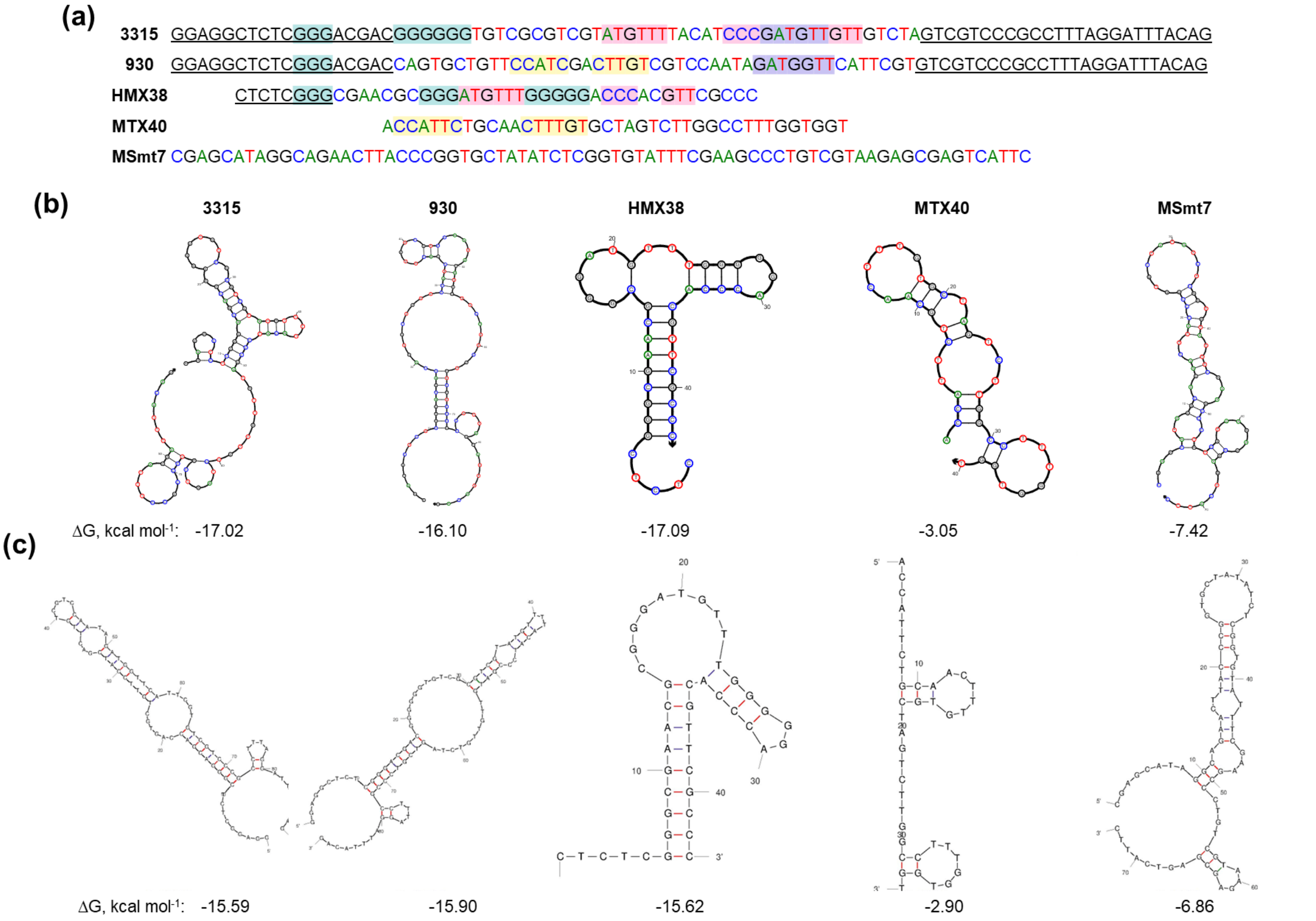

**Figure S3. Comparison of sequences and predicted secondary structures of selected and previously reported MTX aptamers.** (a) Sequence alignment of the MTX-binding aptamers identified in this study, 3315 and 930, with previously reported MTX aptamers HMX38, MTX40, and MSmt7. Overall sequence conservation was limited, indicating that MTX can be recognized by structurally diverse DNA scaffolds. Partial similarity was observed between G-rich regions of 3315 and HMX38, suggesting that G-rich structural elements may contribute to MTX recognition in distinct aptamer architectures. Conserved or partially similar sequence regions are highlighted. Predicted secondary structures were generated using (b) NUPACK and (c) Mfold under DNA folding conditions at 22 °C with 100 mM $Na^+$ and 2 mM $Mg^{2+}$. Predicted Gibbs free energy values (ΔG, kcal $mol^{-1}$) are indicated below the corresponding structures.

**Figure S4. Evaluation of G-quadruplex formation by MTX-specific aptamers using SG4-based ELISA.**

**a)**

| | |
|---|---|
| **MycG4** | TGAGGGTGGGTAGGGTGGGTAA |
| **mMycG4** | TGAGTGTGCGTAGTGTGTGTAA |
| **3315-88** | GGAGGCTCTCGGGACGACGGGGGGTGTCGCGTCGTATGTTTTACATCCCGATGTTGTTGTCTAGTCGTCCCGCCTTTAGGATTTACAG |
| **930-88** | GGAGGCTCTCGGGACGACCAGTGCTGTTCCATCGACTTGTCGTCCAATAGATGGTTCATTCGTGTCGTCCCGCCTTTAGGATTTACAG |
| **17-43** | ACTCGTCTCTTGGGGTTTGGCGTCGGGCCTGGCGTGGACGAGT |

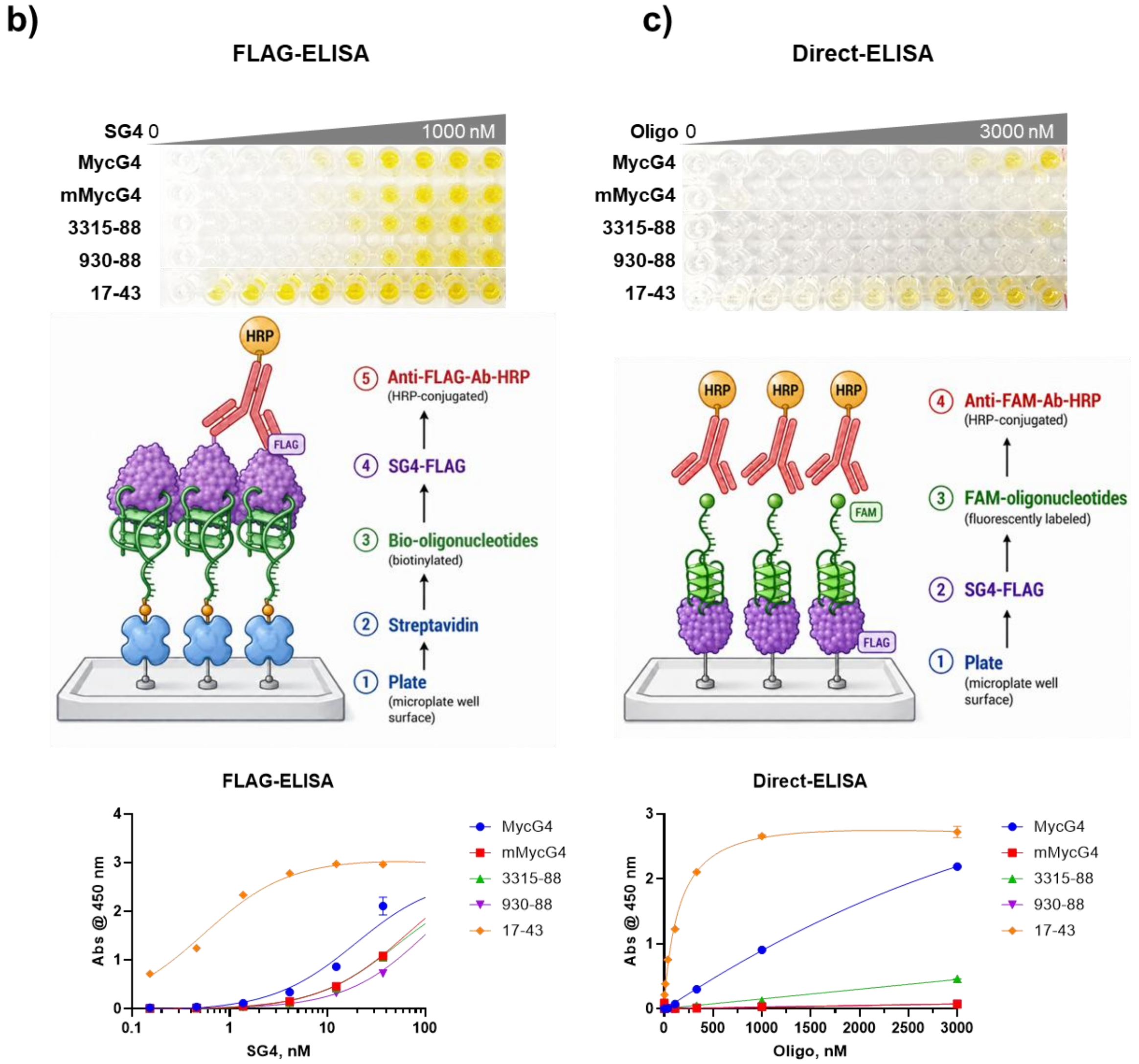

**Figure S4. Evaluation of G-quadruplex formation by MTX-specific aptamers using SG4-based ELISA.** (a) Sequence comparison of MTX-specific aptamers and control oligonucleotides used for SG4-binding analysis. Known G-quadruplex-forming sequences, including the classical MycG4 motif [7] and the previously reported S100A7 aptamer 17-43,[8] were included as positive controls, while the non-G4 mutant mMycG4 was used as a negative control. Putative G-rich tracts capable of contributing to G-quadruplex formation are highlighted in yellow, and mutated residues in the non-G4 control are shown in red. (b) FLAG-ELISA format used to evaluate SG4 binding to immobilized biotinylated oligonucleotides. In this assay, streptavidin-coated plates were incubated with biotinylated oligonucleotides, followed by SG4-FLAG and detection with an anti-FLAG-HRP antibody. (c) Direct-ELISA format used to evaluate binding of FAM-labelled oligonucleotides to immobilized SG4-FLAG. In this assay, plates were coated with SG4-FLAG, incubated with FAM-labelled oligonucleotides, and detected with anti-FAM-HRP antibody. Dose–response curves show SG4-dependent recognition of G-quadruplex-forming positive controls, whereas MTX aptamers 3315 and 930 produced signals comparable to the non-G4 control under the tested conditions. Data represent mean ± SD from 3–5 replicate measurements.

**Figure S5. Biolayer interferometry (BLI) kinetic analysis of selected MTX-specific aptamers**

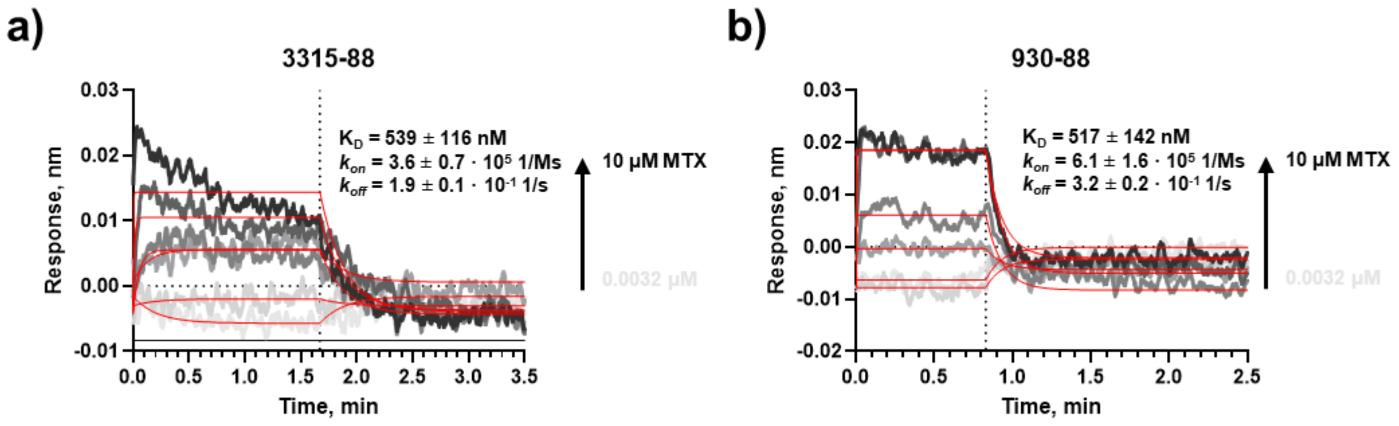

**Figure S5. Biolayer interferometry (BLI) kinetic analysis of selected MTX-specific aptamers.** Representative BLI sensorgrams showing association and dissociation kinetics of MTX binding to immobilized aptamers measured on the Octet RED96 platform. (a) BLI kinetic profile of aptamer 3315-88. (b) BLI kinetic profile of aptamer 930-88. Curves were fitted using a global 1:1 binding model to determine kinetic parameters and equilibrium dissociation constants ($K_D$).

**Figure S6. Fluorescent strand-displacement assay for MTX detection using selected aptamers.**

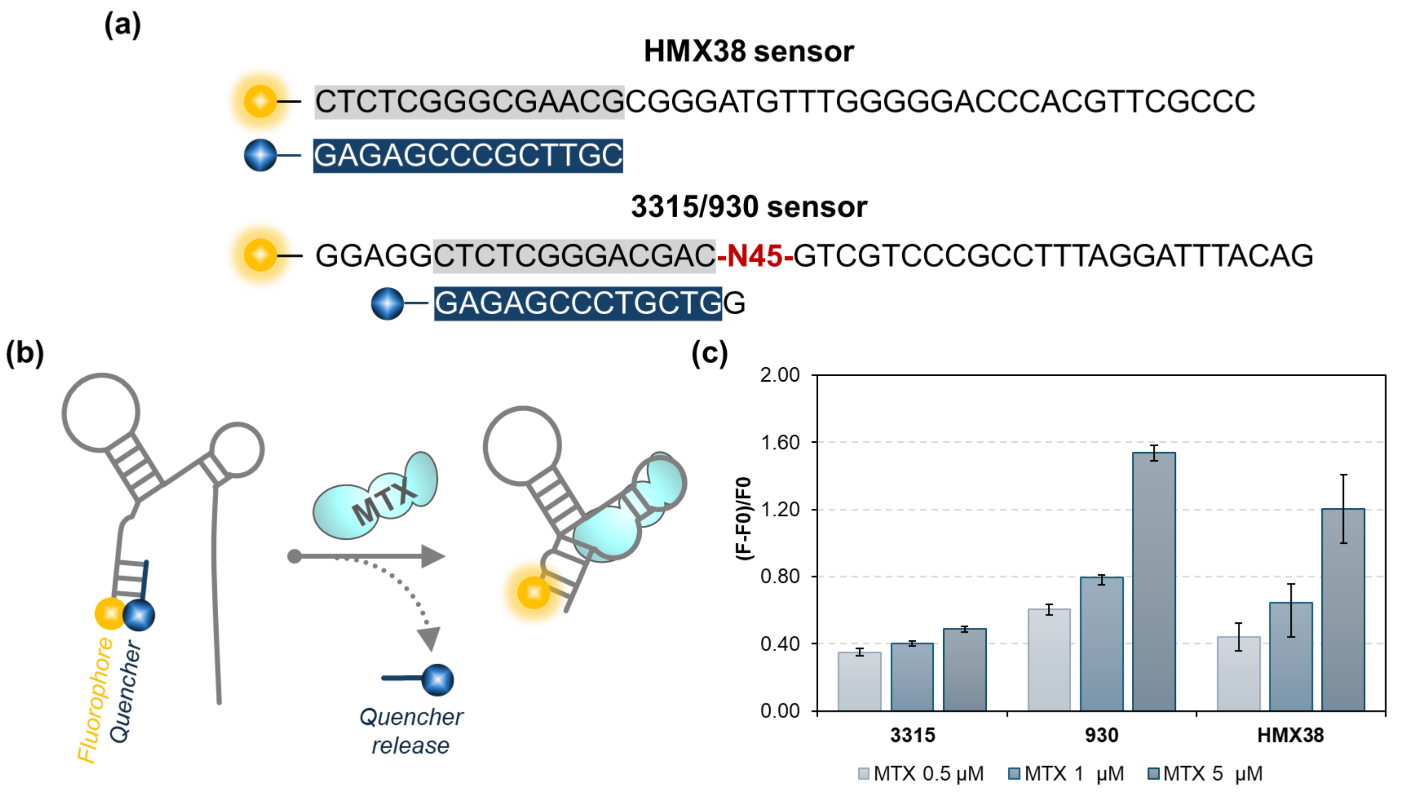

**Figure S6. Fluorescent strand-displacement sensing of MTX using selected aptamers.** (a) Sequence design of previously reported HMX38-based MTX fluorescent sensor together with selected 3315 and 930 aptamer-based sensor constructs, including fluorophore- and quencher-labeled complementary strands. (b) Schematic representation of the strand-displacement sensing mechanism employing a FAM-labelled aptamer hybridized to an Iowa Black quencher strand. MTX binding induces a conformational rearrangement of the aptamer, displacing the quencher strand and restoring the fluorescence signal. (c) Fluorescence response of strand-displacement sensors in the presence of increasing MTX concentrations (0, 0.5, 1, and 5 μM). Assays were performed using 0.1 μM FAM-labelled aptamer strand and 0.2 μM quencher strand in selection buffer (SB) at room temperature. Data represent the mean ± SD from five independent measurements.

**Figure S7. Temperature-dependent ITC analysis of MTX-specific aptamers.**

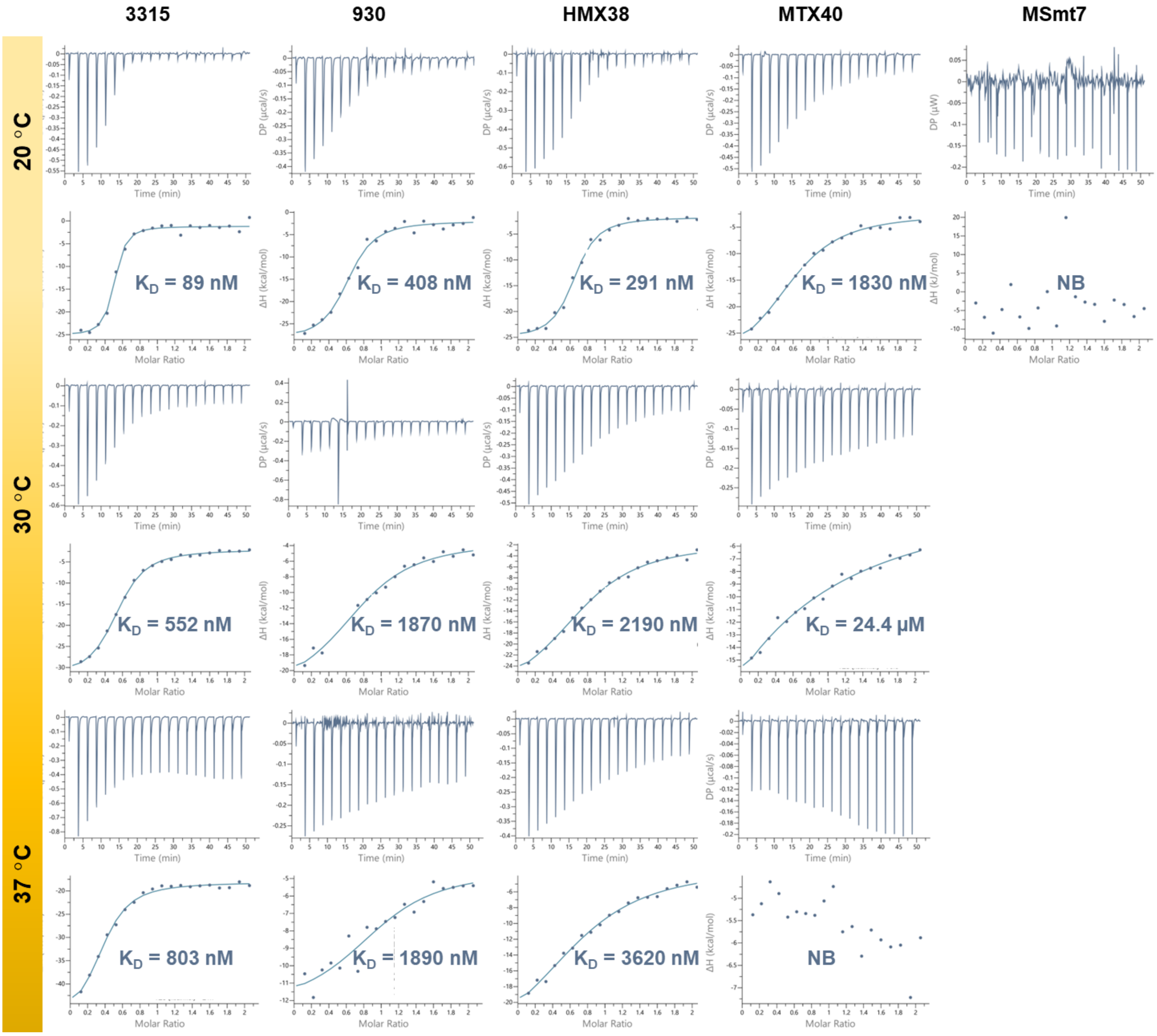

**Figure S7.** Temperature-dependent ITC analysis of MTX-specific aptamers. Representative ITC isotherms of selected aptamers 3315 and 930 together with previously reported MTX aptamers HMX38, MTX40, and MSmt7 measured in selection buffer (SB) at 20, 30, and 37 °C. Experiments were performed using 100 μM MTX in the syringe and 10 μM aptamer in the cell.

**Figure S8. Effect of buffer composition on MTX aptamer binding determined by ITC.**

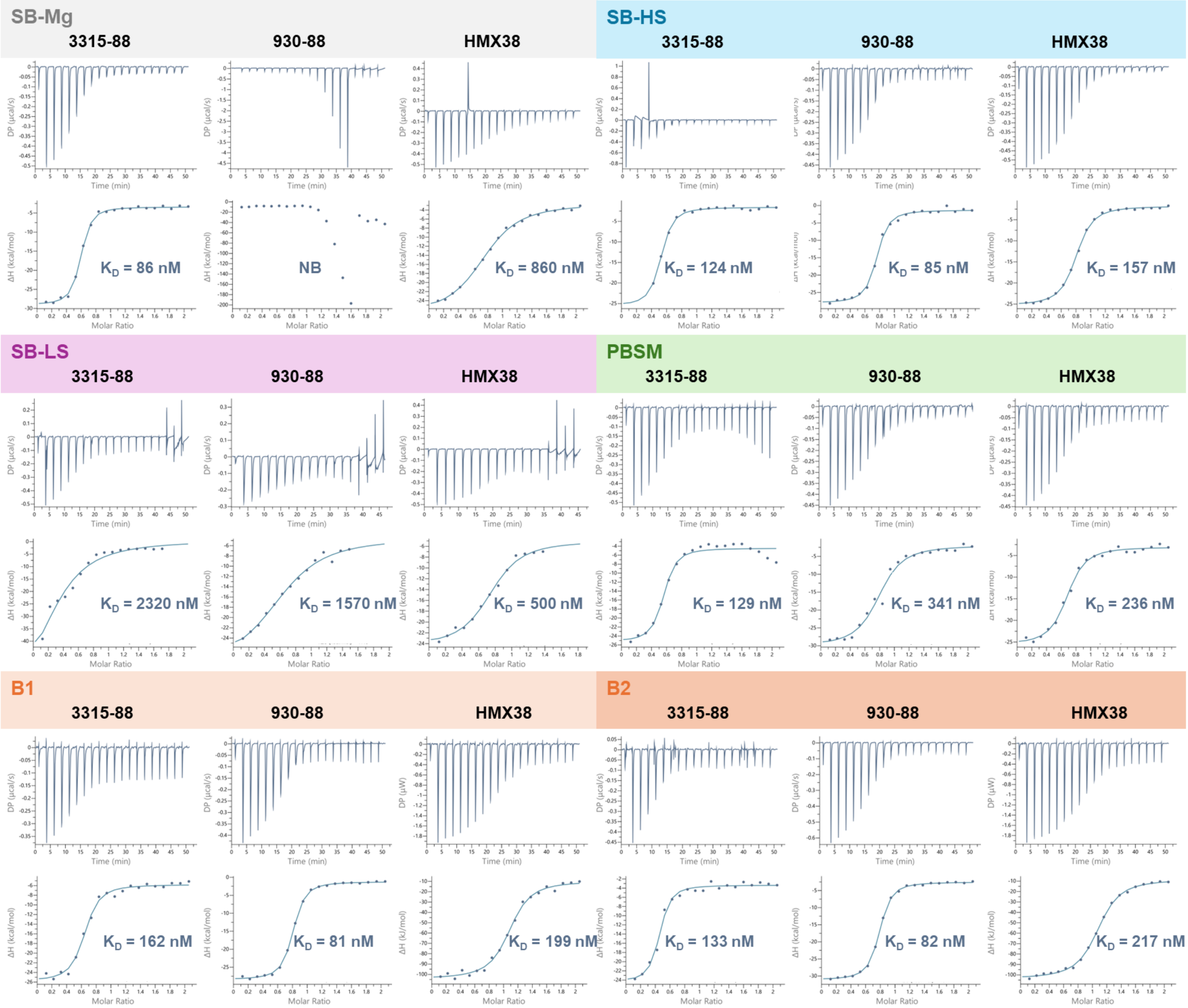

**Figure S8. Effect of buffer composition on MTX aptamer binding determined by isothermal titration calorimetry (ITC).** Representative ITC thermograms (top panels) and integrated binding isotherms (bottom panels) for aptamers 3315-88, 930-88, and the previously reported MTX aptamer HMX38 interacting with MTX under different buffer conditions. Measurements were performed in SB-Mg (40 mM Tris-HCl, 100 mM NaCl, 1 mM $CaCl_2$, pH 7.4), SB-HS (20 mM Tris-HCl, 250 mM NaCl, 5 mM $MgCl_2$, 1 mM $CaCl_2$, pH 7.4), SB-LS (20 mM Tris-HCl, 20 mM NaCl, 0.5 mM $MgCl_2$, 1 mM $CaCl_2$, pH 7.4), PBSM (10 mM phosphate buffer, 137 mM NaCl, 2.7 mM KCl, 5 mM $MgCl_2$, pH 7.4), B1 (50 mM Tris-HCl, 100 mM NaCl, 10 mM $MgCl_2$, pH 7.9), and B2 (50 mM Tris-HCl, 100 mM NaCl, 10 mM $MgCl_2$, 100 µg $mL^{-1}$ recombinant albumin, pH 7.9). Apparent dissociation constants ($K_D$) obtained by fitting the data to a one-site binding model are indicated within each panel. No measurable binding (NB) was observed for 930-88 in SB-Mg. Experimental conditions were identical to those described in Figure 2, with 100 µM MTX in the syringe and 10 µM aptamer in the cell at 20 °C.

**Figure S9. ITC analysis of MTX binding by selected aptamers in 10% human serum.**

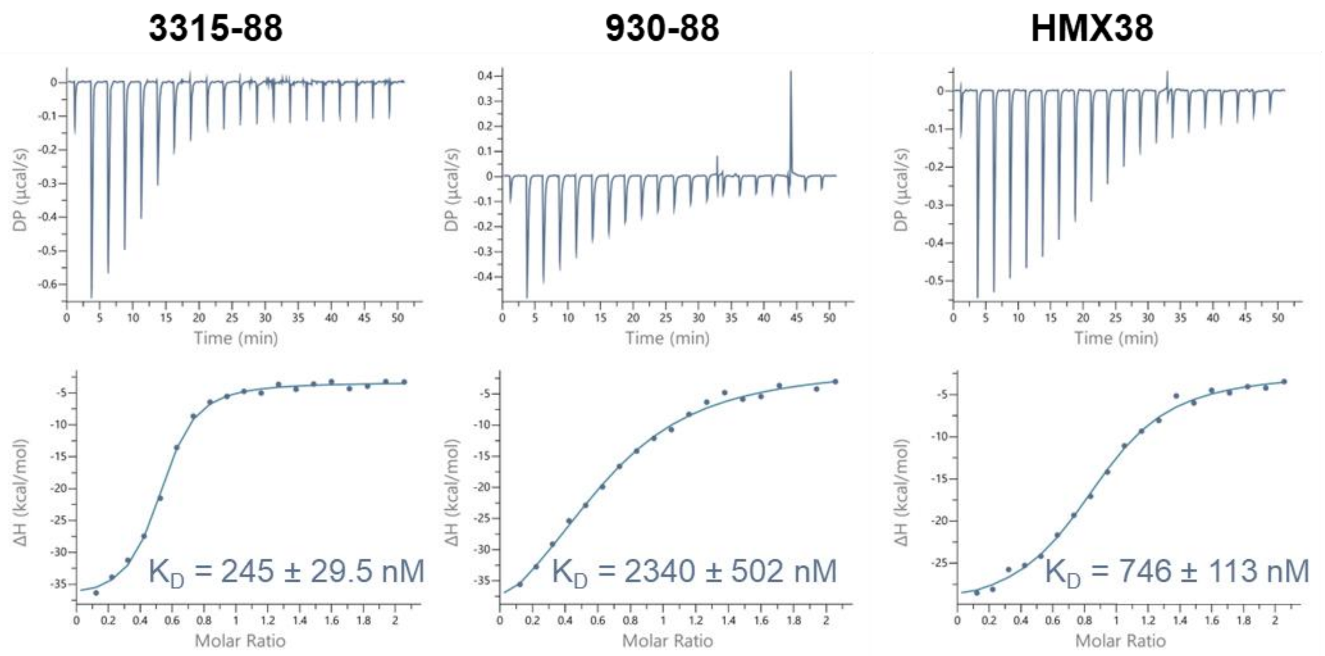

**Figure S9. ITC analysis of MTX binding by selected aptamers in 10% human serum.** Representative ITC thermograms and corresponding integrated binding isotherms for selected MTX-binding aptamers measured in selection buffer supplemented with 10% human serum. Experiments were performed at 20 °C using 100 µM MTX in the syringe and 10 µM aptamer in the cell. Binding isotherms were fitted using a single-site binding model after correction for heats of dilution. Aptamer 3315-88 retained nanomolar MTX binding in serum-containing buffer, whereas 930-88 showed substantially weaker binding under the same conditions. The previously reported HMX38 aptamer was included for comparison. Fitted apparent $K_D$ values are shown on each plot.

**Figure S10. Specificity analysis of selected MTX-binding aptamers by ITC.**

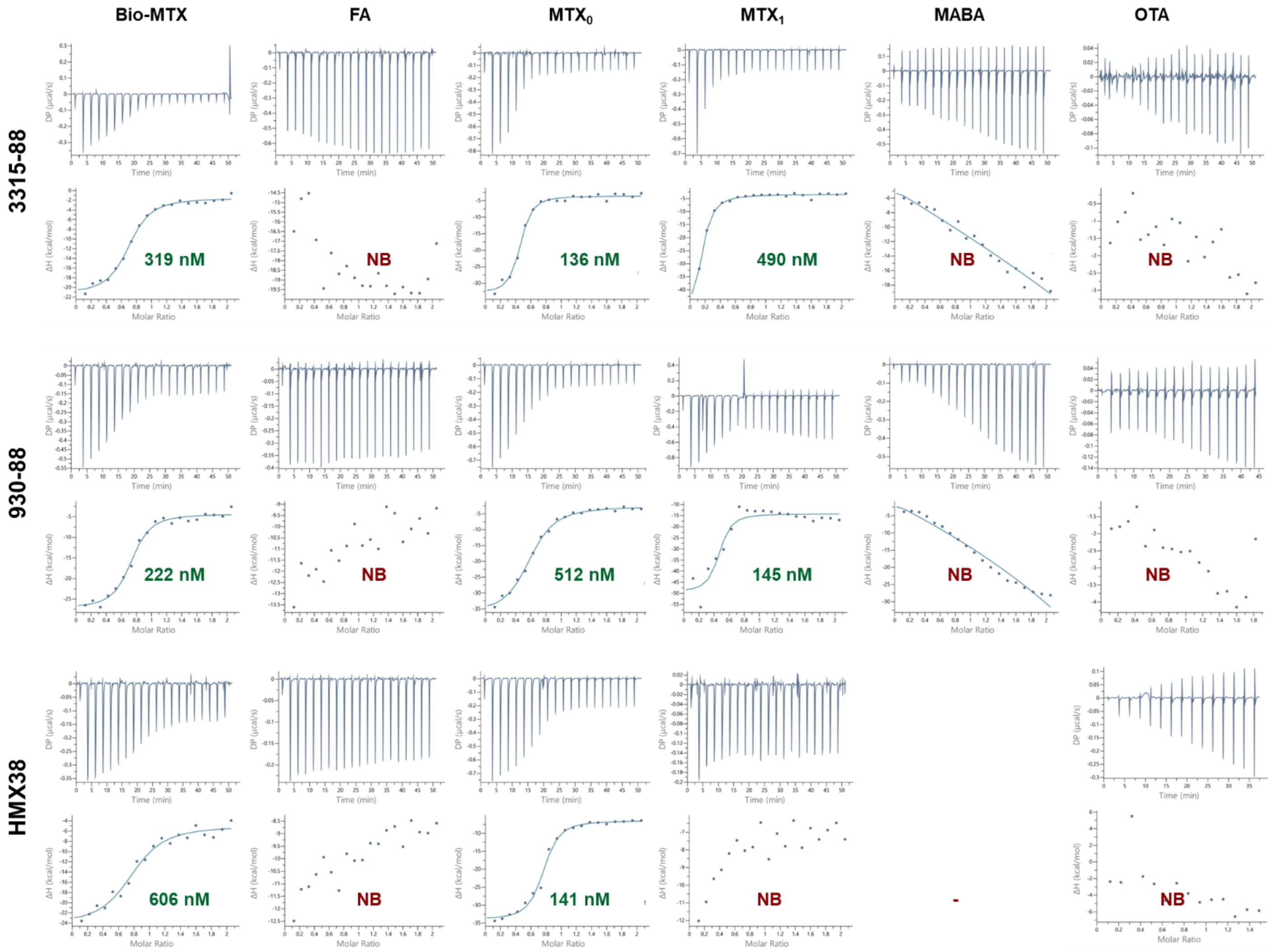

**Figure S10. Specificity analysis of selected MTX-binding aptamers by ITC.** Representative ITC raw thermograms and integrated binding isotherms for aptamers 3315-88, 930-88, and the previously reported HMX38 aptamer measured against MTX and structurally related or unrelated compounds. Tested analytes included biotinylated MTX (Bio-MTX), folic acid (FA), 4-deoxy-4-amino-N10-methylpteroic acid ($MTX_0$/DAMPA), 2,4-diamino-6-(hydroxymethyl)pteridine ($MTX_1$), 4-methylaminobenzoic acid (MABA), and ochratoxin A (OTA). Experiments were performed at 20 °C using 100 µM analyte in the syringe and 10 µM aptamer in the cell under matched buffer and solvent conditions. Apparent $K_D$ values obtained from single-site fitting are indicated below each isotherm; NB indicates no detectable binding under the tested conditions. Both 3315-88 and 930-88 showed binding to MTX and $MTX_0$/DAMPA, but no detectable binding to FA, MABA, or OTA, supporting selective recognition of the MTX-related scaffold rather than nonspecific binding to unrelated aromatic or folate-like molecules.

### Figure S11. Truncation design and MTX-binding performance of 3315 aptamer variants.

| | Length. nt | Sequence, 5' → 3' | $K_D$, nM |
|---|---|---|---|
| 3315 | 88 | GGAGGCTCTCGGGACGACGGGGGGTGTCGCGTCGTATGTTTTACATCCCGATGTTGTTGTCTAGTCGTCCCGCCTTTAGGATTTACAG | 89 |
| | 63 | CGGGACGACGGGGGGTGTCGCGTCGTATGTTTTACATCCCGATGTTGTTGTCTAGTCGTCCCG | 37 |
| | 61 | GGGACGACGGGGGGTGTCGCGTCGTATGTTTTACATCCCGATGTTGTTGTCTAGTCGTCCC | 146 |
| | 60 | ATACGACGGGGGGTGTCGCGTCGTATGTTTTACATCCCGATGTTGTTGTCTAGTCGTCCC | 231 |
| | 60a | AGGACGACGGGGGGTGTCGCGTCGTATGTTTTACATCCCGATGTTGTTGTCTAGTCGTCC | 197 |
| | 57 | GCCGGCGGGGGGTGTCGCGTCGTATGTTTTACATCCCGATGTTGTTGTCTAGCCGGC | 226 |
| | 55 | CCGGCGGGGGGTGTCGCGTCGTATGTTTTACATCCCGATGTTGTTGTCTAGCCGG | 63 |
| | 55a | ACGACGGGGGGTGTCGCGTCGTATGTTTTACATCCCGATGTTGTTGTCTAGTCGT | 146 |
| | 53 | CGGCGGGGGGTGTCGCGTCGTATGTTTTACATCCCGATGTTGTTGTCTAGCCG | 80 |
| | 52 | GGAGGCTCTCGGGACGACGGGGGGTGTCGCGTCGTATGTTTTACATCCCGAT | NB |
| | 50 | AATACGACGGGGGGTGTCGCGTCGTATGTTTTACATCCCGATGTTGTTGT | NB |
| | 47 | ACGACGGGGGGTGTCGCGTCGTATGTTTTACATCCCGATGTTGTCGT | NB |
| | 45 | GGGGGGTGTCGCGTCGTATGTTTTACATCCCGATGTTGTTGTCTA | NB |
| | 45a | CCGGCGGGGGGTGTCGCGTCGTATGTTTTACATCCCGATAGCCGG | NB |
| | 39 | CCGGCGGGGGGTGTCGCGTCGTATGTTTTACATCCCGAT | NB |
| | 39a | ACGACGGGGGGTGTCGCGTCGTATGTTTTACATCCCGAT | NB |
| | 38 | CGGCGGGGGGTGTCGCGTCGTATGTTTTACATCCCGAT | NB |

**Figure S11. Truncation design and MTX-binding performance of 3315 aptamer variants.** Sequence alignment of the full-length 3315 aptamer and its truncated variants, with apparent $K_D$ values determined by ITC shown on the right. The original randomized region is highlighted in yellow, primer-derived regions are shown in grey and underlined, and mutations introduced relative to the original 3315 sequence are indicated in red. Apparent $K_D$ values are color-coded according to binding affinity; NB indicates no detectable binding under the tested conditions.

**Figure S12. Validation of MTX binding by truncated 3315 aptamer variants using ITC.**

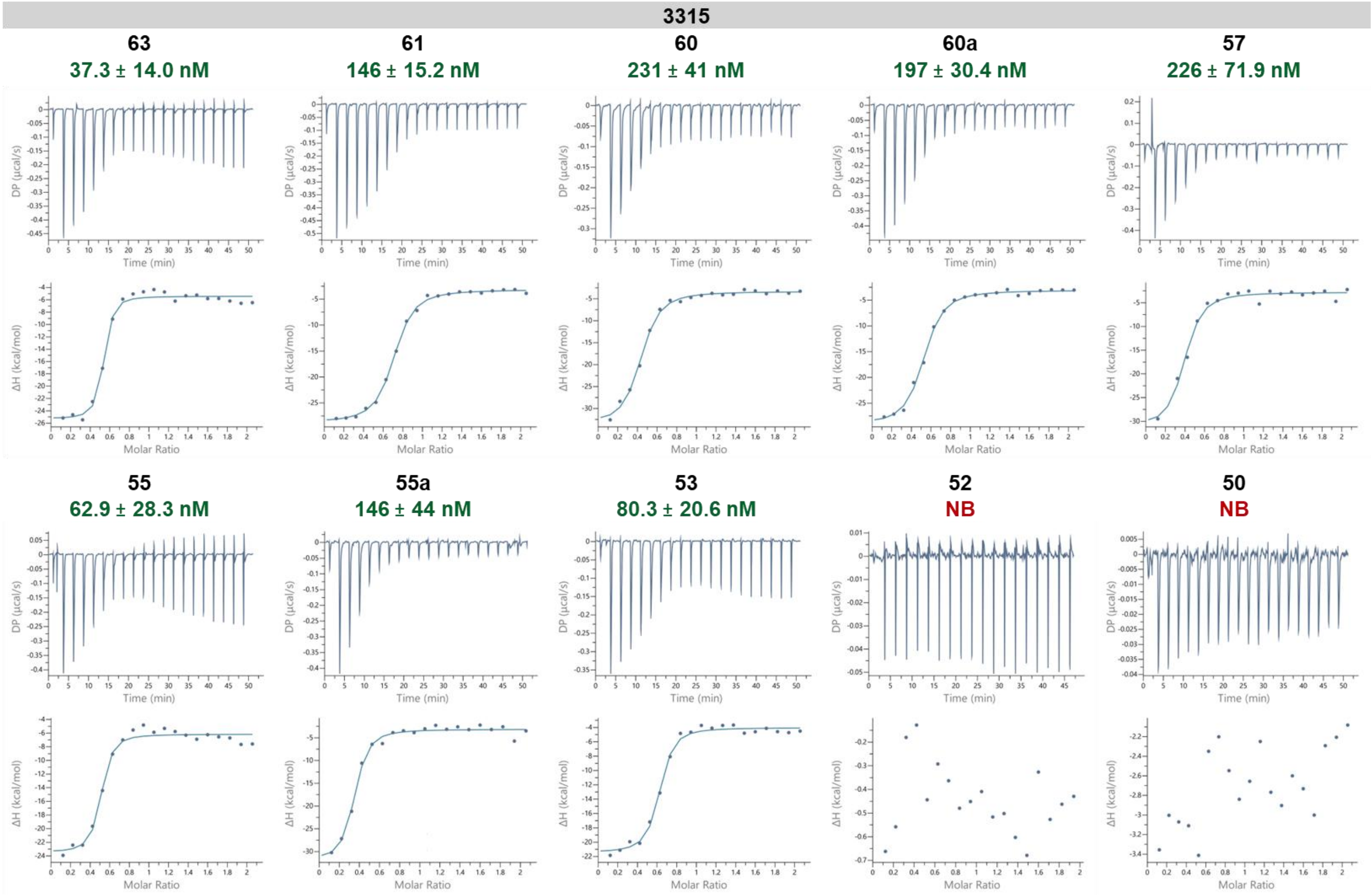

**Figure S12. Validation of MTX binding by truncated 3315 aptamer variants using ITC.** Representative ITC raw thermograms and integrated binding isotherms for selected truncated variants of the 3315 aptamer. Experiments were performed under standard ITC conditions at 20 °C, with 100 µM MTX titrated into a 10 µM aptamer solution. $K_D$ values obtained from fitting the binding isotherms are indicated above each dataset and reported as $K_D$ ± fitting error. NB indicates no detectable binding under the tested conditions. Isotherms for truncation variants shorter than 50 nt are not shown because these variants showed no detectable MTX binding under the tested conditions.

**Figure S13. Truncation design and predicted secondary structures of the 930 MTX-specific aptamer.**

| | Length, nt | Sequence, 5' → 3' | $K_D$, nM | ΔG, kcal/mol Mfold | |
|---|---|---|---|---|---|
| | | | | 1 | 2 |
| 930 | 88 | GGAGGCTCTCGGGACGACCAGTGCTGTTCCATCGACTTGTCGTCCAATAGATGGTTCATTCGTGTCGTCCCGCCTTTAGGATTTACAG | 408 | -16.97 | -16.57 |
| | 71 | CGGGACGACCAGTGCTGTTCCATCGACTTGTCGTCCAATAGATGGTTCATTCGTGTCGTCCCGCCTTTAGG | 590 | -15.53 | -15.13 |
| | 67 | GGCGGGACGACCAGTGCTGTTCCATCGACTTGTCGTCCAATAGATGGTTCATTCGTGTCGTCCCGCC | 862 | -19.91 | -19.51 |
| | 65 | CCGGGACGACCAGTGCTGTTCCATCGACTTGTCGTCCAATAGATGGTTCATTCGTGTCGTCCCGG | 786 | -17.41 | -17.01 |
| | 63 | CGGGACGACCAGTGCTGTTCCATCGACTTGTCGTCCAATAGATGGTTCATTCGTGTCGTCCCG | 578 | -15.38 | -14.98 |
| | 53 | CGGCCAGTGCTGTTCCATCGACTTGTCGTCCAATAGATGGTTCATTCGTGCCG | 839 | -7.76 | -6.97 |
| | 45 | CAGTGCTGTTCCATCGACTTGTCGTCCAATAGATGGTTCATTCGT | NB | -6.20 | -5.24 |

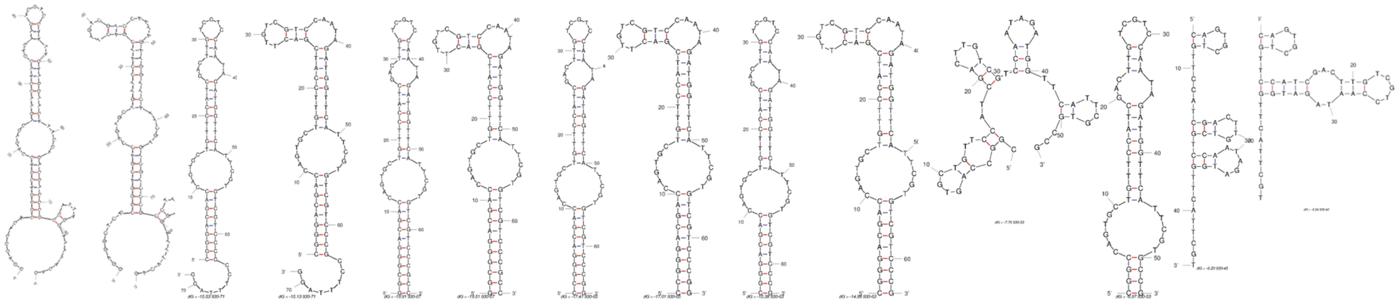

| 88 | 71 | 67 | 65 | 63 | 53 | 45 |
|---|---|---|---|---|---|---|
| ΔG = -16.97 kcal/mol | ΔG = -15.53 kcal/mol | ΔG = -19.91 kcal/mol | ΔG = -17.41 kcal/mol | ΔG = -15.38 kcal/mol | ΔG = -7.76 kcal/mol | ΔG = -6.20 kcal/mol |
| ΔG = -16.57 kcal/mol | ΔG = -15.13 kcal/mol | ΔG = -19.51 kcal/mol | ΔG = -17.01 kcal/mol | ΔG = -14.98 kcal/mol | ΔG = -6.97 kcal/mol | ΔG = -5.24 kcal/mol |

**Figure S13. Truncation design and predicted secondary structures of the 930 MTX-specific aptamer.** a) Sequence alignment of the full-length 930 aptamer and its truncated variants. The randomized region is highlighted in yellow, mutations introduced relative to the original 930 sequence are indicated in red, self-complementary regions are shaded in grey, and primer-binding regions are underlined. b) Mfold-predicted secondary structures of selected 930 truncation variants. All structures were predicted under standard conditions of 22 °C, 100 mM $Na^+$, and 2 mM $Mg^{2+}$.

**Figure S14. Validation of MTX binding by truncated 930 aptamer variants using ITC.**

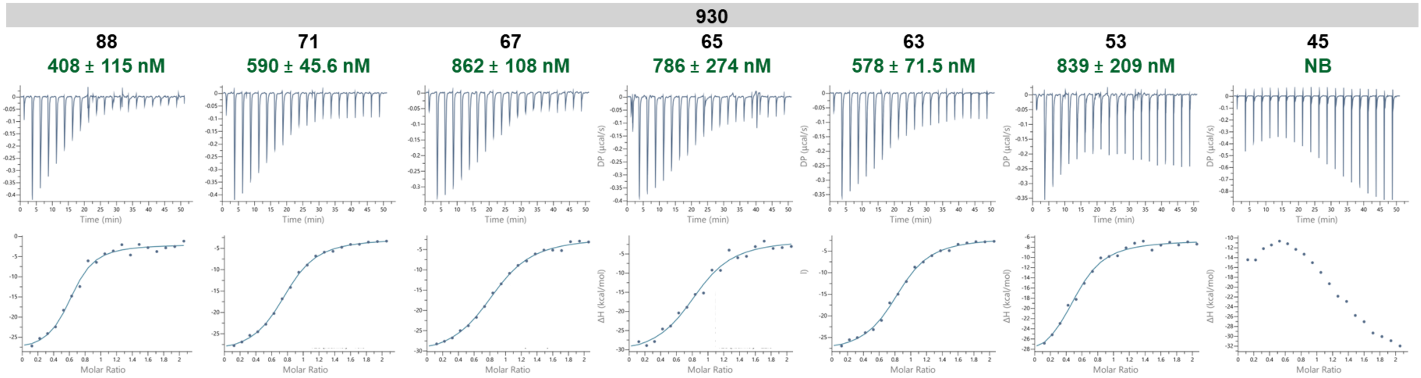

**Figure S14. Validation of MTX binding by truncated 930 aptamer variants using ITC.** Representative ITC raw thermograms and integrated binding isotherms for selected truncated variants of the 930 aptamer. Experiments were performed under standard ITC conditions at 20 °C, with 100 µM MTX titrated into 10 µM aptamer. $K_D$ values obtained from fitting the binding isotherms are indicated above each dataset and reported as $K_D$ ± fitting error. NB indicates no detectable binding under the tested conditions.

**Figure S15. Analysis of 930 aptamer truncation variants in B1 buffer.**

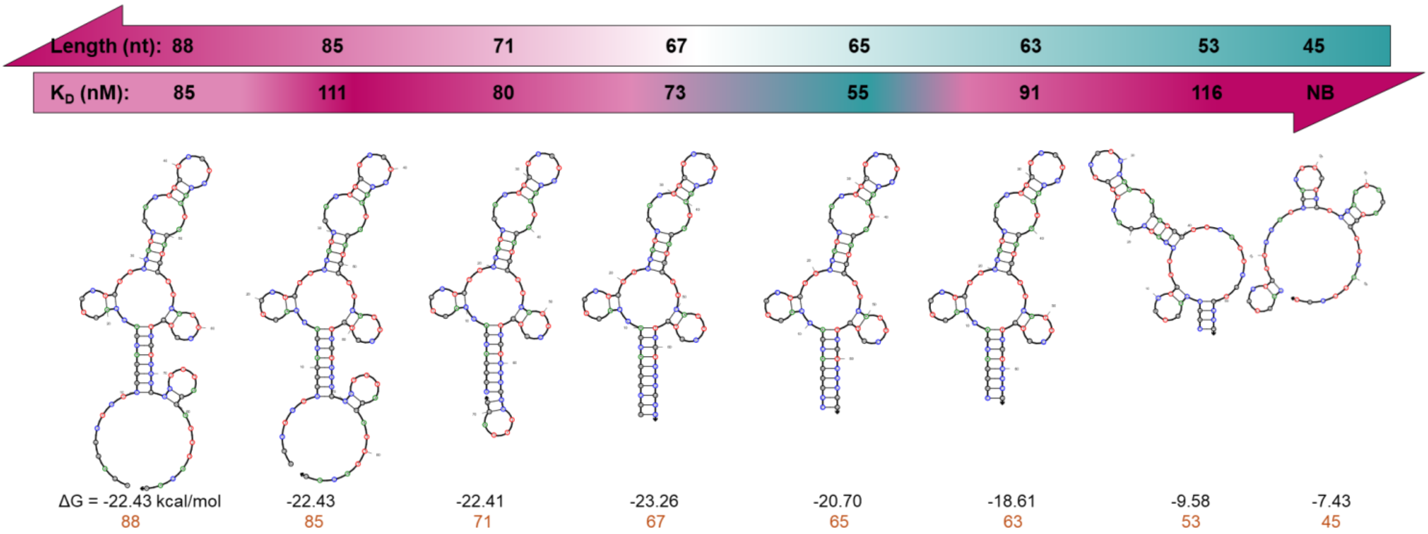

**Figure S15. Analysis of 930 aptamer truncation variants in B1 buffer.** Sequential truncation variants of the 930 aptamer were analyzed by ITC for MTX binding in B1 buffer containing 10 mM $Mg^{2+}$. Predicted secondary structures and corresponding $K_D$ values are shown for each variant, and binding affinities were plotted as a function of aptamer length. The 65-nt variant displayed the strongest MTX binding, whereas further truncation decreased affinity or resulted in no detectable binding.

**Figure S16. Effect of temperature on the binding affinity of truncated 3315 aptamer variants.**

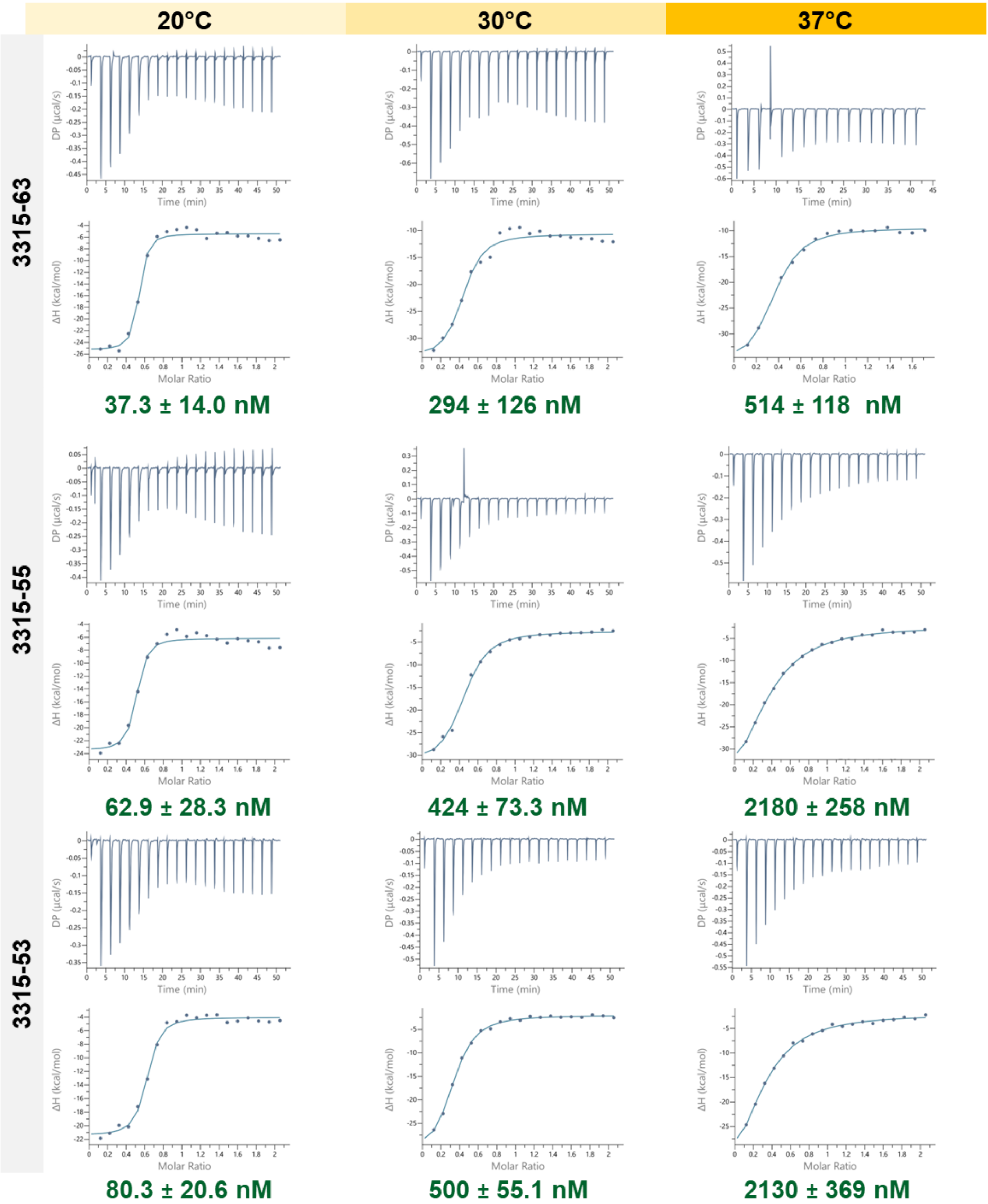

**Figure S16. Effect of temperature on the binding affinity of truncated 3315 aptamer variants.** Representative ITC raw thermograms and corresponding integrated binding isotherms for 3315-63, 3315-55, and 3315-53 measured at 20, 30, and 37 °C. All experiments were performed in SB buffer under standard ITC conditions, with 100 µM MTX titrated into 10 µM aptamer. $K_D$ values derived from fitting the binding isotherms are shown below each dataset and reported as $K_D$ ± fitting error.

**Figure S17. Effect of temperature on the binding affinity of truncated 930 aptamer variants.**

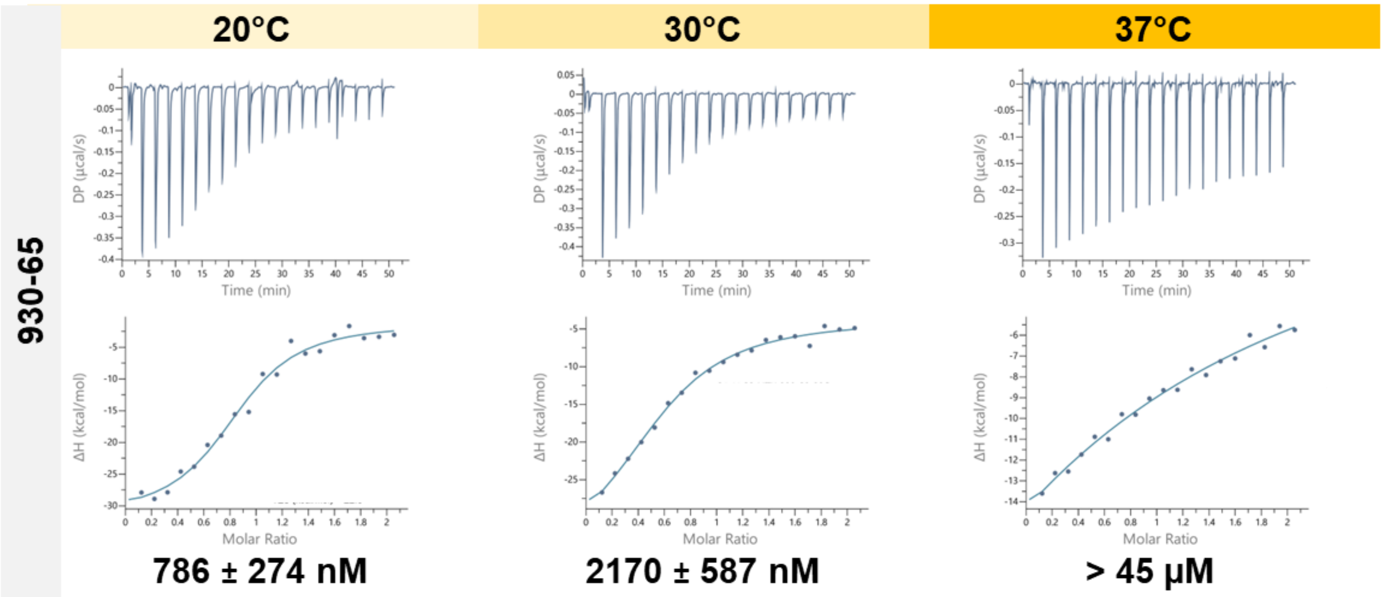

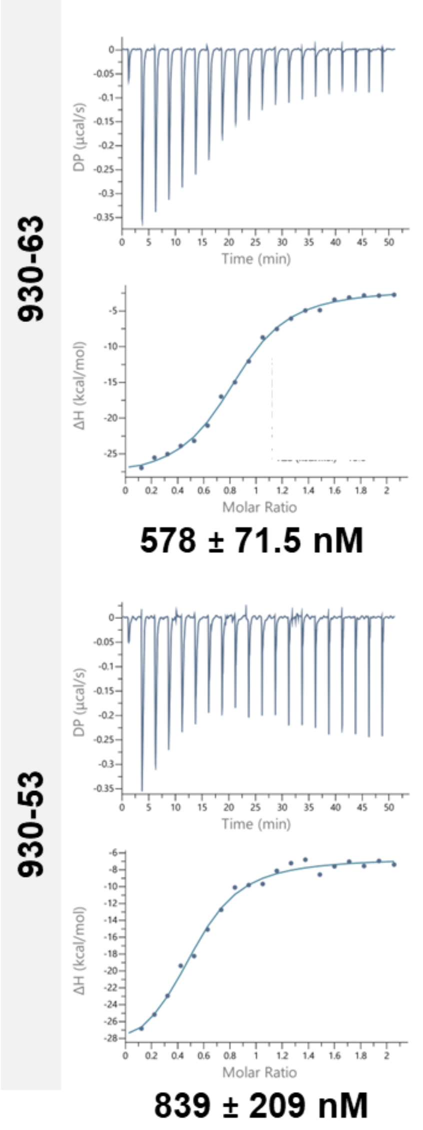

| Aptamer | Length, nt | Temperature 20°C | 30°C | 37°C |
|---|---|---|---|---|
| 3315 | 88 | 89 | 552 | 803 |
| | 63 | 37 | 294 | 514 |
| | 55 | 63 | 424 | 2180 |
| | 53 | 80 | 500 | 2130 |
| 930 | 88 | 408 | 1870 | 1890 |
| | 65 | 786 | 2170 | >45,000 |
| | 63 | 578 | | |
| | 53 | 839 | | |
| HMX38 | 36 | 291 | 2190 | 3620 |
| MTX40 | 40 | 1830 | >24,400 | NB |

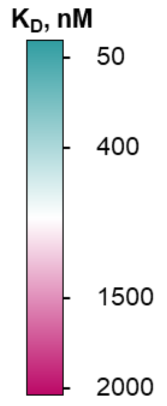

**Figure S17. Effect of temperature on the binding affinity of truncated 930 aptamer variants.** Representative ITC thermograms and integrated binding isotherms for 930-65 at 20, 30, and 37 °C, and for 930-63 and 930-53 at 20 °C. All measurements were performed in SB buffer using 100 µM MTX in the syringe and 10 µM aptamer in the cell. $K_D$ values are shown below each dataset and reported as $K_D$ ± fitting error. The heat map summarizes temperature-dependent binding affinities for all tested aptamer variants.

**Figure S18. Buffer-dependent binding performance of truncated 3315 and 930 aptamer variants**

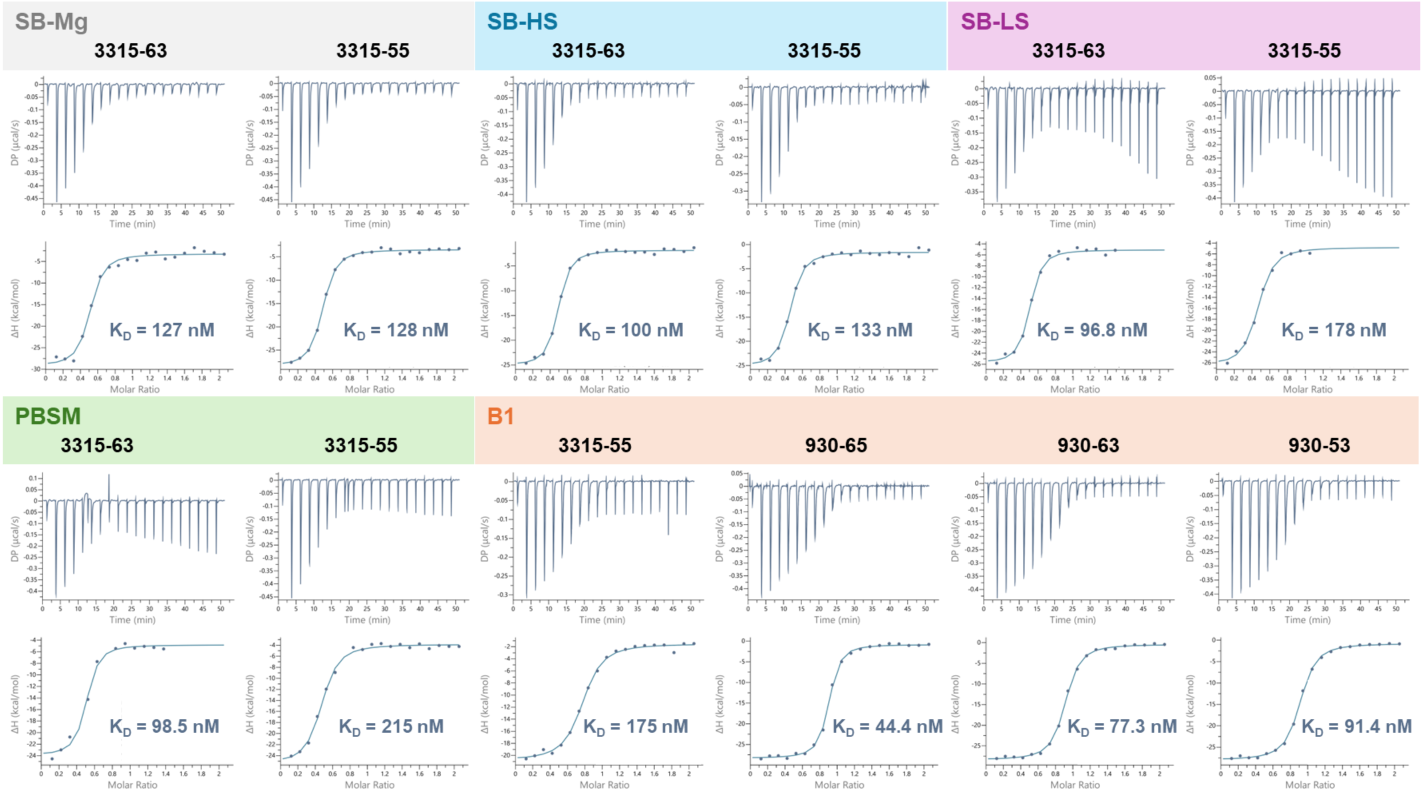

**Figure S18. Buffer-dependent binding performance of truncated 3315 and 930 aptamer variants.** Representative ITC thermograms and integrated binding isotherms showing the effect of buffer composition on MTX binding by selected truncated 3315 and 930 aptamer variants. Measurements were performed at 20 °C with 100 µM MTX titrated into 10 µM aptamer. Binding was evaluated in SB without $Mg^{2+}$, high-salt SB, low-salt SB, PBSM, and B1 buffer. Fitted $K_D$ values are indicated below each dataset and reported as $K_D$ ± fitting error. Buffer compositions are listed in Table S2.

**Figure S19. Binding performance of truncated 3315 aptamer variants in human serum-containing buffer.**

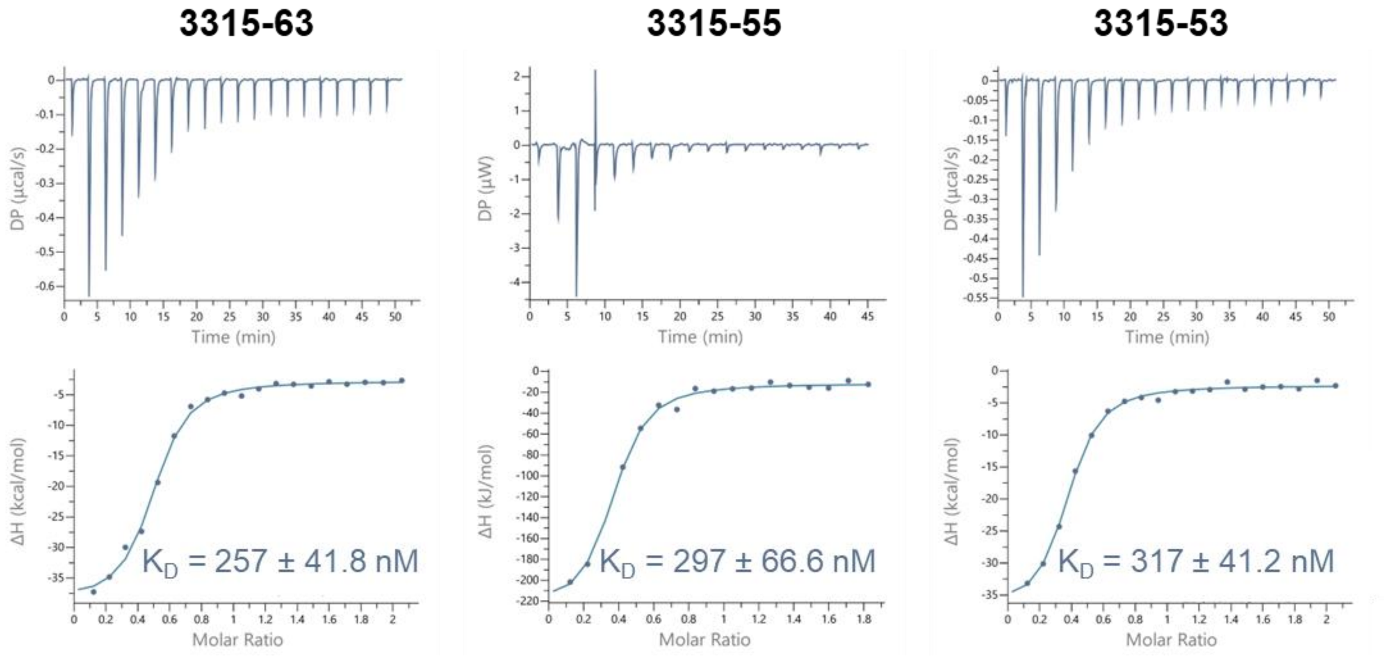

**Figure S19. Binding performance of truncated 3315 aptamer variants in human serum-containing buffer.** Representative ITC thermograms and integrated binding isotherms for selected truncated 3315 aptamer variants measured in the presence of 10% heat-inactivated human serum. Experiments were performed at 20 °C with 100 µM MTX in the syringe and 10 µM aptamer in the cell. $K_D$ values derived from isotherm fitting are shown below each dataset and reported as $K_D$ ± fitting error.

**Figure S20. Binding specificity of truncated MTX aptamer variants.**

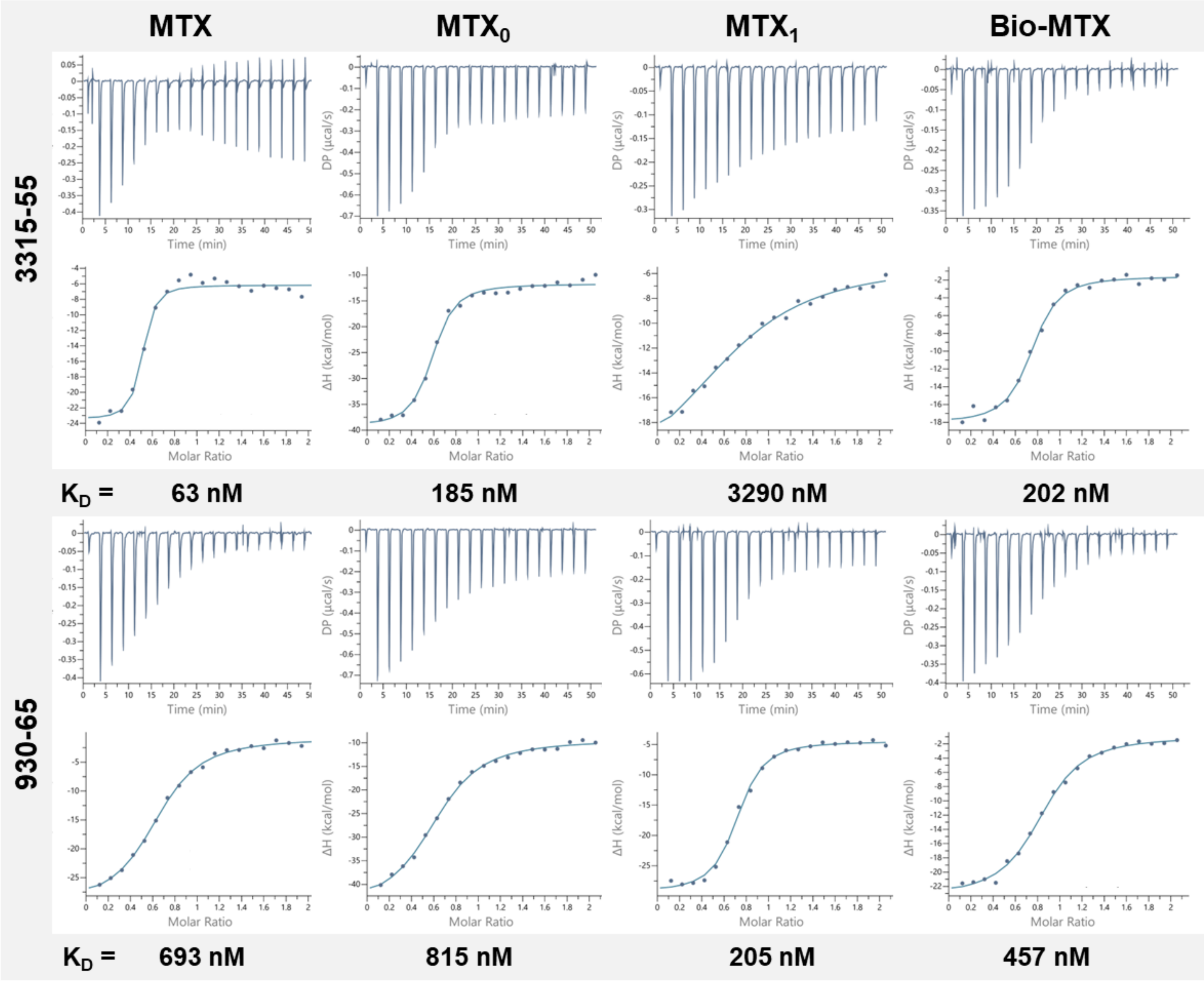

**Figure S20. Binding specificity of truncated MTX aptamer variants.** Representative ITC raw thermograms and corresponding integrated binding isotherms for 3315-55 and 930-65 measured against MTX and structurally related compounds, including MTX0, MTX1, and Bio-MTX. Experiments were performed at 20 °C in SB-K buffer [SB with 5 mM KCl], with 100 µM compound in the syringe and 10 µM aptamer in the cell. DMSO concentration was matched between syringe and cell solutions within the range of 0.1–1%. $K_D$ values derived from isotherm fitting are shown below each dataset and reported as $K_D$ ± fitting error. ITC isotherms for folic acid, MABA, and OTA are not shown because no detectable binding was observed under the tested conditions.

**Figure S21. BLI characterization of MTX binding to selected aptamers.**

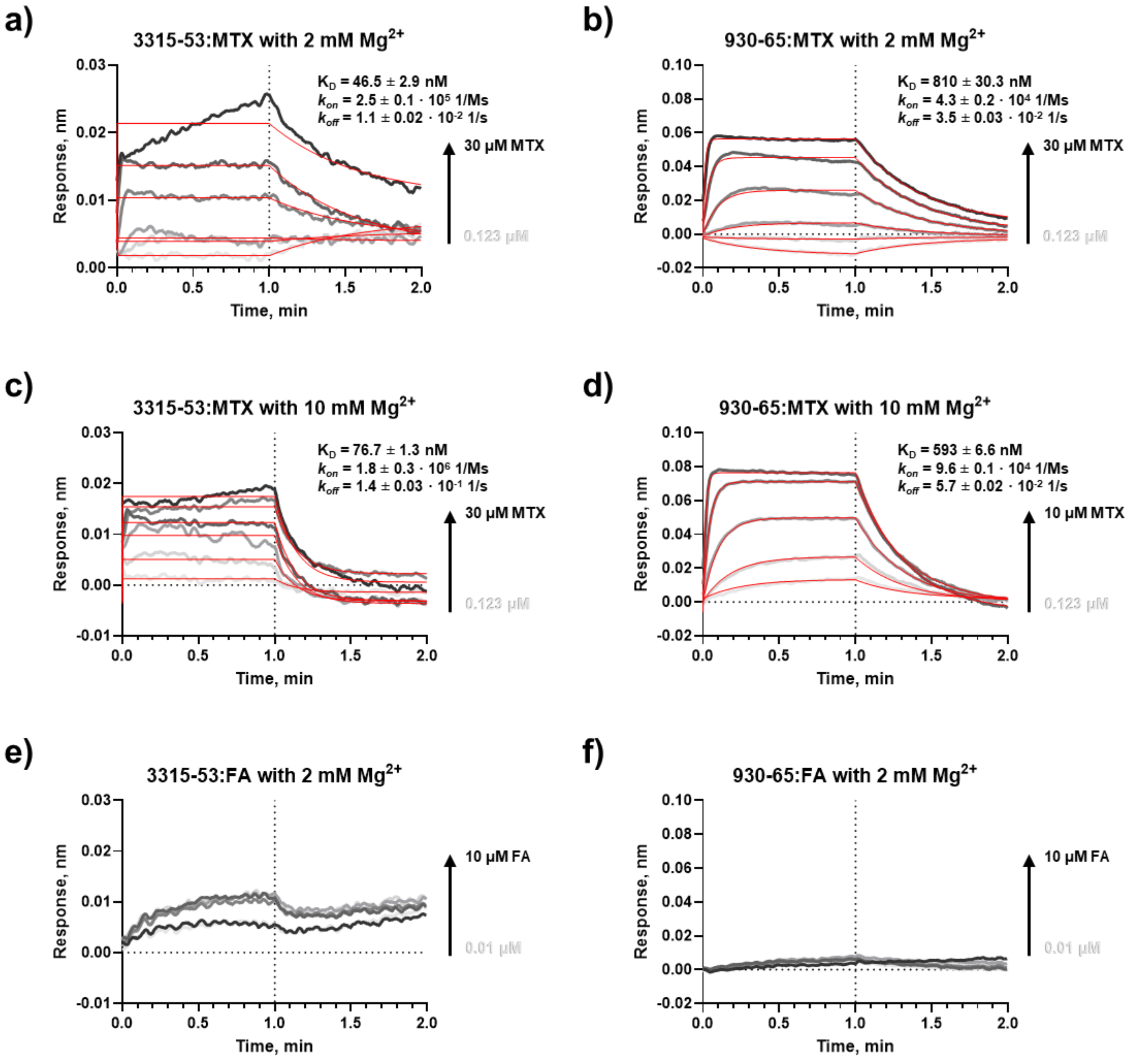

**Figure S21. BLI characterization of MTX binding to selected aptamers.** Representative biolayer interferometry (BLI) sensorgrams showing concentration-dependent binding of MTX (a–d) and folic acid (FA; e–f) to the optimized 3315-53 (a,c,e) and 930-65 (b,d,f) aptamer variants. Analysis was performed in standard selection buffer (SB; a,b,e,f) or in SB supplemented with 5 mM KCl and 10 mM MgCl2 (c,d). Experimental traces are shown in gray, and global kinetic fits in red. Association and dissociation phases were monitored across a serial MTX/FA concentration range. The sensorgrams show reproducible, concentration-dependent responses and rapid association and dissociation kinetics characteristic of the aptamer–MTX interactions, but not of aptamer–folic acid interactions, which produced only minimal responses. These results confirm that the binding signals observed for the selected aptamers were not attributable to substantial nonspecific interactions with the sensor surface or with folic acid. Kinetic parameters obtained from the global fits are indicated in each panel.

**Figure S22. Circular dichroism (CD) analysis of MTX-specific aptamers.**

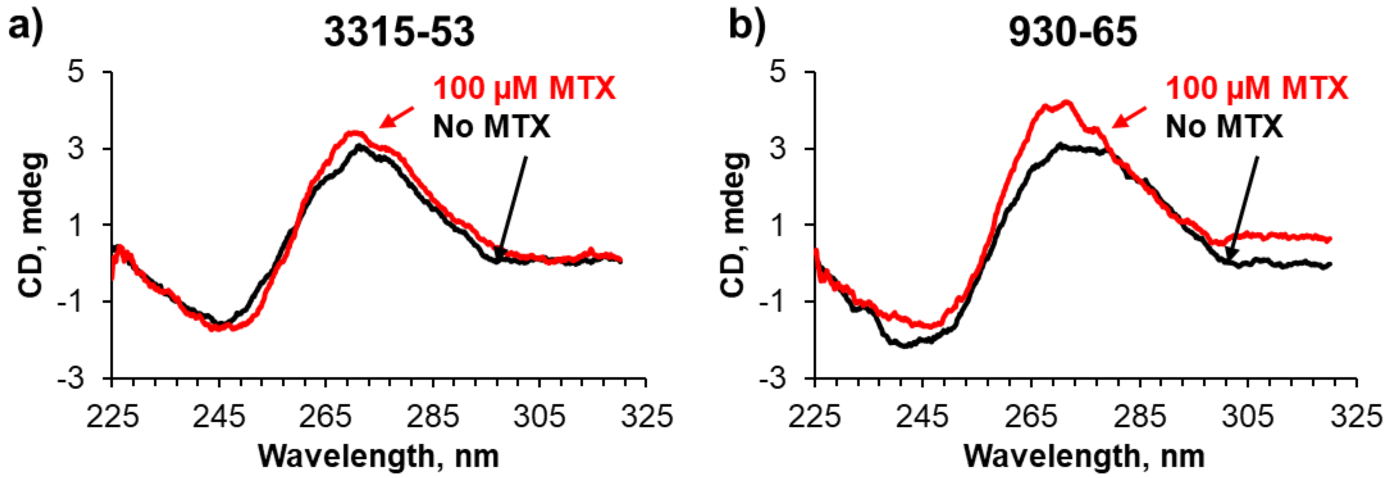

**Figure S22. Circular dichroism (CD) analysis of MTX-specific aptamers.** CD spectra of the two selected MTX aptamers recorded in the absence and presence of 100 µM MTX. Addition of MTX produced changes in the CD profiles, including altered intensity and shape of the major positive band, consistent with target-induced changes in aptamer secondary structure and/or conformational rearrangement upon MTX binding.

**Figure S23. Serum stability analysis of MTX-specific aptamer variants.**

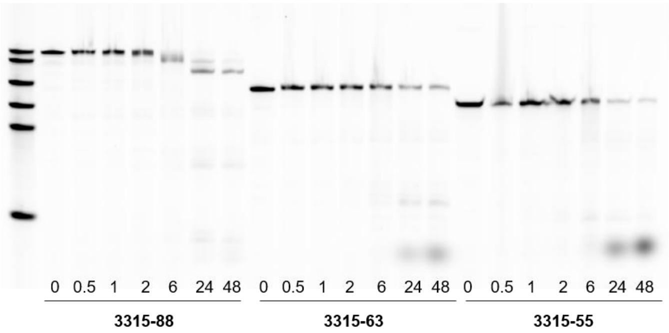

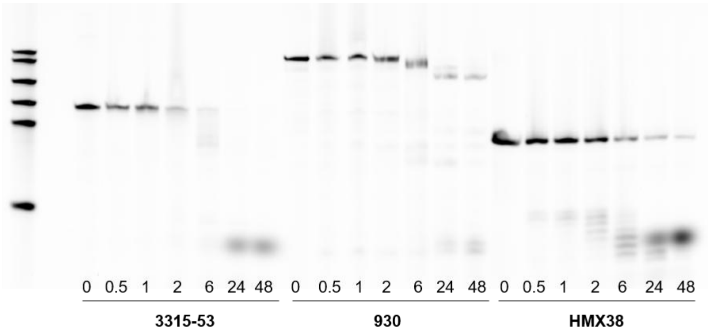

| Time, h | 0 | 0.5 | 1 | 2 | 6 | 24 | 48 |
|---|---|---|---|---|---|---|---|
| 3315-88 | 100% | 77% | 83% | 79% | 51% | 17% | 4% |
| 3315-63 | 100% | 54% | 48% | 46% | 38% | 18% | 15% |
| 3315-55 | 100% | 43% | 79% | 60% | 37% | 10% | 5% |
| 3315-53 | 100% | 45% | 52% | 19% | 7% | 1% | 0% |
| 930 | 100% | 54% | 47% | 71% | 54% | 7% | 4% |
| HMX38 | 100% | 65% | 68% | 50% | 22% | 10% | 6% |

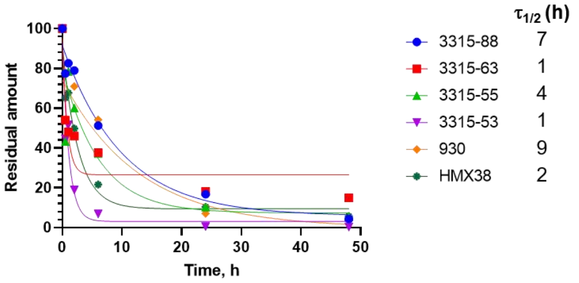

**Figure S23. Serum stability analysis of MTX-specific aptamer variants.** Aptamer variants 3315-88, 3315-63, 3315-55, 3315-53, 930-88, and HMX38 were incubated in 95% human serum at 37 °C to assess nuclease resistance under near-physiological matrix conditions. Aliquots were collected after 0, 0.5, 1, 2, 6, 24, and 48 h and analyzed by denaturing PAGE. (a) Representative gel images show time-dependent degradation of each aptamer, with intact aptamer bands progressively decreasing over the incubation period and lower-molecular-weight degradation products appearing at later time points. Band intensities for full-length aptamers were quantified and normalized to the 0 h time point, set to 100%. (b) The heat map and (c) degradation curves summarize the fraction of intact aptamer remaining over time, revealing variant-dependent serum stability. Among the tested sequences, truncated 3315 variants retained a measurable intact aptamer after prolonged incubation, whereas some variants showed rapid degradation within the first hours. This analysis demonstrates that aptamer truncation can influence serum stability and supports the selection of compact 3315 variants for further biosensor development in biological matrices.

**Figure S24. Mutational analysis of the MTX-binding aptamer 3315.**

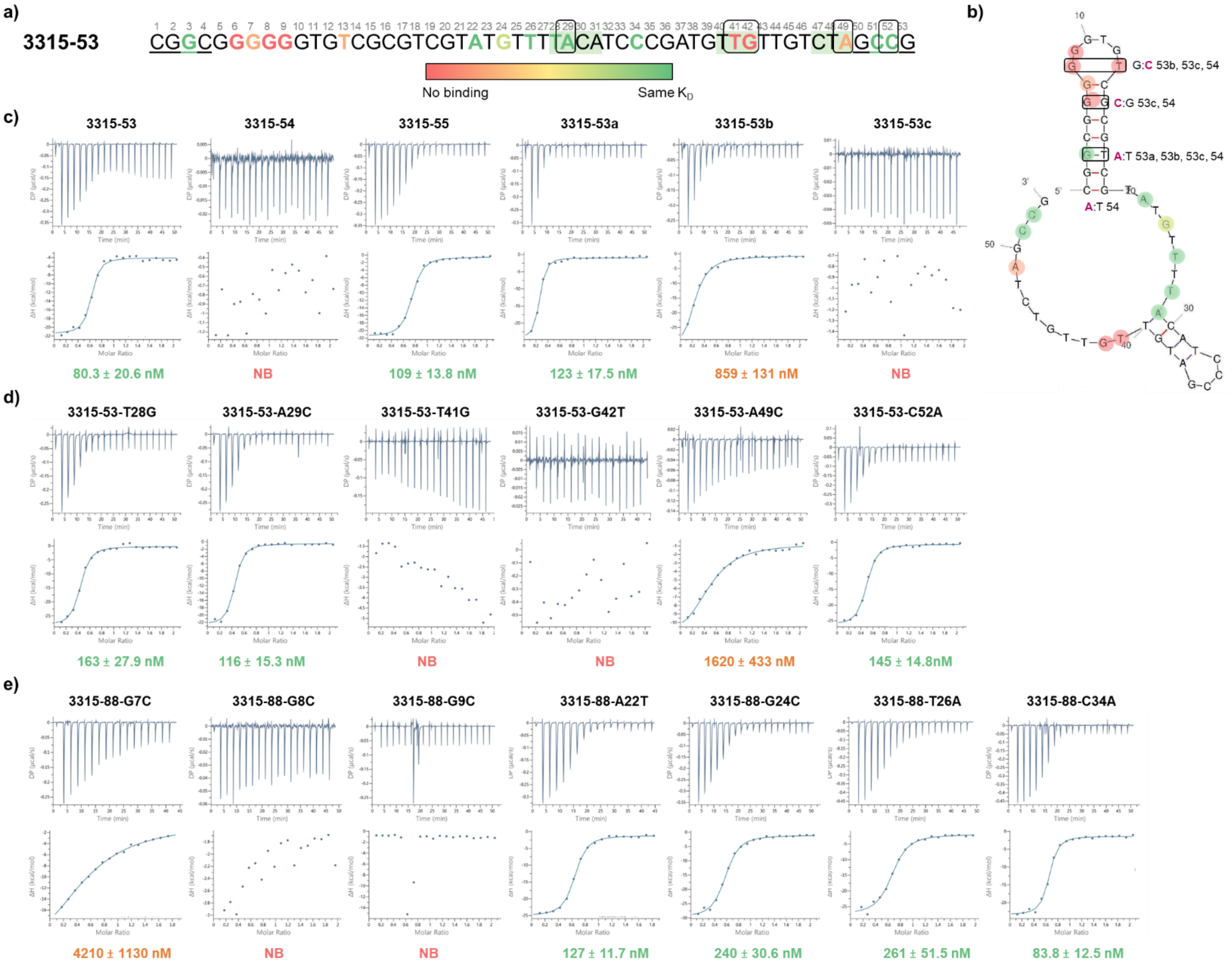

**Figure S24. Mutational analysis of the MTX-binding aptamer 3315.** (a) Sequence map of 3315-53 showing the positions of residues investigated by mutagenesis. Nucleotides predicted by MD/MM-GBSA analysis to contribute to MTX recognition are highlighted, with the color scale indicating the experimentally observed effect of mutation on MTX binding, from preserved binding (green) to reduced affinity (orange) or no detectable binding (red). (b) Mfold-predicted secondary structure of 3315-53 showing the corresponding mutated residues and binding effects. (c–e) Representative ITC thermograms and integrated binding isotherms for variants designed to evaluate the contribution of stem architecture (c), predicted MTX-binding-site residues in 3315-53 (d), and corresponding mutations in the full-length 3315-88 aptamer (e). Apparent $K_D$ values are indicated below each binding isotherm. Mutations at T41 and G42 abolished detectable MTX binding, whereas A49C substantially reduced affinity; substitutions at A29 and C52 were better tolerated. Alterations designed to extend or stabilize the stem region generally did not improve affinity and, in several cases, disrupt detectable binding, indicating that both the predicted binding-site residues and the native local stem-loop architecture contribute to formation of the MTX-binding conformation. NB: no detectable binding under the tested conditions.

**Figure S25. ITC-based mutational analysis of the previously reported MTX-binding aptamer HMX38.**

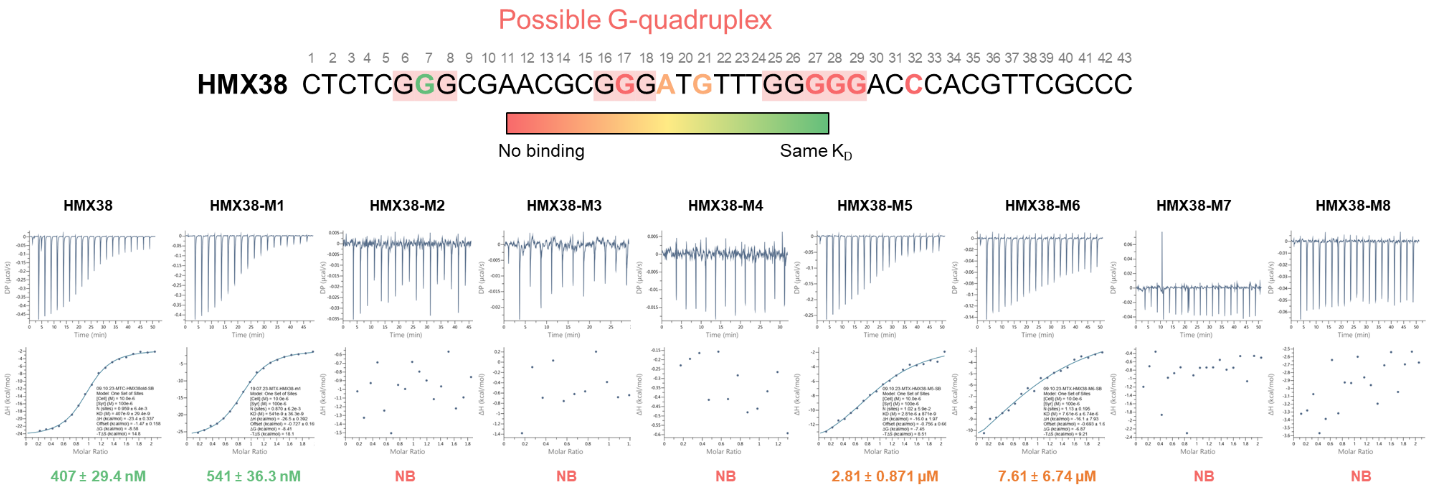

**Figure S25. ITC-based mutational analysis of the previously reported MTX-binding aptamer HMX38.** Single-point mutations were introduced into the predicted G-rich/G-quadruplex-forming regions of HMX38 and selected flanking positions to evaluate their contribution to MTX recognition. The sequence map indicates the position of each mutation, with residues colored by the mutation's effect on apparent MTX-binding affinity. Representative ITC thermograms and integrated binding isotherms are shown for HMX38 and its mutant variants.

**Figure S26. Mutational validation of predicted MTX-binding pockets in aptamer 930-65.**

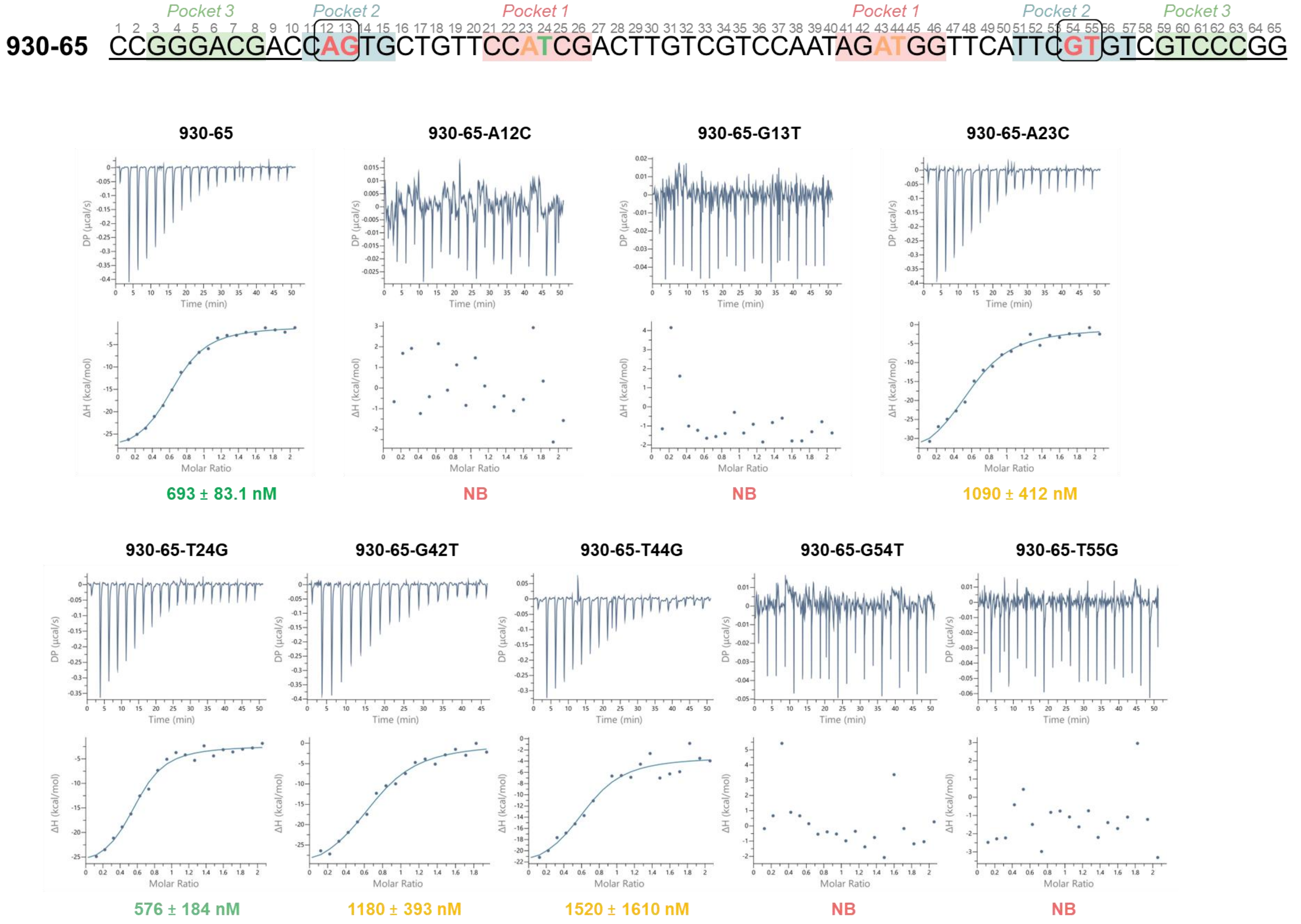

**Figure S26. Mutational validation of predicted MTX-binding pockets in aptamer 930-65.** ITC analysis was performed to evaluate the contribution of nucleotides located within the predicted MTX-binding pockets of 930-65. The sequence map shows the positions of the three candidate binding pockets identified by docking and MD simulations, with mutated residues highlighted according to their effect on MTX binding affinity.

**Figure S27. ITC validation of scrambled aptamer variants.**

**a)**

| | |
|---|---|
| **3315-53** | CGGCGGGGGGTGTCGCGTCGTATGTTTTACATCCCGATGTTGTTGTCTAGCCG |
| **3315-53-SCR** | TACATAAGCCGGGTGAGGCTCCTTCGTTTTCGCTTCCGCGGTTGTGGGGTGTG |
| **930-65** | CCGGGACGACCAGTGCTGTTCCATCGACTTGTCGTCCAATAGATGGTTCATTCGTGTCGTCCCGG |
| **930-65-SCR** | ACTCGCCTGTGTACACGCCGCGGTGTTATCTGGTCTTATGACGTTACGTGCTCCGACTACGAG |

**b)**

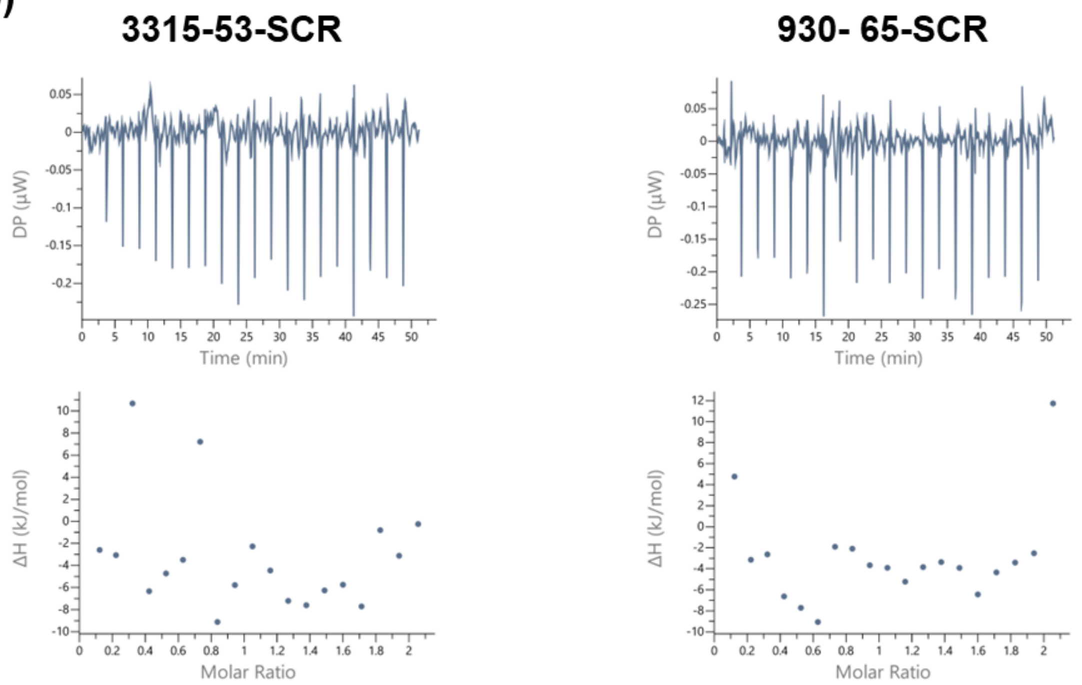

**Figure S27. ITC validation of scrambled aptamer variants.** a) Scrambled control sequences derived from the active 3315-53 and 930-65 aptamers were analyzed by ITC to evaluate whether MTX binding depends on the specific aptamer sequence and folded architecture. Sequence maps show the scrambled 3315-53 and 930-65 variants, with nucleotides color-coded by base identity. b) Representative ITC raw thermograms and integrated heat plots are shown for MTX titration into the scrambled aptamer variants. Experiments were performed at 20 °C under standard ITC conditions, with 100 μM MTX in the syringe and 10 μM aptamer in the cell. No detectable binding was observed for either scrambled sequence, confirming that MTX recognition by the parental aptamers is sequence- and structure-dependent rather than arising from nonspecific interaction with DNA.

**Figure S28. Electrochemical characterization of MTX EAB sensor.**

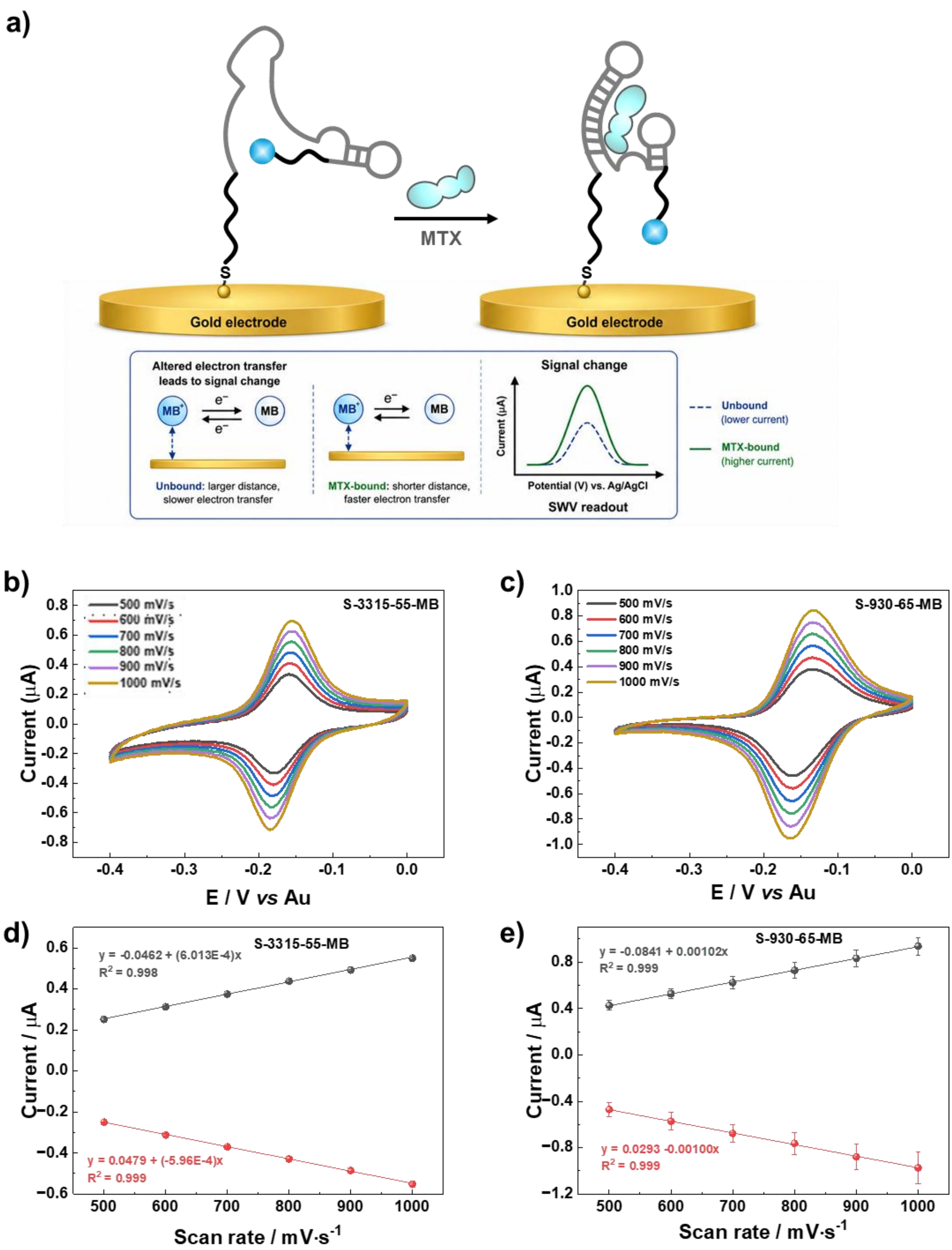

**Figure S28. Electrochemical characterization of MTX EAB sensor constructs by scan-rate-dependent cyclic voltammetry.** (a) Schematic representation of the electrochemical aptamer-based (EAB) MTX sensor architecture, consisting of a thiolated DNA aptamer immobilized on a gold electrode and labeled with a methylene blue redox reporter (MB). MTX binding induces an aptamer conformational change that alters electron transfer between methylene blue and the electrode surface, generating a measurable signal change. Representative cyclic voltammograms of (b) S-3315-55-MB and (c) S-930-65-MB recorded in PBS at scan rates of 500, 600, 700, 800, 900, and 1000 mV $s^{-1}$. Corresponding plots of anodic and cathodic peak current as a function of scan rate for (d) S-3315-55-MB and (e) S-930-65-MB. The linear dependence of peak current on scan rate indicates that electron transfer is dominated by a surface-confined redox process, consistent with immobilization of MB-labeled aptamers on the gold electrode surface. Aptamer immobilization concentration: 0.5 μM.

**Figure S29. Square wave voltammograms at 600 Hz of the 3315-55 and 930-65 aptamers.**

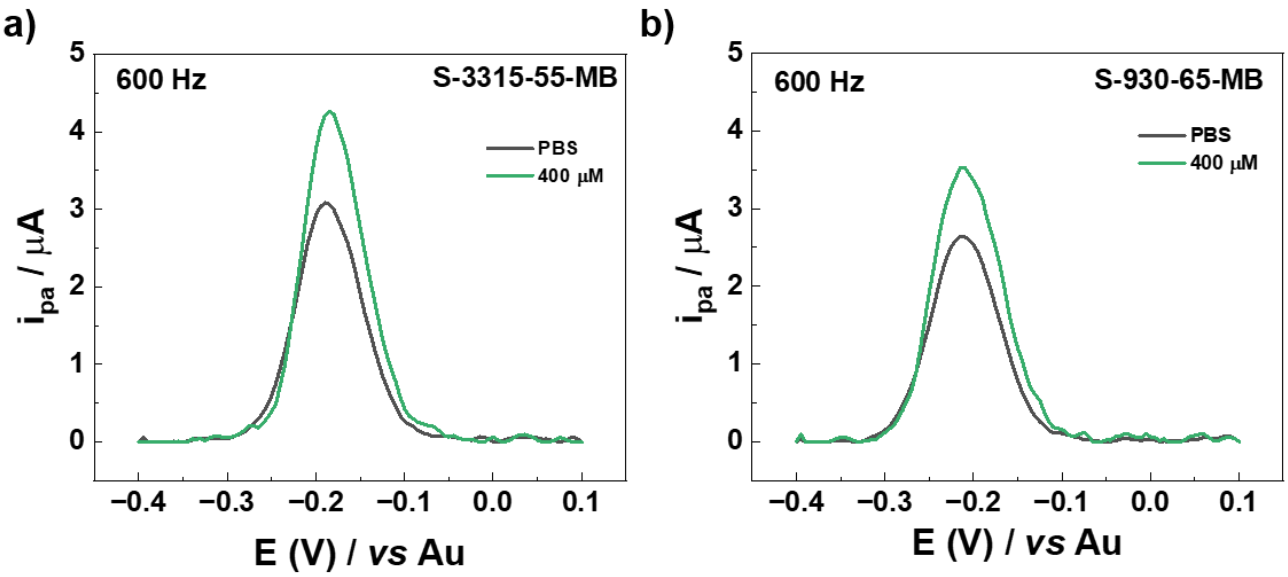

**Figure S29. Square wave voltammograms (SWV) at 600 Hz for the 3315-55 and 930-65 aptamers.** a) 3315-55 and b) 930-65, in PBS (black line) and in the presence of 400 µM MTX. MTX solutions were prepared in PBS, and 100 µL was added to the electrode surface and incubated for 10 min. The measurements were recorded immediately afterward.

**Figure S30. MTX calibration curves for electrochemical aptamer-based sensors 3315-55 and 930-65.**

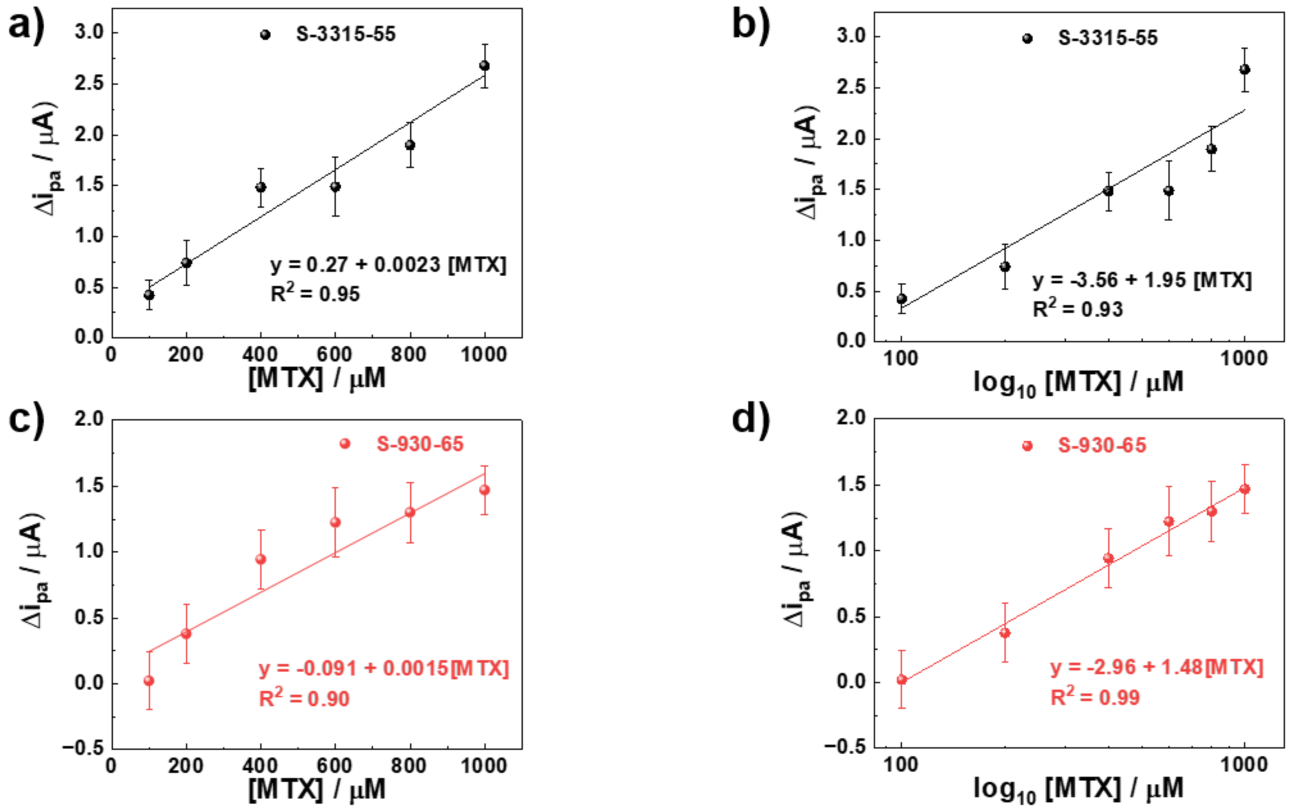

**Figure S30. MTX calibration curves for electrochemical aptamer-based sensors 3315-55 and 930-65.** Calibration curves were obtained using MB-labeled aptamers 3315-55 and 930-65 immobilized on gold electrodes. MTX solutions were prepared in PBS at concentrations of 100, 200, 400, 600, 800, and 1000 μM. For each measurement, 100 μL of MTX solution was added to the electrode surface and incubated for 10 min prior to electrochemical analysis. Calibration responses are shown for 3315-55 as (a) $\Delta i_{pa}$ versus MTX concentration and (b) $\Delta i_{pa}$ versus $\log_{10}$[MTX], and for 930-65 as (c) $\Delta i_{pa}$ versus MTX concentration and (d) $\Delta i_{pa}$ versus $\log_{10}$[MTX]. $\Delta i_{pa}$ is $i_{pa,analyte} - i_{pa,PBS}$. Data represent mean ± SD from 3 independent electrodes. Linear regression equations and $R^2$ values are shown on each plot.

**Figure S31. Selectivity analysis of MTX EAB sensors against folic acid.**

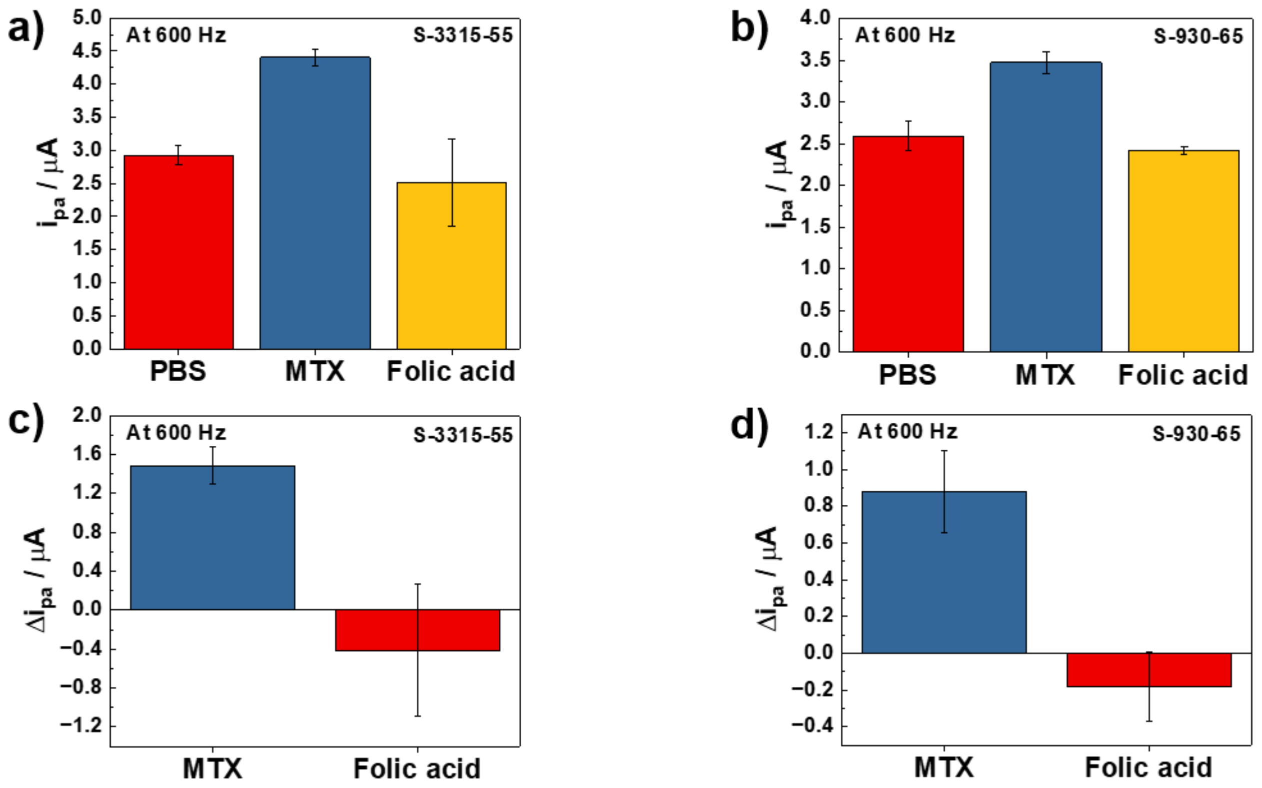

**Figure S31. Selectivity analysis of MTX EAB sensors against folic acid.** Selectivity of the electrochemical aptamer-based sensors was evaluated using square-wave voltammetry at 600 Hz. Peak anodic currents ($i_{pa}$) were extracted for (a) 3315-55 and (b) 930-65 in PBS alone, in the presence of MTX as the target analyte, and in the presence of folic acid as a structurally related off-target control. MTX and folic acid were tested at 400 μM. The corresponding current changes, calculated as $\Delta i_{pa} = i_{pa,analyte} - i_{pa,PBS}$, are shown for (c) 3315-55 and (d) 930-65. Both sensors showed a stronger electrochemical response to MTX than to folic acid, supporting selective target recognition under the tested conditions. Data represent mean ± SD from independent electrodes.

(15) Case, D. A.; Aktulga, H. M.; Belfon, K.; Cerutti, D. S.; Cisneros, G. A.; Cruzeiro, V. W. D.; Forouzesh, N.; Giese, T. J.; Götz, A. W.; Gohlke, H.; Izadi, S.; Kasavajhala, K.; Kaymak, M. C.; King, E.; Kurtzman, T.; Lee, T. S.; Li, P.; Liu, J.; Luchko, T.; Luo, R.; Manathunga, M.; Machado, M. R.; Nguyen, H. M.; O'Hearn,

K. A.; Onufriev, A. V.; Pan, F.; Pantano, S.; Qi, R.; Rahnamoun, A.; Risheh, A.; Schott-Verdugo, S.; Shajan, A.; Swails, J.; Wang, J.; Wei, H.; Wu, X.; Wu, Y.; Zhang, S.; Zhao, S.; Zhu, Q.; Cheatham, T. E.; Roe, D. R.; Roitberg, A.; Simmerling, C.; York, D. M.; Nagan, M. C.; Merz, K. M. AmberTools. *J. Chem. Inf. Model.* **2023**, *63* (20), 6183–6191. https://doi.org/10.1021/acs.jcim.3c01153.

(38) Moghbel, A.; Zand-Moghaddam, A.; Rezaee, S.; Pedram, M. *High-Performance Liquid Chromatography Determination of Methotrexate in Plasma*; 2003.
